# Petrosal morphology and cochlear function in Mesozoic stem therians

**DOI:** 10.1101/490367

**Authors:** Tony Harper, Guillermo Rougier

## Abstract

Here we describe the bony anatomy of the inner ear and surrounding structures seen in three of the most plesiomorphic crown mammalian petrosal specimens in the fossil record. Our study sample includes the stem therian taxa *Priacodon fruitaensis* from the Upper Jurassic of North America, and two isolated petrosal specimens colloquially known as the Höövör petrosals, recovered from Aptian-Albian sediments in Mongolia. The second Höövör petrosal is here described at length for the first time. All three of these stem therian petrosals and a comparative sample of extant mammalian taxa have been imaged using micro-CT, allowing for detailed anatomical descriptions of osteological correlates of functionally significant neurovascular features, especially along the abneural wall of the cochlear canal.

The high resolution imaging provided here clarifies several hypotheses regarding the mosaic evolution of features of the cochlear endocast in early mammals. In particular, these images demonstrate that the membranous cochlear duct adhered to the bony cochlear canal abneurally to a secondary bony lamina before the appearance of an opposing primary bony lamina or tractus foraminosus. Additionally, while corroborating the general trend of reduction of venous sinuses and plexuses within the pars cochlearis seen in crownward mammaliaformes generally, the Höövör petrosals show the localized enlargement of a portion of the intrapetrosal venous plexus. This new excavation is for the vein of cochlear aqueduct, a structure that is solely or predominantly responsible for the venous drainage of the cochlear apparatus in extant therians. However, given that these stem therian inner ears appear to have very limited high-frequency capabilities, the development of these modern vascular features the cochlear endocast suggest that neither the initiation or enlargement of the stria vascularis (a unique mammalian organ) is originally associated with the capacity for high-frequency hearing or precise sound-source localization.

## Introduction

Therian mammals today display an impressive variety of auditory features facilitating the adept detection of airborne sound, making this group arguably the most acoustically sophisticated and diverse living or extinct clade of terrestrial vertebrates. The widespread capacity for high-acuity hearing (in terms of sensitivity, specificity, and highest detectable frequency) among the majority of extant therian mammals has reinforced the assumption among neontologists that the Mesozoic members of the therian stem linage were stereotypically nocturnal forms that leveraged their auditory faculties to locate small prey and escape gigantic predators (i.e. [1]). However, this supposition has not been supported by the known fossil record of stem therians, with prior descriptions [2-7] demonstrating that these forms lacked the level of petrosal organization characterizing plesiomorphic marsupials [8], afrotheres [9], eulipotyphlans, and primatomorphs [10]. Additionally, these reports have relied heavily on the description of the external morphology of the ear region, and for the most part remained silent or equivocal regarding the performance and/or physiological implications of the bony anatomy seen in fossil specimens.

Conversely, several biomechanical/experimental studies on auditory anatomy and physiology across extant tetrapods [11-15] highlight the unique nature of the therian cochlear apparatus (with its well-ordered acoustic hair-cell populations arrayed along the organ of Corti, high endolymphatic potential, and absence of the lagenar macula), as well as the plesiomorphic nature of the monotreme cochlea with respect to many modern and fossil therians [16, 17]. The complete loss of the lagenar macula in particular has been posited as a therian-lineage evolutionary breakthrough that allowed for the later elongation and sophistication of the cochlear apparatus for non-linear amplification of high-frequency stimuli [18]. Conversely, the retention of a functional lagenar macula, along with its accompanying otoconial and neurovascular structures, in the monotreme and sauropsid lineagesis hard to conciliate with the soft tissue adaptations seen in modern therians. These “therian” features include: 1) the exclusive reliance on the stria vascularis as the major endolymph producing organ, 2) the well-developed electromotility of prestins and other molecular components of the “cochlear amplifier”, and 3) the radical elongation and geometrical reorganization of the cochlear sensory epithelium. The distribution of these characteristics in extant animals, within the wider framework of cynodont evolution, points to a fundamental transformation of the cochlear apparatus somewhere within the phylogenetic vicinity of the therian stem [18].

The lack of anatomical depiction within the Mesozoic fossil record is understandable given the general lack of high-fidelity bony correlates for key soft tissue structures such as the cochlear duct, lagenar macula, and stria vascularis. Additionally, prior studies using high-resolution imaging of the internal anatomy in pertinent fossils have not commented on or depicted the possible existence of several physiologically significant bony structures that are not familiar to most paleontologists. This report uses high-resolution micro-CT information to update and qualify the descriptions of petrosal anatomy provided in the representative sample of stem therians focused on by [2] and [3], two large-scale studies characterizing the fossil record of early mammalian petrosal evolution (Fig 1). These images and reconstructions are framed within a broader comparative and functional setting of modern mammalian auditory physiology, and morphology with the hope of bridging both fields which, due to the steep learning curves involved, have had only limited cross-referencing. These new high-resolution images allow the first descriptions of the labyrinthine anatomy in some of the most morphologically plesiomorphic petrosal regions known in the crown mammalian fossil record. The three specimens focused on here include the relatively well-known triconodontid *Priacodon fruitaensis* and the two isolated petrosal specimens known as the Höövör petrosals. The second Höövör petrosal (Fig 1 a, b) has not been described or figured, and has only been cursorily referred to in previous publications (i.e. [2]). We regard this second petrosal as distinct from that of the previously described petrosal in [2], and therefore provide here a complete description and assessment of this specimen.

**Fig 1.**
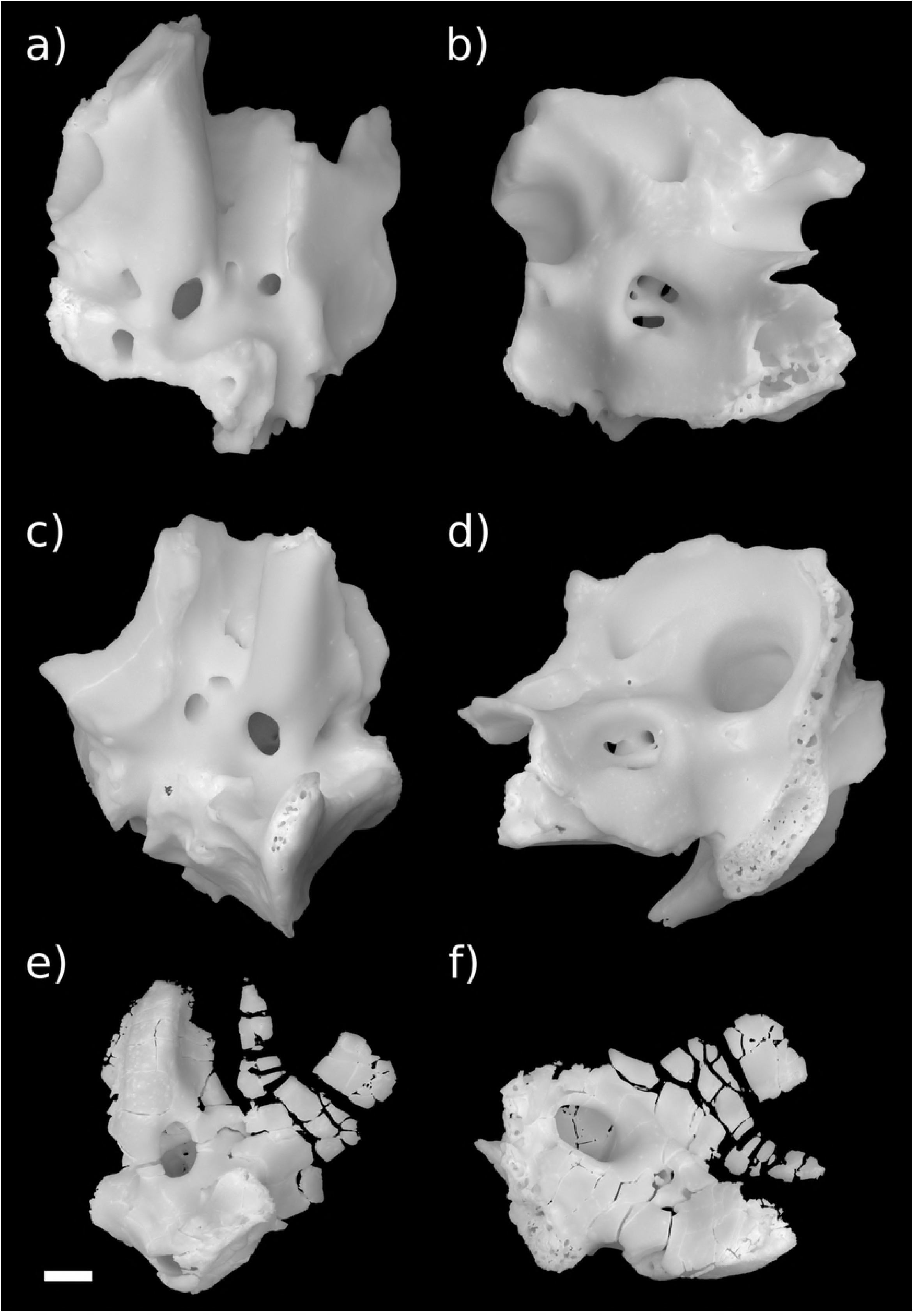
Stem therian petrosal specimens described in this report. a,b Höövör Petrosal 2 (PSS-MAE-119); c,d, Höövör Petrosal 1 (PSS-MAE-104); e,f left petrosal of *Priacodon fruitaensis* (LACM 120451). a,c,e in ventral view; b,d,f in dorsomedial view. Scale bar is 1 mm.

The novel information presented here does little to overturn previous taxonomic attributions of these stem therians especially the first and second Höövör petrosals, one or both of which are tentatively referred to taxa within Gobiconodontidae and/or Tinodontidae. However, the new anatomical representation provided by the images used here demonstrates the presence of a combination of plesiomorphic and derived character states of the cochlear canal that support their ancestry with crown therian and cladotherian mammals. In particular the stem therian endocasts presented here show the presence of a secondary lamina, and in the Höövör petrosals, the earliest reported appearance of the vein of the cochlear aqueduct (the main venous drainage for the cochlear apparatus in therian mammals today). For reasons outlined in the discussion section, these osteological features are most likely associated with the elaboration of the macromechanical form of tuning, a unique cochlear functionality that provides the majority of modern therian mammals ultrasonic frequency sensitivity. However, because of the plesiomorphic geometry of the cochlear canal seen in these specimens, the ability to detect mid-range to high frequencies was at most only incipiently developed in these earliest members of the stem therian lineage. The osteological evidence provided here therefore suggests that many of the unique features of therian cochlear histology and physiology evolved gradually after their split from the lineage leading to monotremes, and that these features originally functioned in service of highly sensitive and selective low-frequency hearing.

## Materials and methods

Osteological nomenclature used in the following descriptions of petrosal morphology are taken primarily from [19-21]. Terms specific to the description of labyrinthine endocasts are also taken from [22,23] for therian mammals; and [24] and [17] for non-mammals and monotremes, respectively. Nomenclature for cranial vasculature is taken from [25-28].

Anatomical abbreviations used in figures and text:

ac: aqueductus cochleae (for perilymphatic duct)
acf: aperture of cochlear fossula (external to fenestra cochleae)
adm: arteria diploëtica magna
al: anterior lamina
amp-a: anterior semicircular canal ampulla
amp-h: horizontal semicircular canal ampulla
amp-p: posterior semicircular canal ampulla
av: aqueductus vestibuli (for endolymphatic duct)
bpsal: border of periosteal surface of alnterior lamina
bs: basisphenoid sutural surface
cas: cavum supracochleare
cc-p: primary common crus
cc-s: secondary common crus
cdh: centripetal diverticulum of horizontal semicircular canal
ce: cavum epiptericum
cf: cochlear fossula
ci: crista interfenestralis
ct: crista transversa (crista falciformis in [17])
cVIII: cochlear branch of vestibulocochlear nerve
cvp: circumpromontorial venous plexus
coc: cochlear canal
dag: dorsal ascending groove
eapc: endocranial aperture of prootic canal
ect: ectotympanic bone
eps-a: anterior epipromontorial sinus (“trans-promontorial sinus a” in [29])
eps-p: posterior epipromontorial sinus (“trans-promontorial sinus p” in [29])
fac: facial canal (aqueductus Fallopii in [30])
fc: fenestra cochleae (for secondary tympanic membrane)
fo: fenestra ovalis
gpn: greater petrosal nerve
hF: hiatus Fallopii (for greater petrosal nerve)
hfn: hyomandibular branch of facial nerve
hps: hypopromontorial sinus
hs: half-pipe shaped sulcus
iam: internal acoustic meatus
ips: inferior petrosal sinus
ijv: internal jugular vein
ir: inferior ramus of stapedial artery
jn: jugular notch
lf: lateral flange
lhv: lateral head vein
li: lagenar inflation
mm: manubrium of malleus
ntr-a: notch for temporal ramus a
ntr-b: notch for temporal ramus b
pcs: sinus around prootic canal
pf: perilymphatic foramen
pfc: prefacial commissure (suprafacial commisure)
pop: paroccipital process
pos: paroccipital sinus
pov: prootic vein
pptc: petrosal contribution to post-temporal canal
?ptv: possible location of post-trigeminal vein
re: recessus ellipticus (for utricle)
rpm: rostral (anterior) process of malleus (= goniale/ prearticular)
rs: recessus sphericus (for saccule)
rso: ramus supraorbitalis
ssc-a: anterior semicircular canal
sa: stapedial artery
saf: subarcuate fossa
sas: subarcuate venous sinus
sl: secondary (abneural) bony lamina of cochlear canal
sf: stapedius fossa
sff: secondary facial foramen
sr: superior ramus of stapedial artery
tapc: tympanic aperture of prootic canal
tf: tractus foraminosus
tph: tympanohyal
tr-a: anterior temporal ramus
tr-b: posterior temporal ramus
uaf: foramen for utriculoampullar branch of vestibular nerve)
vag: ventral ascending groove
VII: facial nerve
vVIII: vestibular branch of vestibulocochlear nerve
vcaq: bony canal accommodating vein of the cochlear aqueduct (Canal of Cotugno)

Other abbreviations taken from [31], and [15]:

H1: Höövör petrosal specimen 1 (PSS-MAE-104)
H2: Höövör petrosal specimen 2 (PSS-MAE-129)
IHC: inner hair cell
OHC: outer hair cell
SC: cochlear space constant (in octaves/ mm)
ILD: interaural level difference
ITD: interaural time difference
FHS: functional head size (in microseconds)

The Aptian or Albian Höövör locality (also transliterated Khoobur, Khobur, Khoboor, Khovboor, with varying diacritical marks) in Guchin-Us District, Mongolia is an important source of plesiomorphic, derived, and autapomorphous mammalian specimens from the relatively under-sampled “middle Cretaceous” (Aptian-Albian). Originally discovered by joint Soviet-Mongolian expeditions [32] this locality has been worked during subsequent field seasons by both the Mongolian Academy of Sciences and American Museum of Natural History [33]. Most significantly from the perspective of this report are the three isolated petrosal specimens that this locality has produced, an early stem therian termed “Höövör petrosal 1” (Fig 1 c, d, PSS-MAE-104; [2]), a crown therian petrosal referred to the eutherian taxon *Prokennalestes trofimovi* (PSS-MAE-106; [34]), and an undescribed but previously mentioned specimen termed “Höövör petrosal 2” (Fig 1 a, b, PSS-MAE-129) that is described below.

One of the most plesiomorphic crown mammals known from petrosal remains is *Priacodon fruitaensis* (Fig 1 e, f; [3]). The specific epithet for this taxon is named for the Upper Jurassic (Kimmeridgian) microfossil locality at Fruita Paleontological Area, Mesa County, Colorado [35]. The specimen of *Priacodon fruitaensis* figured and described here (LACM 120451) is the left periotic region figured in [3], that was found in association with a fragmentary left exoccipital and other cranial elements. To facilitate the following discussion, unless otherwise indicated, the generic term *Priacodon* will refer specifically only to *Priacodon fruitaensis.* Also, the contractions H1 and H2 will be used to signify PSS-MAE-104 and PSS-MAE-129, respectively.

Additionally, because this report relies on the precise delimitation of small and poorly understood features of the otic capsule, the use of several nontraditional terminological conventions is necessitated in the following text. Probably the most unconventional of these is the use of the term “endolymphatic labyrinth” to refer to the inner ear structure commonly referred to as the “membranous labyrinth”; and concomitantly the use of “perilymphatic labyrinth” as a substitute for “bony labyrinth” or “osseous labyrinth”. These terms facilitate the the description of several of the more obscure features of the mammalian periotic region, most importantly the perilymphatic duct, that is variable in terms of its enclosure within an accommodating bony canal among modern and extinct mammals. Additionally, while the use of the word “duct” in the term perilymphatic duct is an allusion to its membranous composition (while the word “canal” refers to a bony tubular structure), the membranous perilymphatic duct is itself a component of the greater perilymphatic labyrinth (i.e. the “bony” labyrinth). Because the distinction between the perilymph-and endolymph-filled spaces is the most important attribute to consider in descriptions of the evolutionary transformation of the inner ear, the terms “endolymphatic labyrinth” and “perilimphatic labyrinth” are used instead of terms referring to the connective tissue type enclosing these spaces.

As mentioned in [36], the description of isolated petrosal fossils from taxa insufficiently known from cranial remains often precludes the use of precise global anatomical directional terms. This report attempts to describe petrosal remains using a “local” directional system, whereby “anterior” or “rostral” is used to signify the authors’ best estimate of the direction corresponding to the anterior direction in a complete skull. In the petrosals described here, “anterior” therefore signifies the direction toward the lateral side of the tip of the petrosal promontorium furthest away from the fenestra ovalis, and toward the presumed location of the alisphenoid and entopterygoid bones. The term “posterior” or “caudal” is therefore used to refer to the opposite direction; whereas “medial” and “lateral” are used to refer to the directions perpendicular to the anteroposterior axis, toward the internal and external surface of the skull, respectively. Since the dorsoventral axis is much less ambiguous in these remains, no special definition is needed. These directional terms are chosen to reflect our presumption that the long axis of the promontorium is oriented approximately 45° towards the midsagittal plane, as seen in many Mesozoic mammalian petrosals that have been preserved in situ [37,38].

Finally, novel terms are used here for large-scale middle-ear character states seen in early mammals. These are “Detached Middle Ear” (DME), “Partial Middle Ear” (PME), and “Mandibular Middle Ear” (MME), referring to the adult condition of having a completely independent auditory apparatus and mandible, a primarily cartilaginous connection between the malleus and lower jaw, and a completely integrated jaw and ossicular chain, respectively. This idiosyncratic terminology is meant to substitute for the less descriptive terms used by ([39,40], inter alios) that were defined with implicit taxonomic content. These terms: “Definitive Mammalian Middle Ear” (DMME), “Partial Mammalian Middle Ear” (PMME), Transitional Mammalian Middle Ear (TMME), and “Mandibular Middle ear of Cynodonts” (MMEC), are undesirable for the following discussion because of their assumed (and likely incorrect) application to members of the clades mentioned in their name. For instance, the term Definitive Mammalian Middle Ear suggests that this character state can be shown to be apomorphic for the crown mammalian common ancestor, and therefore unites the sister lineages leading to modern monotremes and therians; however, the DMME is neither definitive (under most hypotheses of the relationships of early mammals), nor is it diagnostic of Mammalia as a clade; making it not only incorrect but pernicious. The terms DME, PME, and MME have an identical phylogenetic distribution (i.e they are extensively identical) as DMME, PMME, and MMEC (respectively), but are defined solely on observable anatomical features of the ear and jaw. This nomenclature allows the following text to more clearly differentiate the anatomical and phylogenetic composition of varying sets of synapsid taxa, especially in the discussion section below.

## Results

### External anatomy of the second Höövör petrosal

Following the heuristic presented in [37], the periotic complex (petrosal with its associated mastoid ossification; [41,42,20]) can be envisaged as a generally tetrahedral structure with four major surfaces: tympanic (Fig 2 a, b), cerebellar (Fig 2 c, d), squamosal (Fig 3 a, b) and mastoid. Because the mastoid region has not been preserved in the H2 specimen, the terms periotic and petrosal will be used synonymously in its following descriptions. The petrosal itself is composed of two synostosed components, the pars canalicularis (containing the utrical and semicircular canals) dorsoposteriorly, and the pars cochlearis (containing the saccule and cochlear duct) anteroventrally [43]. The tympanic surface is most apparent in ventral view and contains structures associated with the suspension of the ossicular chain, the incipient fenestra cochleae, and foramina supporting the distribution of neurovascular organs. The topography of this region is defined mainly by the promontorium and lateral trough anteriorly, and an expanded post-promontorial region posteriorly. The cerebellar surface of the petrosal is exposed endocranially, and has several excavations, that accommodate central and peripheral nervous structures and vasculature. These include the internal acoustic meatus anteromedially, the subarcuate fossa (only the anterior half having been preserved in H2) posteromedially, a depression for the trigeminal ganglion (=semilunar/gasserian ganglion) anterolaterally, and the endocranial aperture of the prootic canal posteriolaterally. The laterally facing squamosal surface, and the caudally facing mastoid surface (poorly preserved in H2) are both rugose because of their complex sutural interdigitations with surrounding bones (most probably the squamosal, exoccipital and basioccipital), and canals transmitting ramifications of the diploëtic and stapedial vessels. If the periotic is considered as a six-sided box, aligned with the principal anatomical axes, it would be fair to ask which structures occupy the anterior and medial (Fig 3 c, d) faces of the periotic as well. However, these margins of the petrosal are exposed cancellous bone and the intramural inferior petrosal sinus governs the topography here.

**Fig 2.**
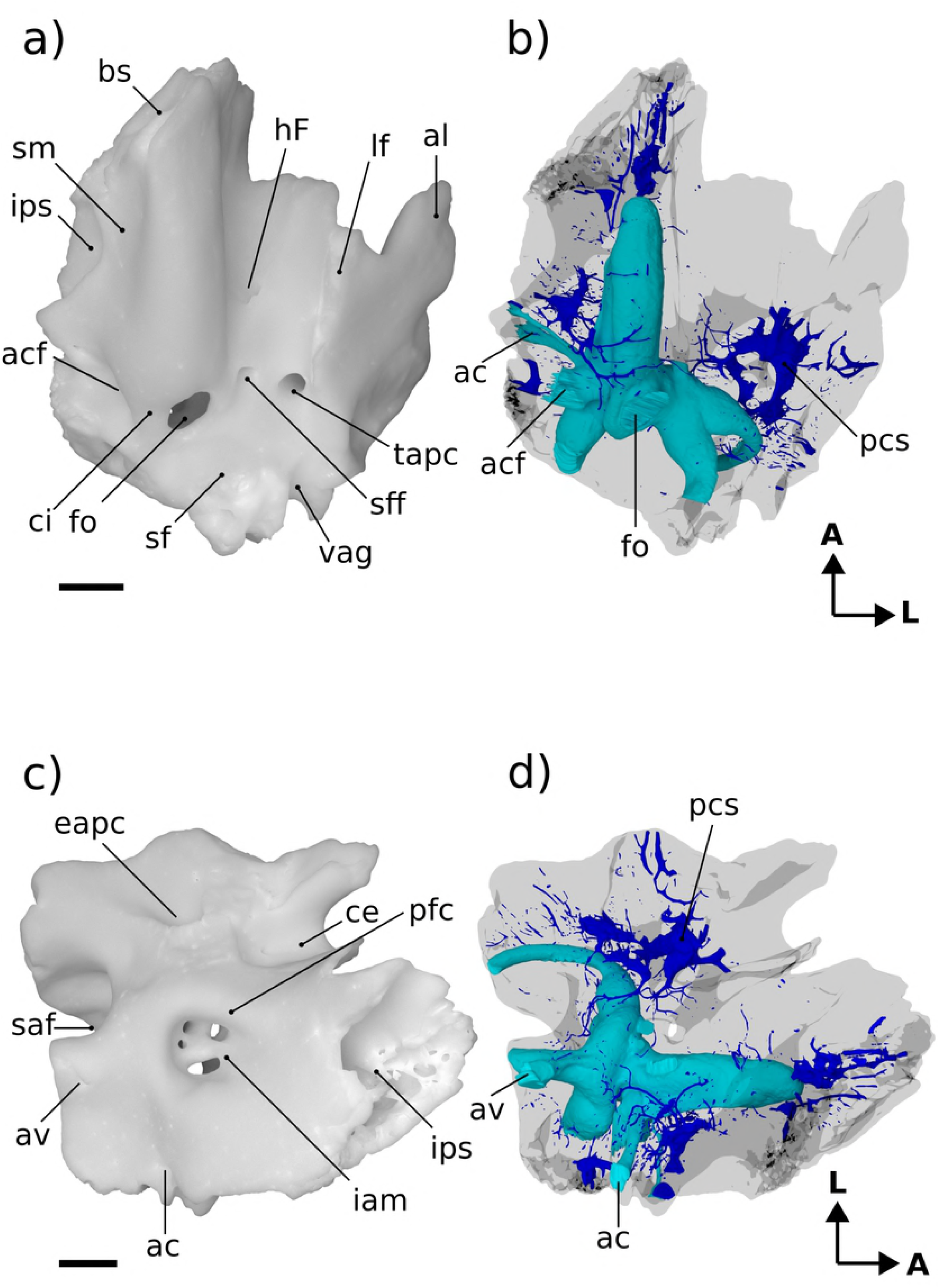
Renderings of Höövör 2 petrosal. a,b ventral view; c,d endocranial view. b,d showing cochlear endocast in green and circumpromontorial venous plexus in blue. **A**, anterior; **L**, lateral.

**Figure 3.**
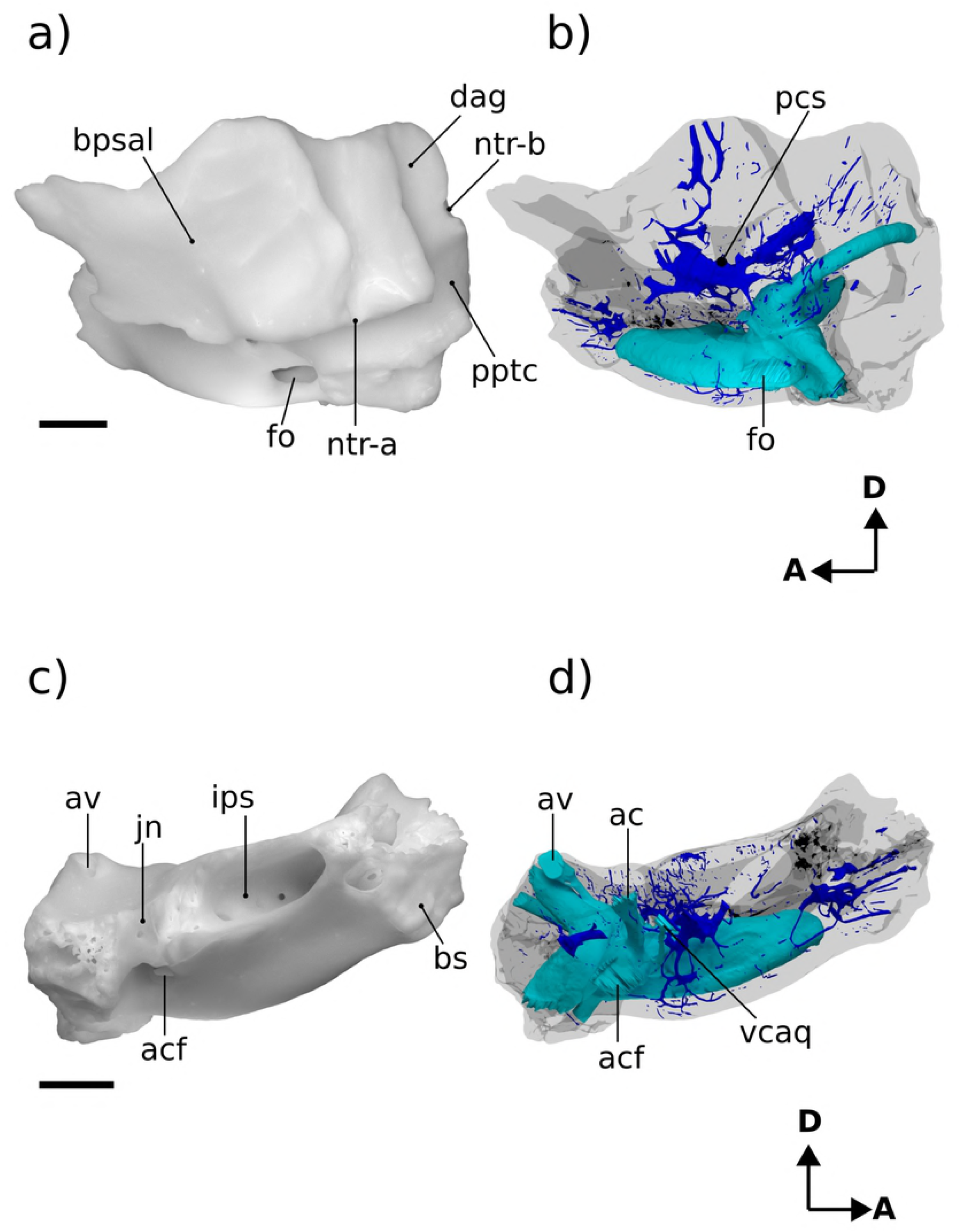
Renderings of Höövör 2 petrosal. a,b lateral view; c,d medial view. b,d showing cochlear endocast in green and circumpromontorial venous plexus in blue. **A**, anterior; **D**, dorsal.

The quality of preservation in H2 is unfortunately inferior to that seen in H1, in most places. Lost structures in H2 include, presumably, the full extent of the tympanohyal, crista parotica, paroccipital process, and caudal tympanic process; the posterior half of the subarcuate fossa and impression of the sigmoid sinus; and the post-temporal canal accommodating the arteria diploëtica magna. Conversely, several phylogenetically informative structures are better preserved in H2, in addition to several derived characters only seen in this taxon. In particular, the anterior extent of the promontoruim is apparent in H2, and the contiguous anterior lamina-lateral flange is better represented than in H1.

### Tympanic surface

The ventral expression of the pars cochlearis is the elongate and laterally steep promontorium (Fig 2a). Almost the full extent of this structure is preserved, except for minor damage at its anterior-most and medial margins. As such, it can be determined that the anterior limit of the promontorium abuts into an anteromedially facing planar suture, most likely with the basisphenoid bone. The dorsal extent of this sutural plane is obscured by damage exposing underlying venous sinuses and cancellous bone. Posteriorly, the promontorium is terminated by the posterolaterally facing fenestra vestibuli. The 0.92 mm horizontal width of the fenestra vestibuli occupies almost the full extent of the posterior aspect of the promontorium; with the posteriorly directed crista interfenestralis projecting from the posteromedial corner of the promontorium as well. In both H1 and H2 the anterolateral margin of the fenestra vestibuli is grooved to accommodate the footplate of the stapes.

The crista interfenestralis is concave ventrally, but generally oriented within the same horizontal plane as the body of the promontorium. The posterodorsal root of the crista interfenestralis (along with all other structures within the posterior one third of the petrosal) is incompletely preserved; however, the surface curvature in this region suggests that the crista interfenestralis did not contact the posterior processes within the mastoid region, and that it blended subtly to lose distinction within the post-promontorial tympanic recess.

Between the anterior and posterior terminations of the promontorium there are no surface impressions of the promontorial artery, stapedial artery, deep petrosal nerve (internal carotid nerve), or ramifications of the glossopharyngeal nerve. The length of the promontorium is therefore shaped into a smoothly rounded prism, with a sub triangular cross sectional profile. While the ventral edge of the promontorium is distinct (more so anteriorly) it does not show the level of salience seen in *Haldanodon* or *Megazostrodon* [3]. There is also no suggestion of a reduced rostral tympanic process (as in *Dasypus;* [21]) or impressions suggesting ventral contact of the promontorium with the lateral flange (as seen in multituberculates; [44]).

The lateral slope of the promontorium dips directly into the lateral trough, showing no distinct impression for the origin of the tensor tympani muscle anterior to the hiatus Fallopii, although the lateral trough is distinctly deeper in this region. Along the anterior two thirds of the promontorium its medial slope dips shallowly towards the sulcus medialis along the medial margin of the petrosal bone [42]. This area also contributes to the intramural enclosure of the inferior petrosal sinus dorsally. The medial aspect of the posterior one third of the promontorium curves directly into the ventral margin of the medially facing aperture of the cochlear fossula [20,21,45]. The aperture of the cochlear fossula lies external to the fenestra cochleae and is fully separated from the incipient perilymphatic canal (aqueductus cochleae) by a thin, horizontally oriented bony strut (the processus recessus; [46,47]); and is approximately 0.59 mm wide anteroposteriorly, and 0.75 mm high dorsoventrally. The space defined by the aperture of the cochlear fossula medially and the processus recessus dorsally leads anteromedially to a small anonymous venous canal (confluent with the inferior petrosal sinus and endocranial cavity) and to the jugular notch posteromedially.

The bony perilymphatic canal in both H1 and H2 is approximately 1.3 mm in length, however H2 shows a greater degree of sophistication of the process recessus structure because of the rounder, more ovoid, cross section of the perilymphatic canal it encloses. The perilymphatic canal in H1 in contrast is dorsoventrally flattened, and has much sharper anterior and posterior borders. Whether the more derived morphology in H2 represents ontogenetic or phylogenetic development is uncertain, however, as variation in the compression of the cochlear aqueduct is commonly seen between individuals of *Dasypus novemcinctus* [21].

Because of its location posterior-dorsal-lateral to the pars cochlearis, the pars canalicularis and mastoid region is exposed ventrally as an “L”-shaped depression, dorsally offset from the promontorium. The perpendicular vertex of this “L” is posterolaterally offset approximately 1.5 mm from the center of the fenestra ovalis. From this vertex, the mediolaterally oriented limb of the “L”-surface is damaged posteriorly, but the bases of three major topographic features are apparent. From lateral to medial these features are: 1) the ventral expression of the anterior lamina; 2) the open canal for the ventral ascending groove (for the proximal segment of the superior ramus of the stapedial artery); 3) the broken base of the paroccipital/mastoid region; and 4) the fossa for the stapedius muscle [30,48], placed directly posterior to the fenestra ovalis. Major damage to the petrosal medial to the stapedius fossa precludes the recognition of other structures; such as the pocket medial to crista interfenestralis as seen in H1 [3].

The posteroventral extent of the anterior lamina is placed caudal to the line demarcating the periosteal surface of the anterior lamina anteriorly from its sutural surface with the squamosal posteriorly. The lateral portion of the horizontal limb of the “L” therefore marks the zone of synostosis of the embryonic lamina obturans with the endochondral bone forming the bulk of the petrosal. Compared to the shape of this region in H1, the vertical line demarcating the periosteal and sutural surfaces of the anterior lamina in H2 is much less laterally offset from the rest of the petrosal. As suggested in [2] the petrosal’s wide, solid, and laterally projecting sutural surface with the squamosal in H1 suggests that the glenoid fossa and other structures associated with the dentary-squamosal contact was relatively robust; a condition seen in gobiconodontids because of their large and transversely widened mandibular condyle.

Medial to the tympanic commencement of the ventral ascending groove (for the ramus superior of the stapedial artery) is the damaged base of the paroccipital/mastoid region. The anatomical landmarks commonly found in this region include a ventrally projecting paroccipital process, a rostrocaudally oriented crista parotica extending anterior to it, and possibly a caudal tympanic process of the petrosal running mediolaterally from the paroccipital process. All evidence of these features has been effaced from H2 due to the horizontal fracturing of their common base medial to the tympanic commencement of the ventral ascending groove. Therefore, while the presence and condition of the crista parotica, paroccipital process and caudal tympanic process of the petrosal cannot be commented on, the shape of this common base does demonstrate several significant contrasts with this region in H1. Most importantly, as mentioned in [2], the sutural surface of the squamosal bone’s contact with the petrosal in H1 extends onto the lateral margin of the crista parotica, and therefore would have formed the bulk of the fossa incudis (the depression accommodating the crus brevis of the incus). In contrast, the intact lateral margin of the broken base of the paroccipital/mastoid region in H2 is vertically steep and lacks a sutural surface with the squamosal. It is unclear what the phylogenetic polarity of these characteristics differentiating H1 and H2 would be; however, it is likely that these differences are a direct result of the more robust attachment of the squamosal to the lateral surface of the petrosal in H1, and the more ventral location of the ventral ascending groove (relative to the fenestra ovalis) in H2. The medial-most structure visible along the mediolateral limb of the “L” is the fossa for the stapedius muscle. The center of this depression in H2 is located further medially (closer to the crista interfenestralis) than in H1 (where it is located diametrically opposite the fenestra ovalis).

Anterior to the tympanic commencement of the ventral ascending groove, the anteroposteriorly oriented limb of the “L”-shaped exposure of the pars canalicularis is located between the lateral aspect of the promontorium medially, and the medial aspect of the lateral flange (ventral extension of the ossified lamina obturans; [37]) laterally. The tympanic surface of the anteroposteriorly oriented limb therefore forms an elongate and concave sulcus, termed the lateral trough [36]. The lateral trough is an anatomical crossroads for several important neurovascular structures (described below), and is perforated by three major foramina. Between the anterior margin of the fenestra vestibuli and the posterior base of the lateral flange, two of these foramina are aligned mediolaterally at the posterior end of the lateral trough. The lateral of the two is the tympanic aperture of the prootic canal (for the prootic vein = middle cerebral vein; [26]). The tympanic aperture of the prootic canal does not approximate or become confluent with the ventral commencement of the canal for the superior ramus of the stapedial artery, as is seen in *Ornithorhynchus* and many multituberculates [49,44]. There is also a small, anonymous, vascular foramen placed anterolateral to the tympanic aperture of the prootic canal, that communicates with the circumpromontorial venous plexus, and has a small sulcus leading into it.

Medial to the tympanic aperture of the prootic canal is the more posteriorly directed secondary facial foramen for the entrance of the hyomandibular branch of the facial nerve into the cavum tympani. Compared to H1 and the spalacotheroid *Zhangheotherium* [38] the tympanic aperture of the prootic canal and the secondary facial foramen in H1 appears much closer. The location of these foramina are however ambiguous in the zhangheotheriid *Maotherium* [50]. In H1 the center of the secondary facial foramen is located anteromedial to the prootic aperture.

Anterior to the secondary facial foramen, the anteriorly oriented hiatus Fallopii perforates the lateral trough near the lateral margin of the promontorium. The hiatus Fallopii admits the palantine branch of the facial nerve (= greater petrosal nerve) in to the lateral trough. Therefore, the (approximately 1.28 mm long) lamina of bone extending between the secondary facial foramen and the hiatus Fallopii represents the bony floor of the cavum supracochleare (the space containing the geniculate ganglion of the facial nerve) and the floor of the hiatus Fallopii. Anterior to the hiatus Falopii the lateral trough is more deeply excavated and roughened, similar to the condition seen in H1. This surface may represent a relatively undeveloped area for the attachment of the tensor tympani muscle.

### Lateral Surface

The preserved squamosal surface of H2 is composed mainly of the anterior lamina anteriorly, with some exposure of the lateral surface of the pars canalicularis posteriorly (Fig 3a). Four grooves are also apparent on this surface, that represent the petrosal’s contribution to arterial canals enclosed laterally by the squamosal. Commencing near the posterior margin of the anterior lamina, posterior to the tympanic aperture of the prootic canal, the approximately 1.74mm long groove for the superior ramus of the stapedial artery (ventral ascending groove) curves posterodorsally in a smooth arc. Almost halfway along its length, this canal gives off a distributary branch to a groove for a minor temporal ramus (of the superior ramus) of the stapedial artery. The posterior termination of the groove for the superior ramus of the stapedial artery is its point of confluence with the groove for the arteria diploëtica magna (remnant of the post-temporal fenestra) and groove for the ramus supraorbitalis of the stapedial artery (dorsal ascending groove; [37]). The large size (0.70 mm diameter) of the canal for the arteria diploëtica magna suggests that this vessel was the major supplier of arterial blood to the cranial connective tissues in this region ([3]; see Fig 2a and Fig 4a and c). However, neither this groove nor the groove for the supraorbital ramus of the stapedial artery are preserved along their full extent or diameter, preventing the description of their precise distributions and possible ramifications. The visible extent of the groove for the supraorbital ramus of the stapedial artery runs vertically along the lateral surface of the petrosal, directly lateral to the endocranial expression of the subarcuate fossa. A subsidiary groove for a possible second temporal ramus of the stapedial artery may be seen branching from the posterior wall of the dorsal ascending groove (for the supraorbital ramus of the stapedial artery). However, this feature may be the result of postmortem fragmentation and rounding.

**Fig 4.**
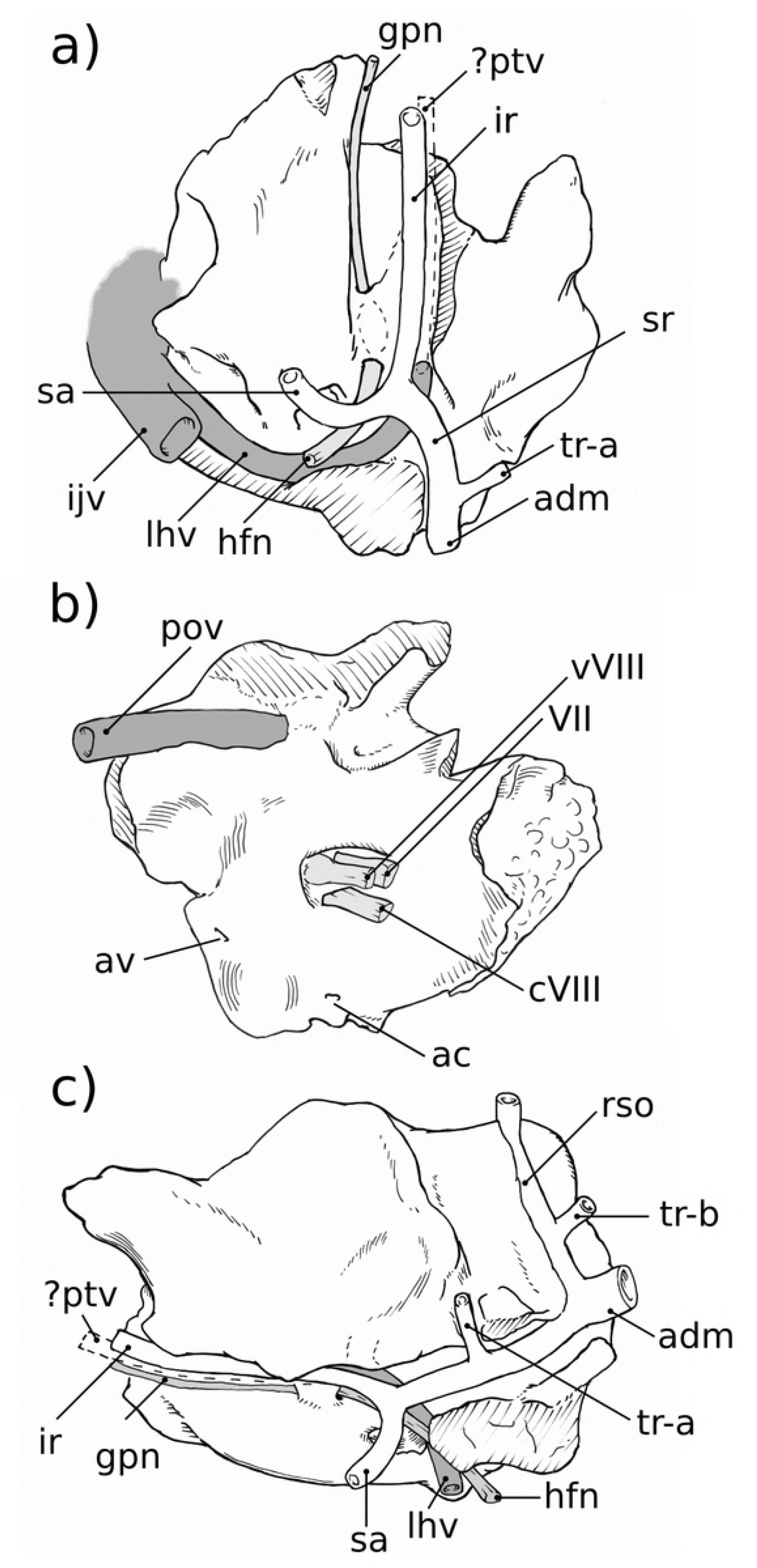
Neurovascular reconstructions of Höövör 2 petrosal. a, lateral view; b, medial view; c, lateral view. Venous structures shown in dark gray; nervous structures shown in light gray; the stapedial artery and its ramifications are unshaded.

Ascending laterally from the lateral flange, the lateral surface of the periosteal (non-sutural) surface of the petrosal is composed of the anterior lamina. The ventral extension of the lateral flange itself is damaged; however, it is probable that not much of its original surface has been lost, and therefore improbable that it could have supported a sutural contact with the quadrate ramus of the alisphenoid.

The lateral aspect of the anterior lamina does not show the mediolaterally projecting and horizontally flattened area (termed the anterolateral recess) seen in H1 [2]. Instead the lateral surface of the anterior lamina in H2 shows a steeper vertical slope, that terminates anteroventrally at the lateral opening of the cavum epiptericum (described below). The posterior extent of the anterior lamina contains a small vertical crest of bone lateral to the tympanic aperture of the prootic canal separating the periosteal and sutural surfaces of the periotic.

### Cerebellar surface

The preserved cerebellar surface of H1 (Fig 1b and Fig 2c) can be visualized as being composed of three endocranially exposed neural invaginations, anteriorly (the cavum epiptericum), medially (the internal acoustic meatus) and posteriorly (the subarcuate fossa) and a venous canal (the prootic canal) laterally; thus forming a “+” shaped pattern of topologically negative spaces in cerebellar view. The intervening elevated surface between these four structures therefore forms a matching “x” shaped pattern, the center of which being composed of a massive elevation of bone. Lateral to this “x” is a large fragment of the vertically oriented anterior lamina, and most of the area medial to the “x” is formed by the pars cochlearis enclosing the cochlear canal. The posteromedial border of the “x” however is formed by a thin sheet of bone near the jugular notch (i.e. the processus recessus, with some contribution from the petrosal bone proper). The endocranial aperture of the aqueductus cochleae (perilymphatic canal) can be seen at the anterior border of this sheet of bone.

The internal acoustic meatus in H1 is an approximately 1 mm deep invagination, terminated by four foramina distally. Because of its oblique angle of descent into the substance of the petrosal bone, the endocranial rim of the internal acoustic meatus is shaped differently in each of its four quadrants, providing a convenient pattern for the description of its contents.

The prefacial commissure (also suprafacial commissure; [20]) is the name given to the portion of the anterior ossified wall of the internal acoustic meatus enclosing the proximal bony conduit of the facial nerve (or aquaductus Fallopii; [30]), and which is not first preformed developmentally in the chondrocranium [20]. In both H1 and H2 the prefacial comissure extends posterolaterally from the shallower curvature of the posterior cranial fossa, and terminates laterally at the lateral-most point of the proximal aperture of the internal acoustic meatus. The entire 90° arc of the anterolateral quadrant of the internal acoustic meatus in H2 can therefore be thought of as consisting of the prefacial comissure, which extends distally (beyond the internal acoustic meatus) to form the anterior wall of the approximately 0.5 mm diameter primary facial foramen (pff).

Directly lateral to, and contiguous with, the prefacial commissure the endocranial surface (outside the internal acoustic meatus) shows a blunt, anteroposteriorly oriented crista petrosal separating the bulk of the cerebellar surface of the petrosal from the petrosal’s contribution to the bony floor of the cavum epiptericum (accommodating the trigeminal ganglion=semilunar ganglion=Gasserian ganglion; [20]). Being the anterior attachment of the tentorium cerebelli in extant mammals, the location of the crista petrosa near the medial margin of the cavum epiptericum likely marks the point of dorsal enclosure of the trigeminal ganglion into the dural folds separating the middle and posterior cranial fossae in H2. In more plesiomorphic forms such as *Morganucodon* [30] and *Priacodon* [3], and in modern *Ornithorhynchus* [51], the prefacial comissure itself forms the medial margin of the cavum epiptericum. Specimens H1 and H2 show a derived location of the crista petrosa laterally offset from the prefacial comissure, and dorsal to the lamina of bone forming the dorsal roof of the lateral trough. This morphology is likely an osteological byproduct of the mediolateral dilation of the endocranial space within the posterior cranial fossa.

The anteromedial quadrant of the internal acoustic is a continuation of the wider curvature of the surrounding pars cochlearis, and so does not form a distinct lip for the meatus. However, in H1 and H2 the ventral surface of the internal acoustic meatus in this region contains a low ridge of bone running distally into its depths, the crista transversa [21]. This low ridge loses distinction before reaching the distal terminus of the meatus (as therefore should not be called the “falciform crest”), but forms a separation between the foramen for the cochlear nerve (in the posteromedial quadrant) and the primary facial foramen (in the anterolateral quadrant). The crista transversa is distinctly higher in H1 and longer in H2, but in neither does it reach the height or salience of the falciform crest seen in *Homo* [41] and other modern mammals. The foramen for the cochlear nerve is ovoid, approximately 0.74 mm long along the axis of the cochlear canal and 0.3 mm wide mediolaterally; and shows smooth margins with no development of a tractus foraminosus (or cribriform plate). Dorsal to the foramen for the cochlear nerve, the distal surface of the internal acoustic meatus contains the two circular foramina for the branches of the vestibular nerve, mentioned above. Based on the orientation and location of these two foramina for the vestibular nerve it is likely that they are homologous to the foramina for its utriculoampullar and sacculoampullar branches [17] seen in both extant therians and monotremes.

In both H1 and H2 the posterolateral and posteromedial quadrants of the internal acoustic meatus are formed by the raised wall of bone partitioning the internal acoustic meatus from the subarcuate fossa. The ventral floor of the internal acoustic meatus in this region contains the anteroposteriorly elongate foramen for the cochlear nerve medially, and two smaller circular foramina laterally or posterolaterally. As mentioned above, these two circular foramina are inferred to have transmitted the utriculoampullar branch (larger, and more lateral) and sacculoampullar branch (smaller, and more medial) of the vestibular nerve [17]. These inferred homologies are supported by the trajectories of these foramina when viewed on the virtual endocast of the bony labyrinth; however, because these vestibular foramina do not form cylindrical canals leading directly to their peripheral targets of the vestibular nerve the precise targets of innervation of the nerves traversing the vestibular foramina cannot be determined conclusively.

Despite this uncertainty, it is important to highlight that the morphology of the floor of the internal acoustic meatus in H2 likely represents an intermediate condition between that seen in H1 and fossil cladotherian mammals. In Mesozoic cladotherians [52,6,5], the foramen acusticum superius is comprised of the discrete depression within the internal acoustic meatus dorsolateral to the crista trasversa, that encompasses the proximal (endocranial) apertures of the primary facial foramen and foramen for the utriculoampullar branch of the vestibular nerve (this region is named “superior vestibular area” in [52], and its distal extent in modern therians is termed the area cribrosa superior). Even though fossil and extant cladotheres unanimously show the apomorphic distribution of peripheral axons within the vestibulocochlear nerve by the formation of osseous cribriform areas within preexisting foramina (the tractus foraminosus forming within the foramen transmitting the cochlear nerve being the most prominent example); most of the contents of the foramen acousticum superius remain free of trabeculated bony outgrowths in crown therians and all fossil cladotheres so far studied. The foramen acousticum inferius on the other hand comprises the common depression for the foramen of the cochlear nerve and the foramina for nerves targeting the saccular macula (penetrating through the macula cribrosa media; [53] and ampulla of the posterior semicircular canal (through the foramen singulare). The contents of the foramen acousticum inferius commonly recruit cribriform bony structures to secondarily infill these ancestral foramina in cladotheres. Additionally, because of its incorporation of a branch of the vestibular nerve (the sacculoampullar branch, or inferior vestibular nerve in *Homo)* along with the cochlear nerve, the foramen acousticum inferius is typically at least twice as large in areal extent than the foramen acousticum superius in Cladotheria. In contrast, the area likely homologous to the foramen acousticum inferius in *Priacodon* and H1 is subequal or smaller than the foramen acousticum superius, and completely lacking cribriform bony infilling. The internal acoustic meatus of H2, however, shows that the foramen for the sacculoampullar branch of the vestibular nerve has rotated posteroventral (relative to H1) to a position inferior to the crista transversa (or a crest projecting posterior from the crista transversa) and therefore is incorporated into the foramen acousticum inferius. Therefore, while not showing a tractus foraminosus or other cribriform bony infilling, H2 shows the characteristic of having a foramen acousticum inferius being approximately twice as large as the foramen acousticum superius, a trait mentioned by [52] as showing the apomorphic condition of their cladotherian specimen relative to *Priacodon.* The relocation of the sacculoampullar branch of the vestibular nerve ventral to the crista transversa, and the formation of a tractus foraminosus are also both seen in extant monotremes.

Directly posterior to the internal acoustic meatus is a wall of bone that forms the medial margin of the subarcuate fossa, enclosing the primary common crus of the bony labyrinth, and supporting the endocranial aperture of the aquaductus vestibuli (containing the membranous the endolymphatic duct). The distal half of this medial wall has been lost in H2 due to postmortem fracturing; and features such as the impression of the sigmoid sinus and contact with the exoccipital bone cannot be confirmed. However, the preserved morphology of this wall provides a reliable estimate of the maximal diameter of the proximal entrance to the subarcuate fossa. The proximal aperture of the subarcuate fossa is the endocranial expression of the anterior semicircular canal, and its 1.86 mm diameter in H2 closely matches the same length measured in H1. However, in both specimens it is apparent that the distal extent of the subarcuate fossa has a significantly larger diameter because of its medial excavation of its medial wall, distal to the constriction formed by the anterior semicircular canal. The medial diversion of the subarcuate fossa projects into the loop of the posterior semicircular canal, similar to the way the proximal aperture of the subarcuate fossa itself projects into the the loop of the anterior semicircular canal. The state of preservation in H2 does not allow the maximal diameter of the distal subarcuate fossa to be confidently estimated.

Anterolateral to the subarcuate fossa, and medial to the posterior region of the anterior lamina, is the endocranial aperture of the prootic canal. A faint groove leading into this aperture can be seen following the curvature of the anterolateral margin of the subarcuate fossa for a short distance before leaving the preserved edge of the petrosal. This structure is termed the groove for the middle cerebral vein in [30], and represents the approximate branching point of the transverse sinus from the middle cerebral vein. The enclosed prootic canal in H2 is substantially shorter (1.72 mm, versus 2.55 mm in H1) and less sigmoidal than the bony canal seen in H1. The diameter of the canal in both specimens is approximately 0.6mm along its entire length (Fig 2c and Fig 5a).

**Fig 5.**
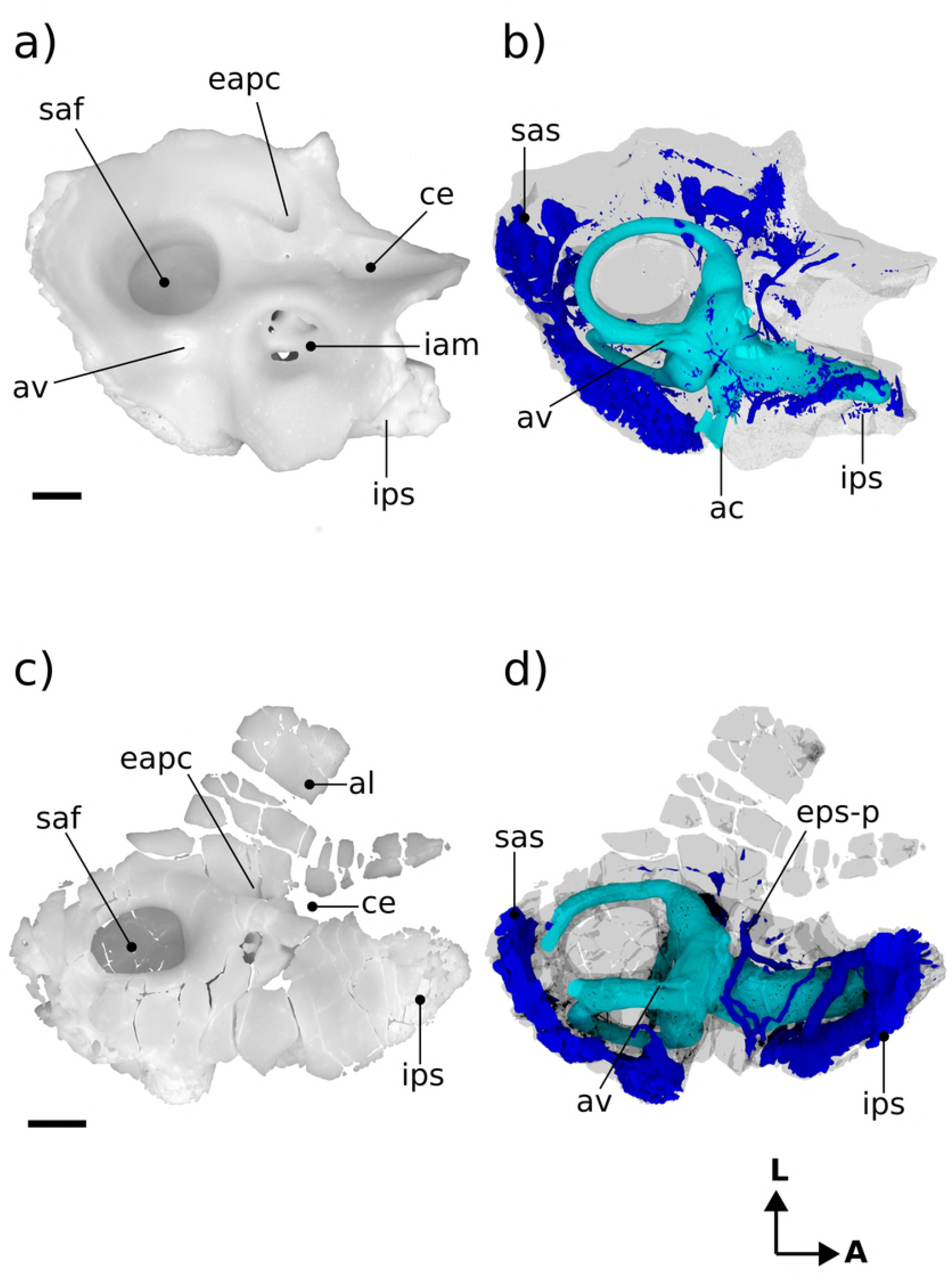
Renderings of stem therians in endocranial view. a,b Höövör petrosal 1 (reflected); c,d *Priacodon.* Showing cochlear endocast in green and circumprontorial venous plexus in blue. Scale bar is 1 mm. **A**, anterior; **L**, lateral.

The petrosal bone’s enclosure of the cavum epiptericum is better preserved in H2 than in H1, allowing for a more precise characterization of this phyogenetically significant structure. As such, in H2 it can be determined that the impression for the cavum epiptericum on the petrosal is rectangular in general dimensions, with its posterior margin formed by an anteriorly facing border of bone placed anteromedial to the endocranial aperture of the prootic canal (this wall may represent the original anteriolateral margin of the ossified otic capsule before its subsequent fusion with the ossified lamina obturans). Anterior to this posterior margin the bony floor of the cavum epiptericum extends approximately 1.64 mm anteriorly before terminating at the anterior margin of the petrosal. As mentioned above, the medial and lateral margins of the cavum epiptericum are formed by the crista petrosa and anterior lamina, respectively. These structures define the 1.19 mm width and 0.86 mm depth of the cavum epiptericum and contribute to two foramina communicating with this space. The posteriormedial corner of the cavum epiptericum contains the semilunar hiatus [37], that extends 0.5 mm medially within the crista petrosa to communicate with the cavum supracochleare. The anterolateral margin of the petrosal’s contribution to the cavum epiptericum is formed by an emargination of the broken rostral margin of the anterior lamina. This emargination would have comprised the majority, or entirety, of the margin of the foramen for the mandibular branch of the trigeminal nerve, and hence would be termed the foramen ovale (or the foramen pseudoovale). The preserved posterior margin of the foramen ovale is an approximately 1.19 mm diameter semicircular indentation. However, preservation prevents the determination of whether this foramen is entirely contained within the anterior lamina, and what its total anteroposterior length would have been.

### Neurovascular Reconstruction

Figure 4 illustrates reconstructions of the major vascular ramifications on the external surface of the H2 petrosal. The vessels for which osteological correlates can be observed include tributary veins of the internal jugular circulation and distributary arteries of the stapedial/occipital circulation [54,55]. While not leaving a distinct impression on the fenestra ovalis, the stapedial artery likely ran laterally to a bifurcation point slightly posterior to the tympanic aperture of the prootic vein. Running posterior from this bifurcation the superior ramus of the stapedial artery occupies the ventral ascending groove, then forms an anastomosis with the wider arteria diplöetica magna, and also gives off a small temporal ramus near its anterior extent. Because of damage to the dorsolateral extent of the petrosal forming the dorsal ascending groove, the course of the ramus supraorbitalis extending from the confluence of the arteria diplöetica magna and superior ramus of the stapedial artery cannot be reliably reconstructed. Likewise, the occipital artery, which likely contributed the majority of arterial blood to the cranial connective tissues through its confluence with the arteria diplöetica magna and elsewhere, has no osteological correlate within the preserved morphology of H2.

The two main tributaries of the internal jugular vein are the lateral head vein and inferior petrosal sinus [49,26]. As will be discussed below both the prootic sinus (middle cerebral vein), before its confluence with the lateral head vein, and the inferior petrosal sinus receive minor venous tributaries along their course through the body of the petrosal bone. Because there is no anterior emargination of the tympanic aperture of the prootic canal, H2 shows no osteological correlate of the post-trigeminal vein, leaving its existence in this taxon uncertain (it is therefore shown only with a dashed outline in Fig 4a and c). If, as in extant therians, H2 lacked a post-trigeminal vein the boundary between the prootic sinus and lateral head vein would become completely arbitrary, and in this report it is taken to be at the point of emergence of the prootic sinus onto the tympanic surface of the petrosal. A small venous foramen anterolateral to the tympanic aperture of the prootic sinus likely conducted a small tributary to the lateral head vein as well.

In most respects the external vascular reconstruction of H2 is broadly similar to that provided by [2] for H1. The most salient contrasts being the relatively more ventral position of the ventral ascending groove (and therefore the superior ramus of the stapedial artery) relative to the foramen for the arteria diplöetica magna in H2. Given the uncertainty and variability in the presence and branching pattern seen in temporal rami, the different reconstructed connectivity of the posterior temporal ramus in H1 (at the point of confluence of the arteria diplöetica magna and superior ramus of the stapedial artery) and H2 (from the ventral extent of the supraorbital ramus) should only be considered tentative.

## Comparison of labyrinthine endocast morphology between *Priacodon*, the Höövör petrosals, and extant mammals

### Pars Cochlearis

The straight distance along the bony cochlear canal, measured from the anterior surface of the recessus sphericus (caudal apex of the saccular expansion; [23]) to the distal terminus (anterior apex) of the cochlear canal, is approximately 0.8 mm shorter in *Priacodon* than in either of the Höövör petrosals. However, the differences in length of the cochlear canal in all three of these closely match the intra-specific variation seen in cochlear canal length reported in *Ornithorhynchus* (CL-cp; [17]), although *Ornithorhynchus* is much larger bodied than any of these stem therians. Also, despite being shorter, the shape of the cochlear canal in *Priacodon* can be still be interpreted as more derived than the cochleae seen in the Höövör petrosals because of its slightly stronger lateral curvature (see Figs 2–9).

**Fig 6.**
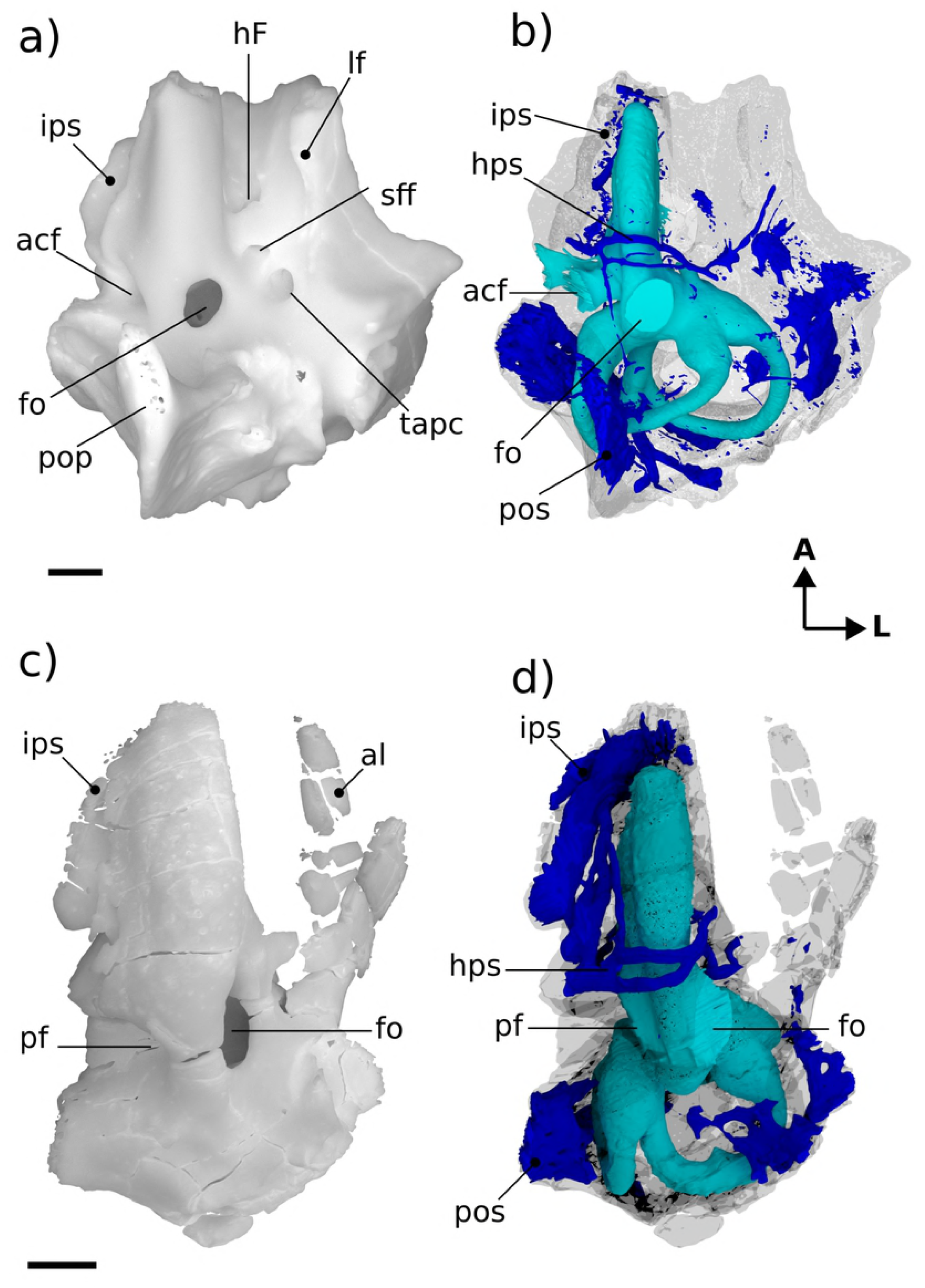
Renderings of stem therians in ventral view. a,b Höövör petrosal 1(reflected); c,d *Priacodon.* Showing cochlear endocast in green and circumpromontorial venous plexus in blue. Scale bars are 1 mm. **A**, anterior; **L**, lateral.

**Fig 7.**
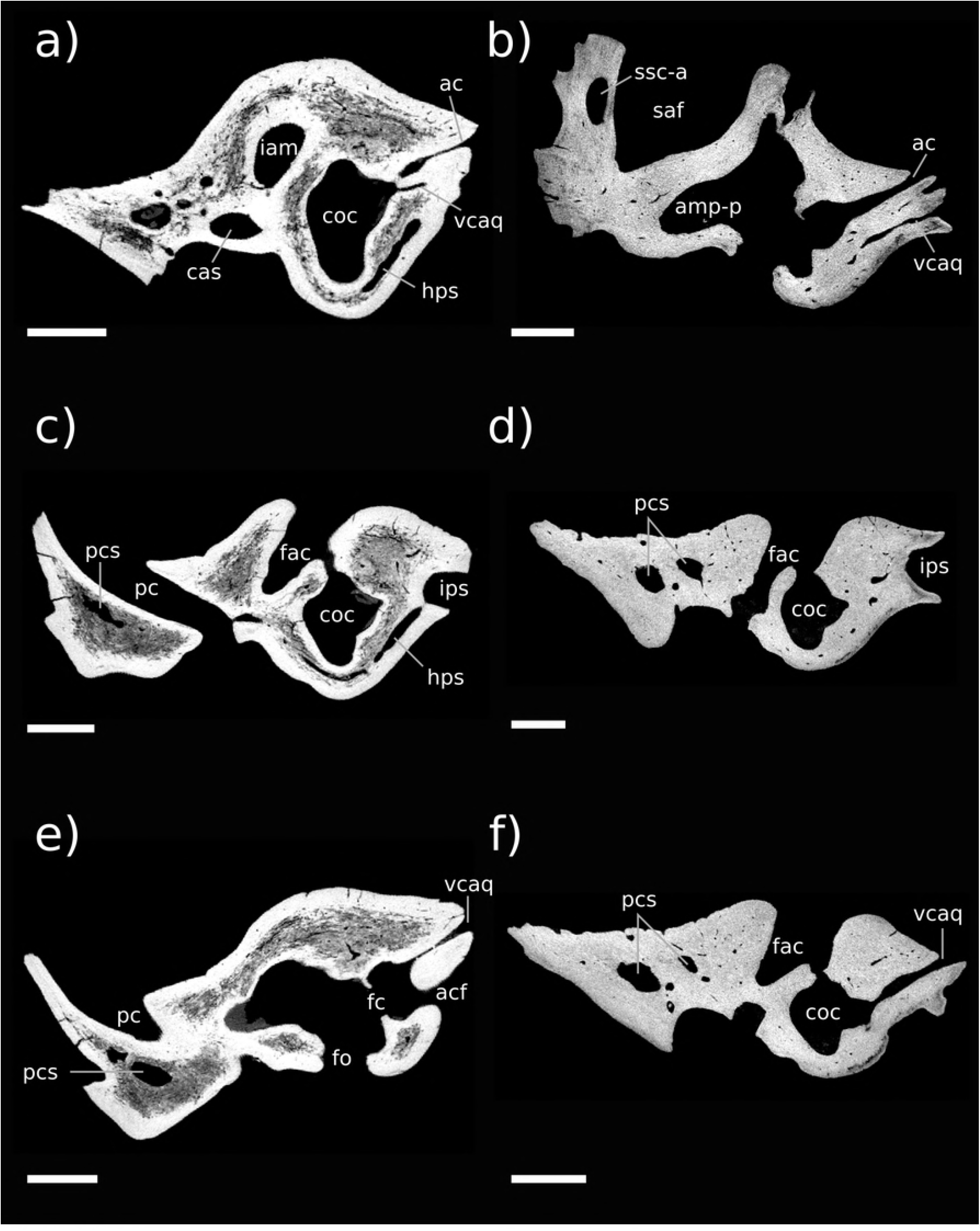
Resliced CT images of Höövör petrosals. a, c, e Höövör petrosal 1; b, d, f Höövör petrosal 2. a, b are oblique planes through both the cochlear aqueduct and canal for the vein of the cochlear aqueduct; c, d, e, f are coronal planes through the promontorium. C, d are taken from a more posterior plane than e and f to show the hypopromontorial sinus in H1. In all images left is lateral and dorsal is approximately toward the top of the page, all scale bars are 1 mm.

**Fig 8.**
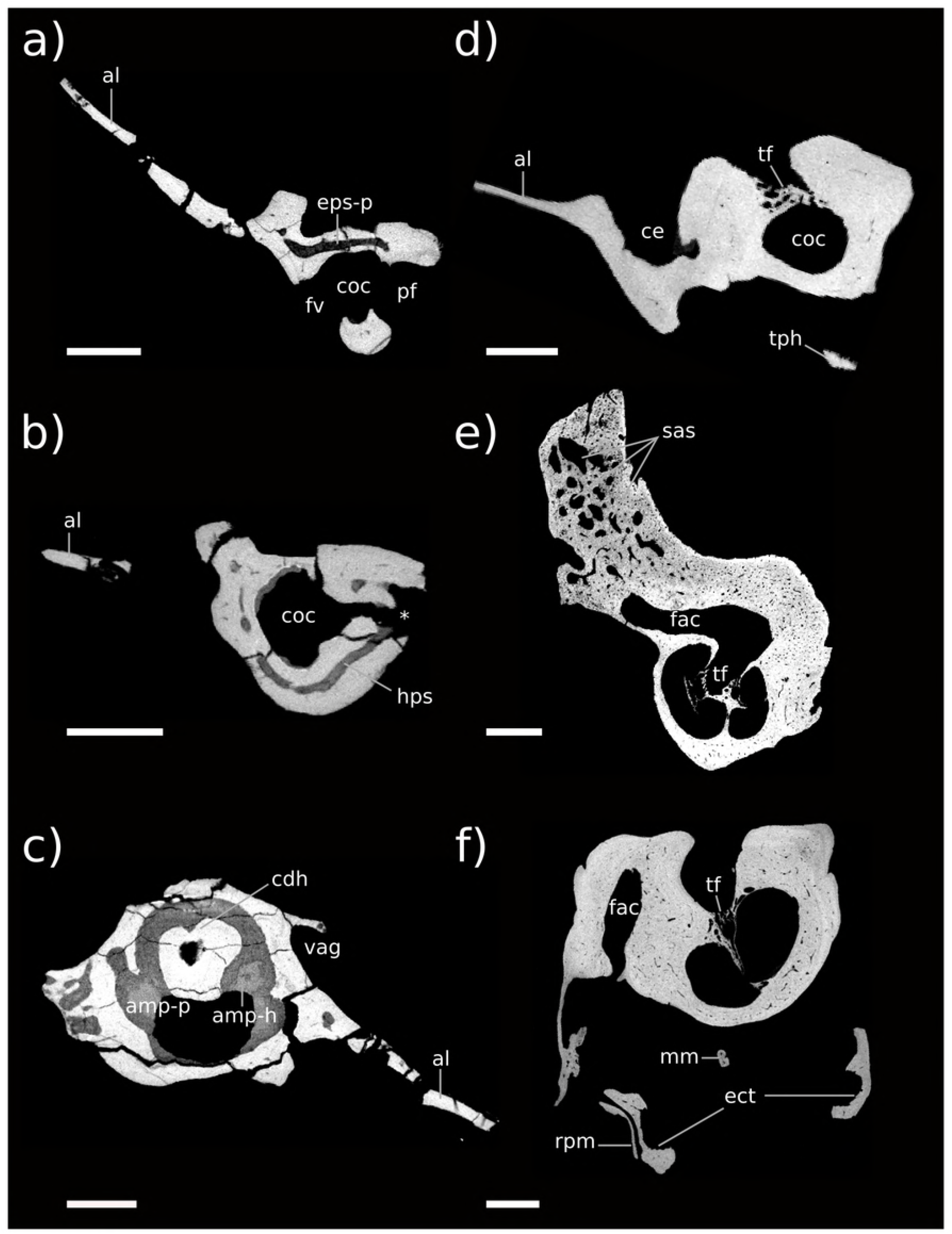
Resliced CT images showing *Priacodon* and several extant mammals. a, b, c images of *Priacodon;* d, coronal section through promontorium of *Ornithorhynchus;* e, coronal section of promontorium of *Didelphis;* f, coronal section through promontorium of *Dasypus.* a, b, are coronal sections through the promontorium of *Priacodon*, a is taken posterior to b to show the posterior epipromontorial sinus; b, is taken rostral to the fenestra vestibuli and perilymphatic foramen to show the hypopromontorial sinus. c, shows horizontal plane through the horizontal semicircular canal and its centripetal diverticulum. All scale bars are 1 mm, in a, b, d, e, f lateral is toward the left of the page and dorsal is toward the top of the page; in c lateral is toward the right of the page and posterior is toward the top of the page. Asterisk shows location of damage to anterior wall of perilymphatic foramen in *Priacodon.*

**Fig 9.**
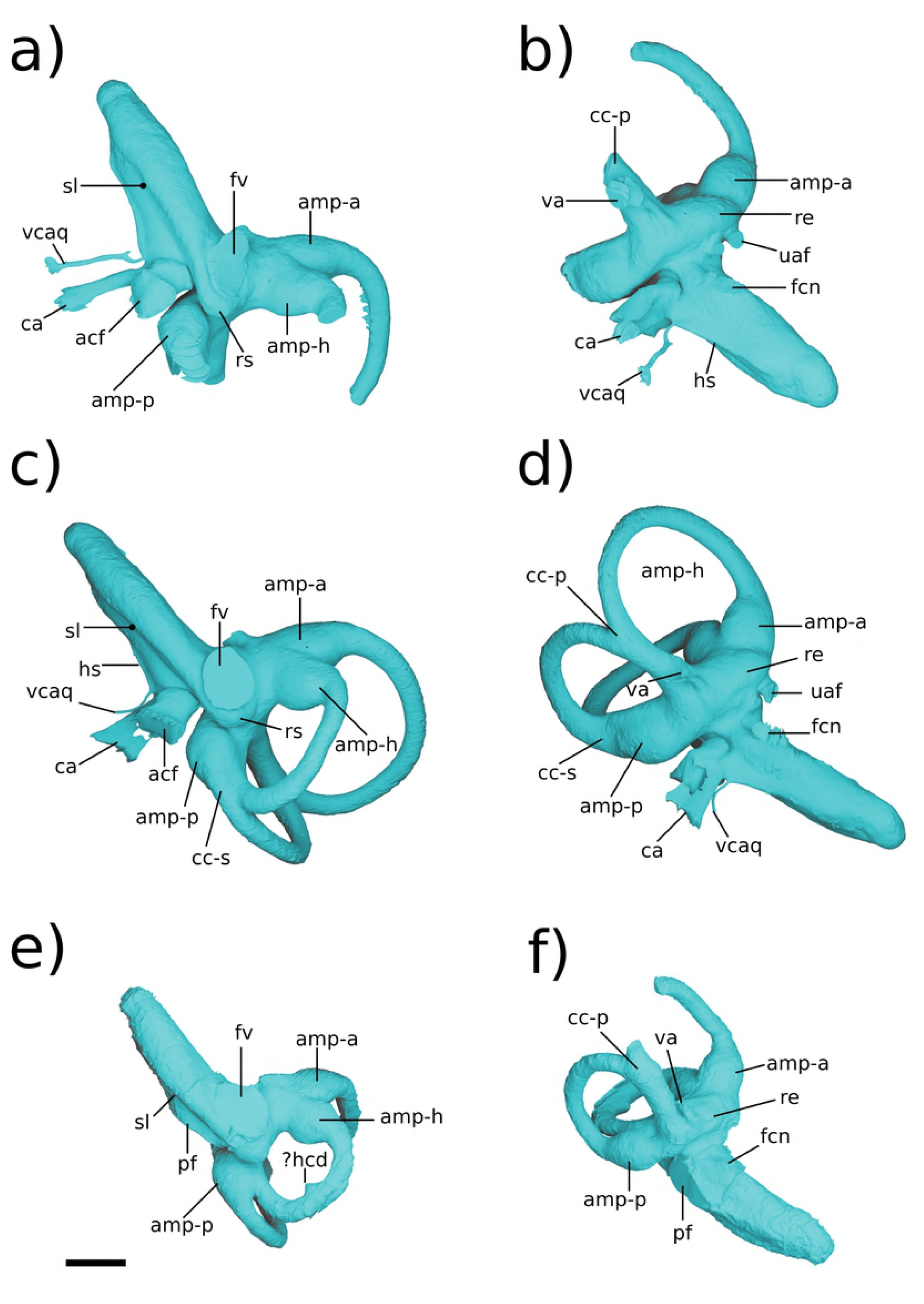
Renderings of labyrinthine endocasts in stem therians. a,b, Höövör petrosal 2; c,d Höövör petrosal 1 (reflected); e,f, *Priacodon.* All specimens shown as left-sided, scale bar is 1 mm.

While being a particularly homoplastic character, lateral curvature of the cochlear canal is seen only in mammaliaformes (especially the most plesiomorphic forms); with other amniotes developing medial curvature (i.e. convex towards the insertion of the cochlear nerve) to accommodate cochlear elongation. However, the incipient cochlear curvature in *Priacodon* only shows lateral deflection near its base, and no dorsoventral coiling. Additionally, the fact that similar, or stronger, degrees of cochlear curvature are reported for mammaliaformes outside the mammalian crown group [56] presents the possibility (dependent on the precise phylogenetic interrelationships hypothesized for the taxa involved) that loss of lateral cochlear curvature may actually be an apomorphic feature of the taxa represented by the Höövör petrosals and later stem therians.

All three of the stem therian endocasts (Fig 9) show that the cochlear canal tapered somewhat towards its distal terminus (much more so in the Höövör petrosals than in *Priacodon).* In particular, none of these endocasts show prominent inflations or emarginations of the cochlear canal capable of accommodating an enlarged lagenar macula. While the loss of its osteological correlate does not logically implicate the absence of a functional lagenar macula in these taxa [14], the morphology of the cochlear canal in these cases at least presents the possibility that these taxa had attained a terminal helicotrema (as in modern therians). This contrasts with the large, terminally positioned lagena and related nervous structures that are apparent in the osseus morphology of monotremes, several multituberculates, and mammaliaformes outside of the mammalian crown group [56]. Significantly, none of these stem therian endocasts show any of the “lagena related osteological characters”, as detailed by [57] for *Haldanodon;* such as a sulcus or canal for the lagenar branch of the cochlear nerve, a fossa for the lagenar macula, and/or canaliculi perforating terminal portion of the cochlear canal for dendrites innervating the lagenar sensory epithelium. The relative restriction of the apex of the cochlear canal suggests the progressive reduction of the lagena within progressively more crownward members of the stem therian lineage. Conversely, the bony accommodation of lagenar function would therefore be a retained symplesiomorphy within allotherians and modern monotremes. All three stem therian endocasts also lack the “sunken” position of the fenestra ovalis relative to the basal commencement of the cochlear canal, described by [52]. The sunken appearance of the fenestra ovalis was mentioned by these authors as being possibly apomorphic for the clade Cladotheria. However, given that this feature is also lacking in the South American cladothere *Vincelestes*, without a wider-scale phylogenetic analysis it seems at least equally plausible that an inset fenestra ovalis may be an apomorphic feature derived within dryolestoid cladotheres or some more exclusive group. A phylogenetic analysis informed by a large sample of cladotherian cochlear endocasts would be required to resolve the ancestral reconstruction of this feature.

One of the most marked features seen in the Höövör petrosals (Fig 7e and f) and not *Priacodon(* Fig 8a and b) and *Ornithorhynchus* (Fig 8d), is the complete segregation of the bony canal supporting the perilymphatic duct from the ossified aperture suspending the secondary tympanic membrane (the fenestra cochleae) and the aperture of the cochlear fossula. The secondary tympanic membrane is a thin epithelial bilayer found in many, if not most, amniotes [51], and segregates the fluid filled leptomeningeal/subarachnoid space from an air-filled intracranial space (such as a cavum tympani). The bony segregation of the secondary tympanic membrane from its confluent perilymphatic duct is, however, a characteristic seen only in advanced stem therians and in mature adult tachyglossids. The bony process of perichondral bone that completes the enclosure of the fenestra cochleae and aqueductus cochleae in the cohclear endocast of therians is termed the processus recessus. The processus recessus therefore ventrally floors a portion of the dorsolateral extent of the cochlear fossula (visible in the stem therian endocasts as a cylindrical inflation between the aperture of the cochlear fossula and the fenestra cochleae [46,47,10,21,45].

The performance implications of this partitioning of the proximal scala tympani are uncertain ([16]: chapter 10). For instance, the processus recessus may prevent the unconstrained flux of perilymph from the scala tympani into the subarachnoid space. Whatever its functional advantage, it is very likely that the homoplastic distribution of the processus recessus in stem therians and in older adult tachyglossids is due to convergence [44,17]. Additionally, the incipient expression of an incomplete “processus recessus” has also been recognized in *Priacodon* [3]; where it forms a linear ridge associated with the reconstructed course of the perilymphatic duct; and many multituberculate taxa [49] also show a variable recessed or enclosed groove for the perilymphatic duct.

Otherwise, the presence of a well-developed processus recessus, fenestra cochleae, and aquaductus cochleae is known among members of the clade Trechnotheria (including the spalacotheres, dryolestoids, and therians; [33]), and in the derived “triconodont” clade Gobiconodontidae (GWR Pers Obs). The consistent retention of these structures in almost all known trechnotherian mammals (Sirenia, Elephantomorpha, and Eschrichtiidae being the main exceptions, due to atavistic reversal; [58]) suggests that once developed this suite of characters provides an important functionality. However, despite its presumed adaptive significance, the tympanic aperture of the scala tympani, (whether this is the perilymphatic foramen as in *Priacodon*, or fenestra cochleae as in the Höövör petrosals) is of similar length width and perimeter in all three stem therians described here.

The impression of the scala tympani on the cochlear endocast (Fig 9) is confluent with the fenestra cochleae and aquaductus cochleae. It is visible in the stem therian endocasts as a medial inflation of the cochlear canal, delimited ventrally by the base of the bony secondary lamina. The posterior margin of this space is further inflated as it meets the anterior margin of the cochlear fossula [46,47,51]. However, because of the lack of a tractus foraminosus, Rosenthal’s canal (spiral ganglion canal), or primary bony lamina in these taxa, the scala tympani does not leave recognizable anterior and dorsal boundaries on the cochlear endocast. The choice of interpretation as to the presence or absence of a vestigial lagenar macula also greatly influences the inferred placement of the scala tympani in the apical areas of the cochlear endocast.

In these short straight cochlear endocasts, the impression of the scala tympani has the profile of a right triangle when viewed ventrally; with the anterior rim of the fenestra cochleae/perilymphatic foramen and the secondary bony lamina forming the triangle’s two perpendicular limbs. The hypotenuse of this triangle is formed by the medial contour of the cochlear endocast, which in the Höövör petrosals is also the location of a half-pipe shaped sulcus (i.e. a cylindrical prominence on the endocast; Fig 9a,c), that originates immediately anterodorsal to the cochlear fossula. In *Priacodon*, damage to the rim of the perilymphatic foramen hinders the reconstruction of the endocast here (Fig 8b). However, it can be confirmed that there is no vascular sulcus along the medial margin of its cochlear canal. Consequently, for *Priacodon* the medial hypotenuse of the triangle slopes anterolaterally more steeply than in the Höövör petrosals, causing the impression of the scala tympani to be limited to the proximal half of the cochlear canal (Fig 9e,f). In the Höövör petrosals the hypotenuse of the triangle has a much more gradual slope, causing the impression of the scala tympani to terminate more distally (approximately three quarters of the length along the cochlear canal). Distally, the impression of the scala tympani in the Höövör petrosals also shows a round secondary inflation, that interrupts the otherwise smooth lateral slant of the hypotenuse representing the medial contour of the cochlear canal. The half-pipe shaped medial sulcus is more distally extensive in H2 than H1, and can be followed along the complete length of the impression of the scala tympani. In H1 the medial sulcus loses distinction approximately half way along the length of the impression of the scala tympani, proximal to its slight terminal inflation.

In both Höövör petrosals (Fig 9a-d) the proximal commencement of the half-pipe shaped medial sulcus within the impression of the scala tympani is confluent with the emergence of two tubular structures from the cochlear endocast. The anterior tubular structure is a small venous canal that, for reasons outlined below, is inferred to have contained the “Vein on the Cochlear Aquaduct” (Fig 7a,b,e,f;[59] also see [45]), and is therefore homologized with the bony “Canal of Cotugno” [60]. In many extant therians this canal follows a tortuous mediolateral trajectory to contact the inferior petrosal sinus (Figs 10–12; [59]). In the Höövör petrosals, given the observed medial connectivity of this canal with a lateral diverticulum of the intramural inferior petrosal sinus, and it medial continuity with the half-pipe shaped sulcus on the impression of the scala tympani (Fig 9a-d), it is very likely that this medial sulcus transmitted venous components as well. In modern therians structures in this region represent the sole (or major; see [61]) outlet of venous blood from the pars cochlearis, and provide a subsidiary role in draining the pars canalicularis (Fig 12a, b and c). In the Höövör petrosals the half-pipe shaped medial sulcus on the impression of the scala tympani would then contain the homolog of what is the common cochlear vein (or one of its ramifications) in extant therian mammals.

**Fig 10.**
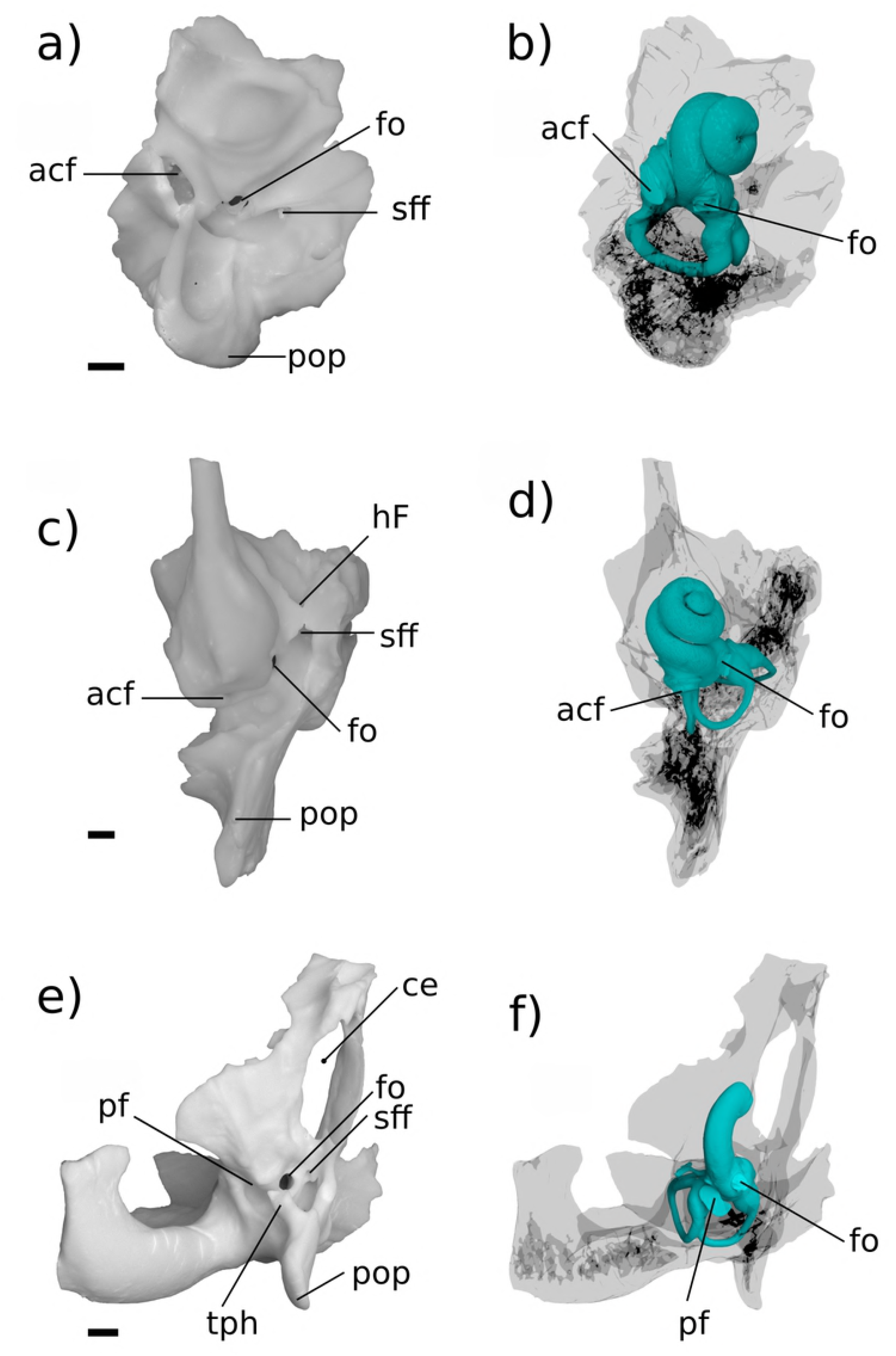
Ventral view of comparative mammalian specimens. a,b *Erinaceus;* c,d *Didelphis;* e,f *Ornithorhynchus.* All specimens are left-sided, venous sinuses are not shown. In e,f a fragment of the exoccipital bone is synostosed to petrosal. Medial is toward the left, rostral is toward the top of the page. All scale bars are 1 mm.

**Fig 11.**
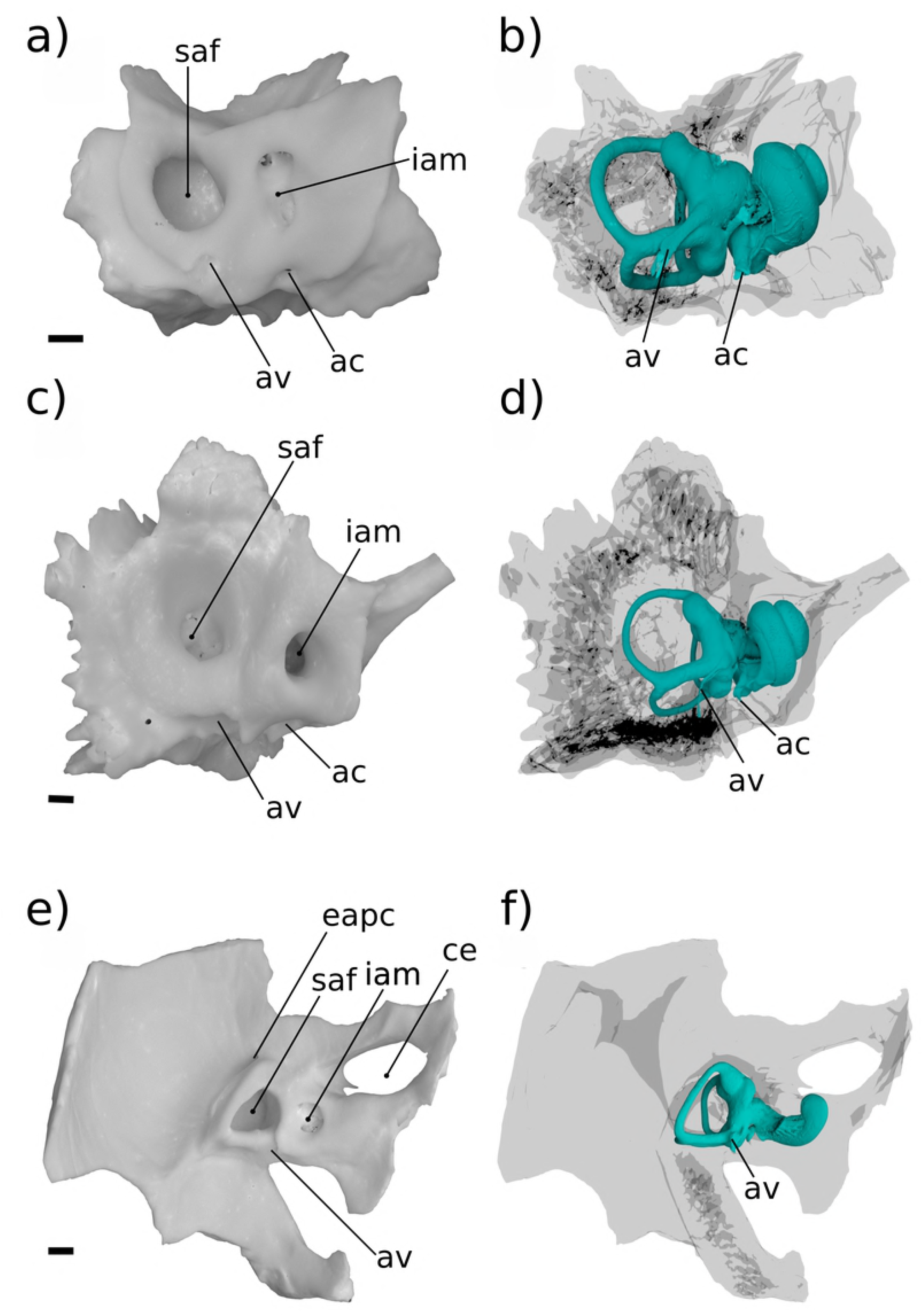
Medial view of comparative mammalian specimens. a,b *Erinaceus;* c,d *Didelphis;* e,f *Ornithorhynchus.* All specimens are left-sided, venous sinuses are not shown. In e,f a fragment of the exoccipital bone is synostosed to petrosal. Rostral is toward the right, dorsal is toward the top of the page. All scale bars are 1 mm.

**Fig 12.**
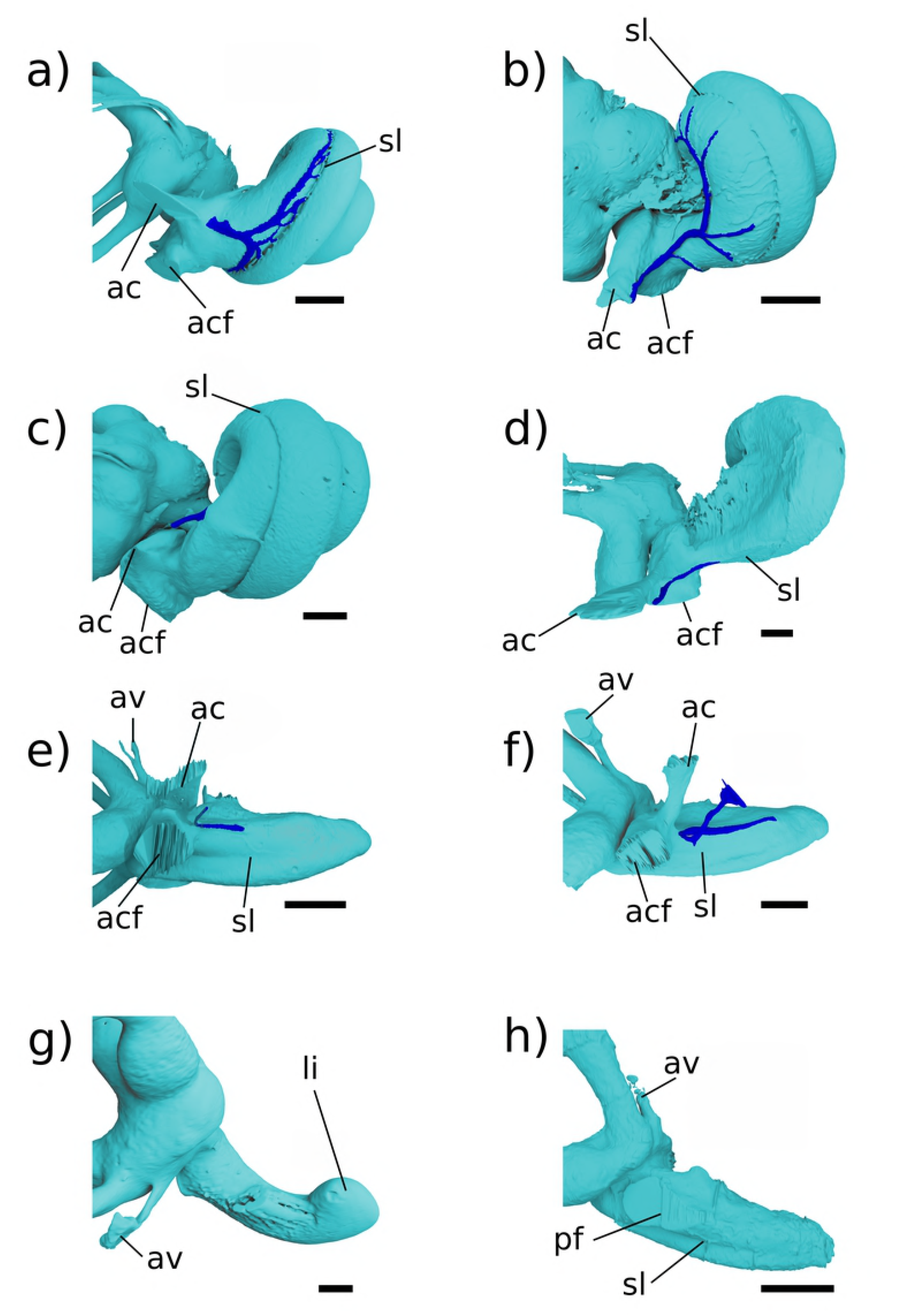
Medial view of cochlear endocast in crown mammals. All specimens are leftsided, vein of the cochlear aqueduct is shown in green. A, *Dasypus;* b, *Erinaceus;* c, *Didelphis*; d, *Vincelestes;* e, Höövör petrosal 1; f, Höövör petrosal 2; g, *Ornithorhynchus*, h, *Priacodon.* All scale bars are 1 mm.

Assuming this inference of homology is correct, the fact that the impressions of these venous structures are so prominently expressed on the endocasts of the Höövör petrosals suggests that the localized increase in venous drainage along the abneural border of the cochlea may have been a response to the increasing metabolic demands of the cochlear apparatus itself. In particular, this neomorphic drainage may be an evolutionary reaction to the increasing energetic and electrolyte requirements of the enlarging stria vascularis; a structure which in modern therian mammals contains some of the most metabolically active tissues in the body (and contains the only vascularized epithelial tissue known anywhere in mammalian anatomy). As summarized in the discussion section, the stria vascularis is responsible for generating and maintaining the highly positive endolymphatic potential in modern therians [62-64,12]. In the extant mammals in which it has been studied it is also an organ with a complex development, recruiting epithelial and connective tissue contributions from cranial ectomesenchyme and other neural crest cell populations [65].

Because the stria vascularis is a noted feature seen in all crown mammals including monotremes, the first appearance of vascular structures associated with this organ in therian ancestors are likely a consequence a the closer geometric and functional association of the bony cochlear canal with the stria vascularis and membranous cochlear duct generally (Fig 13). However, given the observations originally made by [66] of a probable endolymph-producing capillary plexus within the thickened Reissner’s membrane of *Ornithorhynchus*, the impressions of neomorphic veins draining the pars cochlearis in the Höövör petrosals may represent an osteological correlate of the “modern therian” form of cochlear endolymph production, that is accomplished solely through secretion by the highly active stria vascularis (Fig 13b-e). Conversely, the vascular branching within Reissner’s membrane seen in *Ornithorhynchus*, closely matches the position and morphology of a similar plexus in modern sauropsid amniotes (described below; [24]). Where studied, non-mammalian amniotes also lack a strong endolymphatic potential (the strength of the endolymphatic potential is still unknown in monotremes, however) and stria vascularis [67-70].

**Fig 13.**
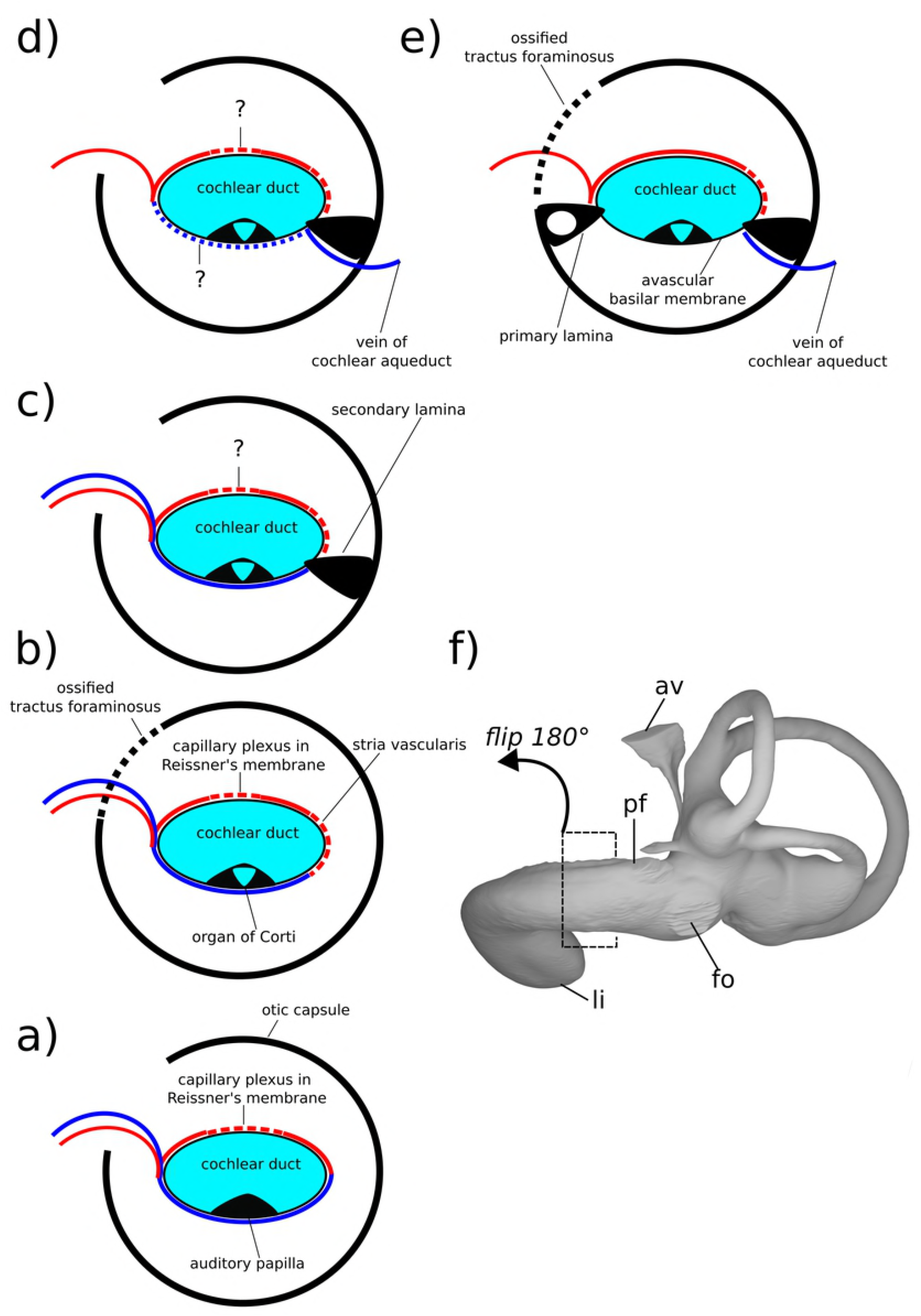
Schematic diagram showing hypothesized character states present in a cross section of the cochlear canal in early crown mammals and their fossil relatives. a, condition similar to that seen in sauropsid amniotes and hypothesized in eucynodonts; b, condition seen in modern monotremes; c, condition seen in *Priacodon* and hypothesized in early stem therian mammals; d, more derived stem therian condition seen in the Höövör petrosals; e, condition seen in modern crown therians, e.g. *Homo;* f) medial view of labyrinthine endocast of *Tachyglossus*, showing location of theoretical section diagrammed in b.

The appearance of a clear intersection of the circumpromontorial venous plexus with the endocast of the cochlear canal in the stem therian lineage (Fig 9a-d), and the localized enlargement of venous structures solely along the abneural side of the cochlear canal (the location of the stria vascularis in modern therians; Figs 12–13) in fossils otherwise showing a reduced proliferation of circumpromontorial venous sinuses, suggests that the Höövör petrosals supported a cochlear apparatus that functioned more like those found in modern therians than modern monotremes or any other vertebrate taxon [71].

As described by [42], many modern therian groups contain petrosals that are characterized by a “trisulcate” petrosal containing a prominent sulcus for the inferior petrosal sinus near the petrosal-basioccipital suture (also see criticism in [25,72] on the use of this term). In crown therians the sulcus for the inferior petrosal sinus (sulcus sini petrosi inferior; [42]) is smoothly concave and exposed endocranially, and is floored ventrally by a medially sheet-like flange of bone (crista promontorii medioventralis) which meets the basioccipital at a sutural contact in most species. In taxa with an intracranial inferior petrosal sinus (such as *Homo;* [27]), its exposure to the posterior cranial fossa is facilitated by the fact that the bony crest flooring the sulcus for the inferior petrosal sinus is much larger than the weaker ridge of bone dorsomedial to it. However, in the Mesozoic stem therians described here, and all known stem mammaliaforms, the two flanges of bone ventrally and dorsally enclosing the inferior petrosal sinus are subequally developed, and both likely contributed to the sutural contact with the basioccipital medially [44]. Additionally, in these Mesozoic forms the space accommodating the inferior petrosal sinus itself is not a smoothly and consistently surfaced sulcus, but is an elongate confluence of a highly ramified network of venous sinuses that form a substantial proportion of the total volume of the pars cochlearis.

The extent of venous excavation within the pars cochlearis has been remarked on in many prior descriptions of the petrosal morphology in Mesozoic mammalian and advanced cynodont fossils ([73,37,74,57] inter alios); and this anastomotic network together with the intramural inferior petrosal sinus has been conceptualized broadly as the circum-promontorial sinus plexus by [30]. However, only with the recent availability of high resolution micro-CT imaging (particularly [29]) has the morphology and connectivity of this venous network been sufficiently characterized so as to allow for the comparison of its discrete and homologous structures. The present report corroborates the existence of the discrete tubular “trans-promontorial sinuses” originally described by [29] (Figs 7a,c and 8a,b), and demonstrates their phylogenetic distribution outside of the clade Docodonta. The general reduction of the venous versus sensorineural contributions to the overall volume of the promontorium in successively more nested clades within mammaliaformes has also been remarked on by [74] among other sources. However, despite the relatively prolific extent of venous excavation of the pars cochlearis in stem mammaliaformes, these forms show few if any clear intersections of the circumpromontorial venous plexus with the endocast of the cochlear canal (e.g. Fig 6d). Because of the imperfect preservation of the stem therian petrosal sample used here, the precise ratio of venous to labyrinthine space within these specimens cannot be quantified; however, it is still apparent that the anatomical extent of venous proliferation in *Priacodon* is greater than that seen in either of the Höövör petrosals, matching phylogenetic expectations.

Despite having the greatest proliferation of venous structures in the pars cochlearis, the *Priacodon* specimen shows no intersection of the circumpromontorial venous plexus with the cochlear endocast (Figs 6d and 13c). This absence may ostensibly be an artifact of preservation because of the highly fractured nature of the specimen, especially along the medial surface of the petrosal where a structure such as the canal of Cotugno would be suspected. However, since *Priacodon* shows consistently large-diameter distributary branches of the inferior petrosal sinus running tangentially to the cochlear canal near the perilymphatic foramen (Fig 8a,b), and lacks branches oriented radially towards the cochlear canal or a half-pipe shaped groove for venous structures on the cochlear endocast itself, it is reasonable to suspect that the venous reservoir within the petrosal of *Priacodon* had no direct confluence with the vessels servicing the interior of the cochlear canal. This pattern of connectivity is consistent with the arrangement off venous drainage seen in modern sauropsids (and, probably, modern monotremes) where the veins draining the cochlea parallel the course of the labyrinthine artery and most likely empty endocranially into the basilar venous plexus [24,75].

In *Priacodon*, as in the docodont *Borealestes* described by [29], a large proportion of the circumpromontorial sinus plexus is formed by tubular “trans-promontorial” sinuses (Fig 14; we suggest a different terminology for these structures below). In this report, we choose to refer to these venous structures running dorsal to the cochlear canal as epipromontorial sinuses, to avoid possible unwanted connotations with the term “trans-promontorial” when used in the contest of the internal carotid artery [25]. These sinuses show no anatomical association with the contents of the cochlear canal and likely form venous anastomoses between several of the larger veins that leave the ventral braincase. In *Borealestes* (Fig 14a) the two epipromontorial sinuses both run mediolaterally within the pars cochlearis dorsal to the cochlear canal, and are termed the anterior epipromontorial sinus (“trans-promontorial sinus a” in [29]) and posterior epipromontorial sinus (“trans-promontorial sinus p” in [29]). The anterior epipromontorial sinus in *Borealestes* connects the inferior perosal sinus medially to a large venous foramen within the cavum supracochleare laterally. In [29] it is hypothesized that because of the enlarged secondary facial foramen in *Borealestes* a neomorphic continuation of the anterior epipromontorial sinus may have left the ventral cranium with the hyomandibular branch of the facial nerve. In *Priacodon* (Fig 14b,c), because of the lack of a large tubular structure connecting the inferior petrosal sinus to the cavum supracochleare, and the generally small space available for the geniculate ganglion, it is almost certain that a venous structure homologous to the anterior epipromontorial sinus did not exist (Fig 5d). However, the posterior epipromontorial sinus, which in *Borealestes* forms a confluence between the inferior petrosal sinus and the prootic sinus, does have a clear anatomical homologue in *Priacodon* based both on its position and connectivity. The posterior epipromontorial sinus in both of these taxa traverses the pars cochlearis dorsal to the cochlear canal, within the bone flooring the incipient internal acoustic meatus (Fig 14a,b). Specifically, the posterior epipromontorial sinus runs within the bar of bone separating the foramen *(Borealestes)* or foramina *(Priacodon)* transmitting the branches of the vestibular nerve from the other contents of the internal acoustic meatus (the primary facial foramen and foramen for the cochlear nerve).

**Fig 14.**
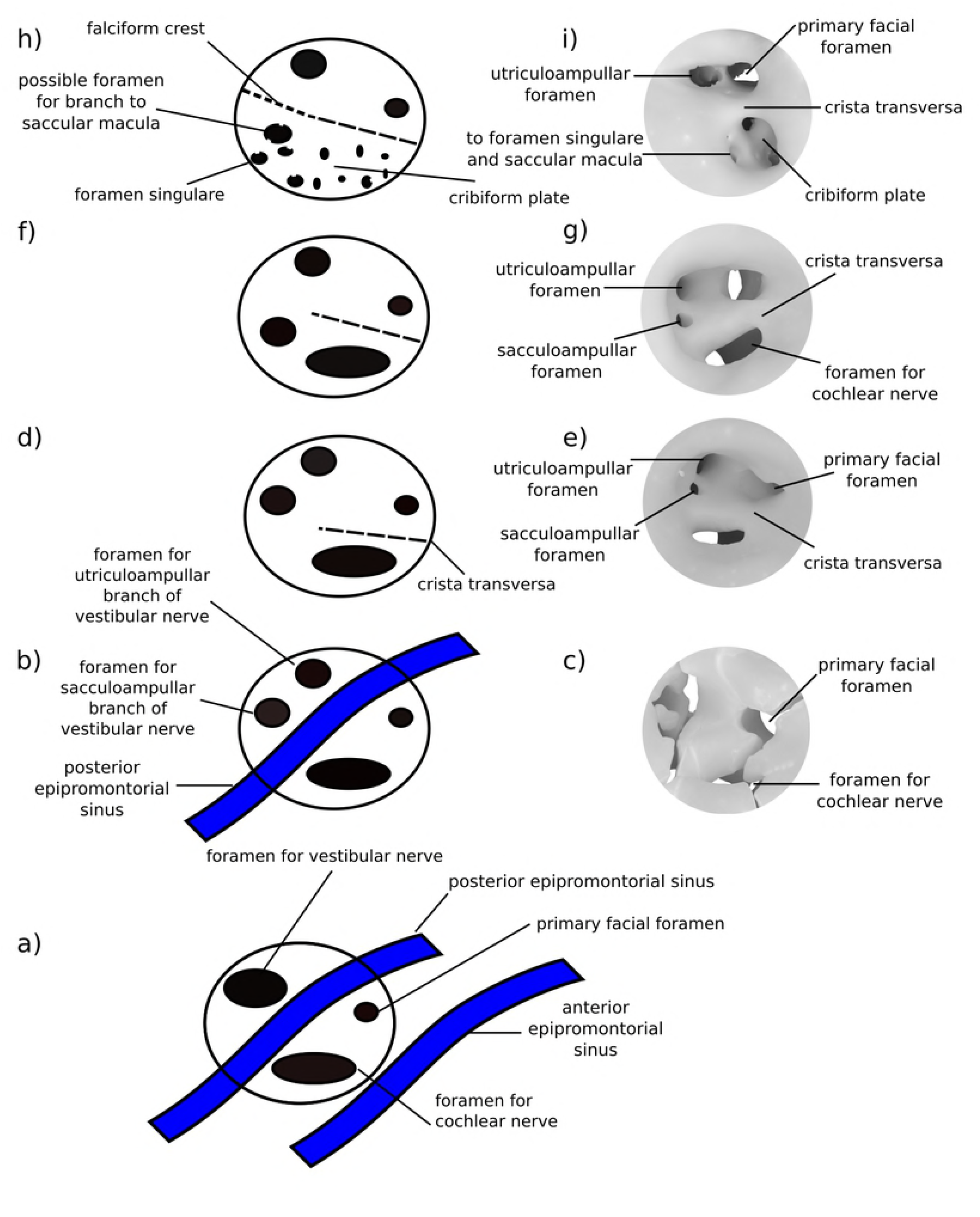
Schematic showing morphology of internal acoustic meatus in stem therians. Illustrations and renderings of left-sided internal acoustic meatus. Anterior is toward the right dorsal is toward top of page. a, condition in *Borealestes* and hypothetically all early mammaliaformes; b,c condition in *Priacodon* and hypothetically many early stem therians; d,e condition in Höövör petrosal 1; f,g condition in Höövör petrosal 2; h,i condition in extant therian mammals (e.g. *Homo).*

In *Priacodon* there is an additional “trans-promontorial sinus” running ventral to the cochlear canal, which is not seen in *Borealestes* (Figs 6d and 8b). This sinus interconnects the inferior petrosal sinus with the same aperture near the prootic canal as the posterior epipromontorial sinus. While not present in *Borealestes*, this sinus is likely a plesiomorphic feature in many mammaliaforms given its presence in *Morganucodon* (TH Pers Obs) and in the first Höövör petrosal (Figs 6b and 7a,c), i.e. taxa both closer and more phylogenetically distant to crown therians than *Priacodon.* It is here termed the hypopromontorial sinus.

The hypopromontorial sinus is the only promontorial sinus shown by H1 (Fig 6b, 7a,c, and Fig 14d,e), and there is noticeably less venous proliferation within this specimen compared with *Priacodon.* However, despite the overall reduction of venous sinuses in H1, as mentioned above, the localized hypertrophy of a neomorphic vessel (within the bony canal of Cotugno) connecting the inferior petrosal sinus with the abneural cochlear wall forms an intersection of the circumpromontorial plexus with the cochchlear endocast. This kind of intersection is not seen in *Priacodon* (Fig 6c,d) or any known stem mammaliaform taxa. Additionally, because of the longer extent of the prootic canal in H1 compared with *Priacodon*, there are a greater number of discrete venous branches stemming from the prootic canal than seen in *Priacodon*, some of which form minor contacts with the cavum supracochleare and the hypopromontorial sinus. Neither H1 or H2 show epipromontorial sinuses dorsal to the cochlear canal (Figs 2d and 5b); the close approximation of the primary facial foramen and foramina for the utriculoampullar and sacculoampullar branches of the vestibular nerve has removed the space available for these venous structures (Fig 14d-g). Finally, H1 shows a possible smooth fracture connecting the medial border of the cavum supracochleare and laterobasal extent of the cochlear canal. Whether this fracture was facilitated by a pre-existing region of highly vascularized bone is uncertain, but seems likely.

In H2 circumpromontorial venous proliferation is much less extensive than in that seen in H1 (Figs 2 and 3 versus Figs 5b and 6b), and there is no consistently wide tubular connection between the inferior petrosal sinus and prootic sinus within the pars cochleris (Figs 2b and 3b,d). This causes a greater degree of separation to exist between the venous sinuses on the medial and lateral borders of the petrosal; even though, with higher resolution micro-CT imaging, a large number of very small anostomotic connections between the prootic canal and inferior petrosal sinus do exist, these generally do not maintain a consistent direction or circumference as they project throughout the general histological fabric of the petrosal bone. As in H1, there is a discrete sinus surrounding the prootic canal, that sends out a number of small (presumably venous) interconnections to the cavum supracochleare and area around the semilunar hiatus. (Fig 2b) There is also a separate dorsoventrally oriented foramen located near the cochlear fossula and jugular notch, that has apertures on both the endocranial and tympanic sides of the petrosal (Fig 3c). This small canal is also confluent with the inferior petrosal sinus anteriorly, and most likely transmitted a small anonymous vein; however, it is conceivable that a portion of the glossopharyngeal nerve may have been enclosed by this canal as well. The posteromedial part of the cochlear fossula also shows several small (too small to be rendered in using the available micro-CT data) venous intersections with terminal tributary branches ventral to the cochlear canal. However, there are no promontorial sinuses (“trans-promontorial sinuses”; [29]) anywhere within the pars cochlearis of H2, similar to the condition in modern therians and adult monotremes (e.g. Fig 8d-f; [44]).

### Pars Canalicularis

The portions of the labyrinthine endocast infilling the pars canalicularis in the stem therians reported here are broadly similar in morphology and most linear dimensions and angles (see Table 1). Of the morphological features differentiating these endocasts, especially those existing between the two Höövör petrosals, the majority are minor differences that do not rise above the level of what is commonly intraspecific variation. The remaining distinguishing features, generally support the more derived vestibular condition of the Höövör petrosals (Fig 9; [23]).

**Table 1.** Vestibule measurements of stem therian petrosals. All distance measurements are in millimeters, angle measurements are in degrees.

|  | Höövör 1 | Höövör 2 | *Priacodon* |
| --- | --- | --- | --- |
| Straight length of cochlear canal: from posterior saccule inflection behind fenestra ovalis to tip of cochlea (mm) | 4.357 | 4.38 | 3.658 |
| Length of secondary bony lamina from saccular bulge behind fenestra ovalis (mm) | 2.566 | 2.69 |  |
| Anterior Semicircular Canal Height (ASCh SZ95) | 2.479 |  |  |
| Anterior Semicircular Canal Width (ASCw SZ95) | 2.562 |  |  |
| Length of primary common crus (mm) | 1.953 |  | 1.837 |
| Posterior Semicircular Canal Height (PSCh SZ95) | 1.73 |  | 1.272 |
| Posterior Semicircular Canal Width (PSCW SZ95) | 1.62 |  | 1.343 |
| Horizontal Semicircular Canal Height (LSCh SZ95) | 1.7 |  | 1.115 |
| Horizontal Semicircular Canal Length (LSCw SZ95) | 1.792 |  | 1.189 |
| Length of lateral expansion of cochlear canal: from front edge of the fenestra cochleae (or perilymphatic duct) (mm) | 2.151 | 1.959 |  |
| Angle Between Ant-Post SSC (degrees) | 89.29 |  | 100.47 |
| Angle Between Ant-Hoz SSC (degrees) | 73.3 |  | 79.04 |
| Angle Between Post-Hoz SSC (degrees) | 95.26 |  | 95.38 |

1075 SZ95 – Measurement from [76]

This is apparent from the thinner diameter and more exaggerated loop of the semicircular canals, especially the anterior semicircular canal, which are both longer and wider in the first Höövör petrosal than in *Priacodon* (Fig 9c-f). The anterior semicircular canal in the Höövör petrosals seem to be extended relative to the other canals by the endocranial invagination of the subarcuate fossa into the endocranial space circumscribed by the anterior semicircular canal (Figs 2c and 5a). In H2, it is also apparent that the subarcuate fossa forms a medial diversion that also projects within the circumference of the posterior semicircular canal as well. Because of damage to the posterodorsal portion of the pars canalicularis in the second Höövör petrosal (Figs 2c and 9a,b), it is not certain if the configuration of the semicircular canals is even more elongate than in H1, however it is still apparent that H2 is much more similar to H1 than *Priacodon* (Fig 9e,f). Both Höövör specimens also show a relatively more inflated bony recessus elipticus [77] for the utricular macula. Conversely *Priacodon* shows greater similarity to H1 compared to H2, extant monotremes, and *Erinaceus*, by its greater projection of the posterior apex of the recessus sphericus (for the saccular macula) caudal to the posterior wall of vestibule (Fig 9).

The canalicular endocast of H1 (and likely H2 as well) is unlike previously described Mesozoic dryolestoids [5,6,52] and more similar to *Priacodon* in that the arc of the horizontal semicircular canal is regularly circular, with their height and width being approximately equal. Strangely, the lateral (horizontal) semicircular canal in *Priacodon* is arguably more derived than those seen in the Höövör specimens because of an anteriorly projecting (centripetally pointing) conical diverticulum in the horizontal semicircular canal placed within the inner contour of the canal (Figs 8c and 9e). Additionally, the lack of a secondary common crus (a feature likely to be plesiomorphic for mammaliaformes generally; [52]) in *Priacodon* may also represent a derived trait in this taxon.

The height of the primary common crus (the bony confluence of the anterior and posterior semicircular canals) is also similar in height between *Priacodon* and H1. The aqueductus vestibuli (the bony accommodation for the endolymphatic duct) is similar in size between H2 and *Priacodon* (~0.1 mm in diameter), even though it appears relatively larger compared to other tubular structures present in *Priacodon.* The paravestibular canaliculus (the bony accommodation for the vein of the vestibular aqueduct, the main veinous drainage for the pars canalicularis, contributing blood to the sigmoid sinus in humans and probably most other amniotes; [27,28]) joins the aqueducus vestibuli along its proximal half in H1 and H2, but remains separate along its entire length in *Priacodon* where it directly intersects the dorsal surface of the vestibular endocast (Fig 9). While this is a salient difference visible in the high resolution images of stem therian endocasts used in this study, the polarity and distribution of this character are difficult to evaluate given the lack of sufficiently high resolution information for most Mesozoic petrosal specimens and the variable connectivity of the paravestibular canaliculus among extant mammals; i.e., the canaliculus can be seen to join the vestibular aqueduct proximaly in *Ornithorhynchus, Tachyglossus*, and *Erinaceus*, but remains separate to the base of the vestibular labyrinth in *Dasypus and Didelphis.*

The angle formed by the intersection the between the planes of the anterior and posterior semicircular canals is however significantly more plesiomorphic in *Priacodon* than in H1 (the only Höövör specimen for which this can be measured reliably). As reported by [78], a range of angles between 103°-157° between the planes of the anterior and posterior semicircular canals is typical of non-mammalian theriapsids, and the ~100° angle measured in *Priacodon* is much closer to this range than it is to the range of values (~90° and smaller) typifying multituberculates and extant small mammals. The ~89° angle seen in H1 is however within the range seen in extant therian mammals.

Two areas of the pars canalicularis in all three stem therians specimens show localized venous excavations (Figs 2, 3, 5 and 6). One of these sinuses, located in the posteroventral portion of the pars canalicularis and extending into the attached mastoid ossification, has been termed the paroccipital sinus by [29]. The greater extent of this structure in H1 is likely related to its better preservation in this specimen and the apomorphically large paroccipital process in this taxon (Fig 6b). A separate venous sinus arcs dorsal to the anterior semicircular canal, and is therefore termed here the subarcuate sinus (Figs 5b,d and 8e). As with other venous structures in the pars cochlearis, these venous sinuses are least developed (and may be missing altogether) in H2.

## Discussion

### Evolution of the Synapsid Ear

The estimation of neurosensory performance in any group of non-model organisms is necessarily more complex and error prone than traditional research using humans or other more-or-less cooperative animal subjects. These difficulties are especially heightened when analyzing fossil material, due to the obvious lack of relevant soft tissues and inability to work within an experimental paradigm. However, these complications do not vitiate the unique place of fossil material in the study of the nervous system [79-81]. In particular, skeletal structures such as the mammalian petrosal and middle ear show an especially close relationship with the central and peripheral nervous tissues that utilize them. The fossil record also provides otherwise unattainable information on the sequence and timing of the evolution of these tissues. As emphasize in [81], the modern mammalian nervous system is the product of many consecutive episodes of reorganization, many of which are best studied in the cranial remains of fossil synapsids. His synopsis of the stem-based clade Pan-Mammalia, inclusive of all taxa more closely related to mammals than any other living organism, by the characterization of successively more exclusive clades based around the therian crown group therefore provides a useful focus and phylogenetic context for understanding neurosensory evolution within this lineage, and the significance of stem therian petrosal structure within it.

From the earliest stretch of stem therian evolution, several extinct but derived groups appear to have diverged [33,82]. These groups include the questionably paraphyletic eutriconodonts [36,83], the multituberculates ([49,84-87]; along with their probable sister taxon Gondwannatheria [88]), and the spalacotheroid symmetrodonts [38,50]. The high-resolution micro-CT images used in this report represent the first observations of the internal structure of the otic capsule in eutriconodonts and “symmetrodont” mammals. Because of the highly derived apomorphies seen in even the earliest representatives of the multituberculate lineage, such as the presence of multiple “foramina ovalae” (apertures for the mandibular branch of the trigeminal nerve; [84]), and even more morphologically unique features found in later members of this lineage [49], the diversity of mutituberculate petrosal morphology is not discussed here.

Aside from contributing to the body of anatomical detail known for these obscure stem therian groups the petrosal observations described above allow for the broad reconstruction of auditory sensitivity, selectivity, and range in the mainline members of the stem therian lineage. These reconstructions will require a wide perspective on petrosal diversity, however, and input from several independent research programs, particularly 1) biomechanical analyses of earlier synapsid fossils [89,90]; 2) comparative and developmental studies of therian and cladotherian anatomy [22,91]; and 3) physiological research on modern mammals and non-mammalian amniotes [92,14]. As outlined below, the reconstruction of which of three non-mutually-exclusive forms of cochlear tuning, or which of four forms of sound localization, were likely present in the stem therians described here will critically depend on the findings of prior analyses of auditory physiology and vascular anatomy in non-mammalian amniotes and the construction and distribution of middle ears and their anatomically precident structures [93-96]. The following discussion therefore parallels [81] in tracing a series of sequentially more recent nodes along the backbone of synapsid phylogeny, beginning with the characterization of the amniote common ancestor and ending with the therian crown group (Fig 15). Following this synopsis, the precise implications of the osteological observations of *Priacodon* and the Höövör petrosals will be interpreted and summarized.

**Fig 15.**
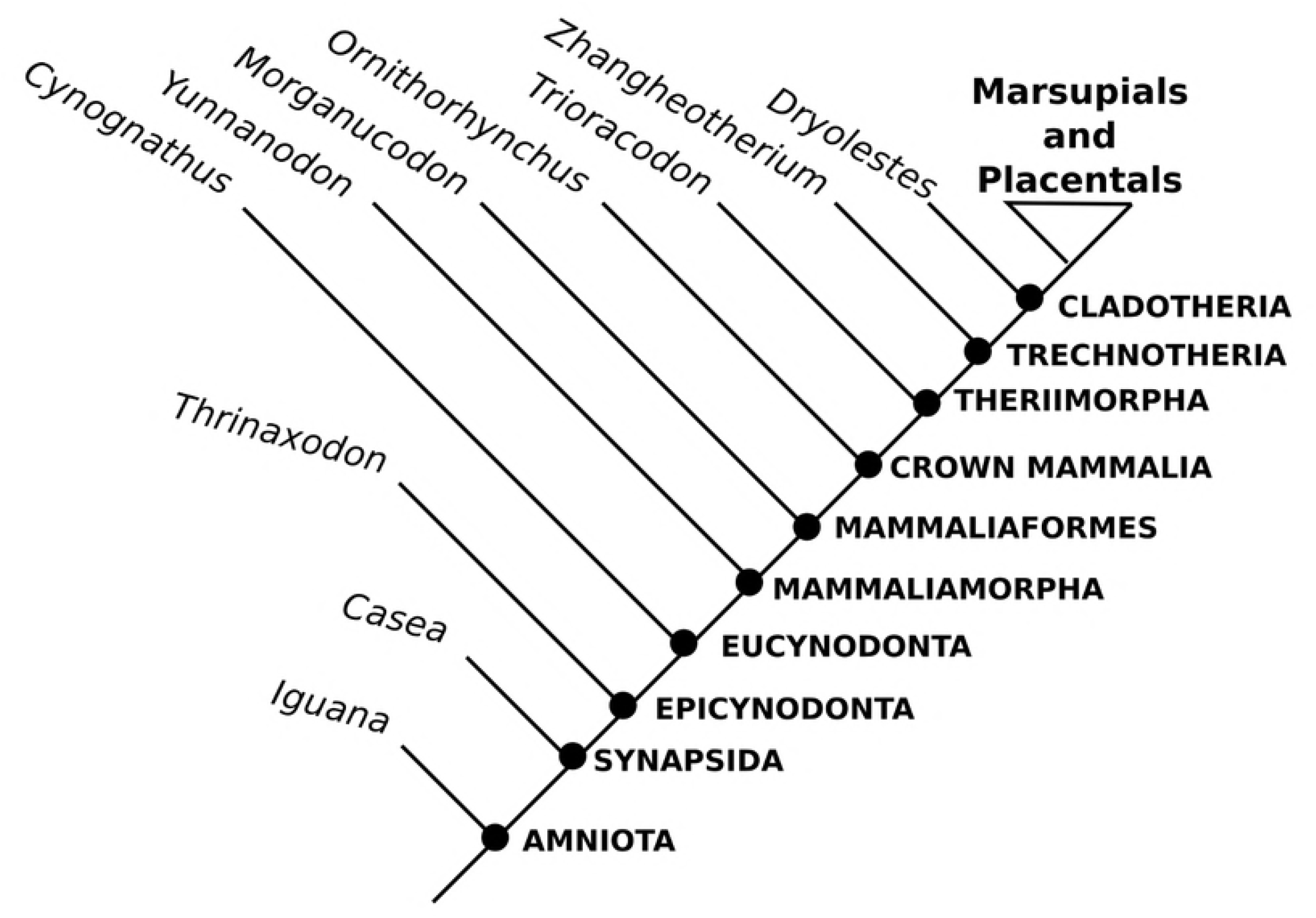
Example cladogram showing consecutively nested clades referred to in discussion.

### Amniotes

Getting biological tissue to transmit high-frequency vibrations, let alone present them to the nervous system in a way that is representative of the external environment, is an astounding technical dilemma. As such it is not surprising to find that it is one from which the first amniotes were unresponsive, for the most part. The late Paleozoic ancestors of the modern amniote lineages (sauropsida and synapsida) are reconstructed as petite “reptiles” with an estimated nine-inch snout-vent length [97]. The anteroventral subdivision of the endolymphatic labyrinth in these forms is termed the pars inferior, initially a minor diverticulum accommodating three specialized sensory epithelia [98,17]. These sensory epithelia include the incipiently subdivided saccular and lagenar maculae, and the basilar papilla (an even more recently acquired extension of the saccule; [80]). While the sensory modalities and performance parameters within which these epithelia operate are considerably more complex in fishes and amphibians, for extant and fossil members of the amniote lineage it is most likely that the saccular macula took on a dedicated role in equilibrium and proprioception, the basilar papilla an increasingly specialized role in sound perception, and the lagenar macula some combination of the two roles which became progressively more redundant thought out synapsid history [80]. The sensory epithelia contained within the pars inferior were in turn accommodated by a matching concavity within the perilymphatic (bony) labyrinth termed the sacculocochlear recess (i.e. the lagenar recess [99,98], inter alios). This configuration allowed the membrane supporting the basilar papilla to oscillate in response to vibrations induced within the endolymphatic labyrinth [100]. The reconstructed lack of a coherent gradient in width, thickness, and material properties in the early amniotic basilar membrane (a condition retained in most extant sauropsids) constrained the entire basilar membrane to oscillate homogeneously in response to the frequency content of external stimuli. In early amniotes (including early synapsids) the fenestra vestibuli was also located at the lateral or ventrolateral margin of the sacculocochlear recess.

The approximately 1 mm long basilar papilla (homolog to the mammalian organ of Corti) in the amniote common ancestor is estimated to have been competent for only a 3 or 4 octave interval of detectable frequencies; corresponding to an equally diminutive acoustic Space Constant (SC, or Space Per Octave) of ~0.3 mm per octave [101,102,103,13,14]. This short theoretical SC, together with the reconstructed consignment of these animals to the low-frequency range, is based on physiological observations of extant sauropsids *(Sphenodon* and turtles), many of which retain the most plesiomorphic form of frequency selectivity, termed electrical tuning [104].

Electrical tuning involves the active and intrinsic calibration of each individual auditory hair cell by the modulation of the density and kinetics of voltage-gated calcium channels in the basolateral cell membrane [105,106]. Calcium is critical for allowing the release of cytosolic potassium in a depolarized hair cell, and therefore its subsequent hyperpolarization. Because of the considerable refractory period in which an individual hair cell is required to import potassium from the endolymph, depolarize, import calcium from the basolateral membrane, export potassium through the basolateral membrane and thus repolarize - the intrinsic tuning of these auditory hair cells can only correspond to a dynamic range of frequencies starting from less than 100 Hz to around 1 kHz maximally in most ectotherms. At the heightened body temperatures of endotherms, theoretical considerations suggest that electrical tuning would be functional up to approximately 4 kHz [107,15].

The likely electrically tuned basilar papilla within the first amniotes was metabolically supported by both vascular and epithelial tissues running throughout the pars inferior, especially the basilar membrane (the “bottom” of the reptilian cochlear duct) and Reissner’s membrane (also termed the vestibular membrane; which forms the “top” of the cochlear duct). The tortuous capillary plexus within Reissner’s membrane in particular is the original and sole endolymph-producing organ in the ancestral amniotes, and is today retained in all extant sauropsids as well as monotremes (Fig 14a,b;[24]). The initial configuration of blood vessels supporting the auditory apparatus therefore encircled the pars inferior with little to no integration of these vessels with the surrounding skeleton of the otic capsule (Fig 13a). Indeed, one of the main observations made during dissections of the auditory apparatus of non-mammalian amniotes is that the cochlear duct can be “scooped” cleanly with dissecting tools from its cartilaginous or bony housing in the otic capsule [75,92]. This lack of bony integration is most dramatically apparent in birds, where the hypertrophied capillary plexus in Reissner’s membrane is termed the tegmentum vasculosum (e.g. in [75] fig 5 section 1; this reference also shows the source of this vasculature to be endocranial, penetrating the otic capsule in proximity with the cochlear nerve). The vestibular membrane, separating the endolymphatic labyrinth from the perilymphatic space, also supports the lagenar macula and its attached otoconial mass in extant sauropsids and monotremes [24,17]. Thus, it can be inferred that the production and composition of endolymph in modern sauropsids, with its characteristically high (~ 30 micromolar to over 1 millimolar; [70]) concentration of free calcium cations and low electrical potential, was initiated in the earliest amniote ancestors possibly as a means of facilitating the electrical tuning of the basilar papilla and stabilizing the lagena and other otoconial masses [18].

### Synapsids

Pelycosaur-grade synapsids were the first large-bodied terrestrial vertebrates to populate earth’s hinterlands. There is a consensus among modern comparative biologists, based on osteological and modern comparative considerations, that forms in this early burst of synapsid evolution lacked any form of tympanic membrane or acoustically relevant cranial air-spaces [92,108]. Additionally, the co-option of the hypertrophied stapes (a second arch viscerocranial element) as a firm structural interconnection between the posterior neurocranial and dermatocranial components of the skull, acted as a mechanism for reinforcing the ancestral jaw apparatus against structural deflections during biting. This probably also limited its auditory sensitivity [109,110,92].

What acoustic receptivity these forms acheived was likely mediated through a mixture of indirect conduction (where the majority of the dermatocranium acted as a vibrational antenna, possibly in service of a semiaquatic lifestyle in many taxa) and direct conduction (where seismic/airborne vibrations are transmitted through the lower jaw [90,111]. Waves of pressure were therefore transmitted into the otic capsule through the skull’s out-of-phase motion with respect to the massive and inertially stabilized stapes [108,56]. These conduction mechanisms are most appropriate for low-frequency and high-amplitude vibrations and electrical tuning would have been more than capable of processing these signals.

Opposite the condition of its lateral border, the endocranial wall of the otic capsule is poorly ossified in pelycosaurs and most other non-mammaliaform synapsids. This is reflected by the absence of an invaginated internal acoustic meatus and lack of ossified partitions separating the perilymphatic foramen and cranial nerves 7-12 along the medial aspect of the braincase [74]. What can be determined about the structure of the otic capsule shows that the pars cochlearis is absent in the earliest synapsids; however, the corresponding pars inferior of the endolymphatic labyrinth would have been accommodated by a shallow sacculocochlear recess that itself did not emarginate the wider contour of the perilymphatic labyrinth [100]. The lack of a pars cochlearis within the early synapsid otic capsule does not imply a more or less monolithic composition of the periotic skeleton, however; and the adult otic capsule in early synapsids is a composite structure mainly formed by the prootic and opisthotic bones, with variable minor contributions from other cranial elements.

By the Middle Permian a diverse group of therapsids had evolved from the carnivorous sphenacodontan pelycosaurs. These animals display a trend of progressive loosening of the quadrojugal and quadrate from the squamosal, possibly resulting in some combination of incipient streptostyly and whole-bone sound conduction. Additionally, the increased development of the reflected lamina of the angular bone (homolog of the mammalian ectotympanic) and formation of the recessus mandibularis as a mandibular resonating chamber, suggest that direct conduction of low-frequency, possibly airborne, sound was possible. The relatively voluminous recessus mandibularis in some of the larger therapsids point to its possible dual function as a resonator for both the reception and production of airborne vocal signals [110]. With the increased development of the recessus mandibularis, a trend of decreasing reliance on sound conduction via intra-bone vibrations and greater reliance on conduction through whole-bone vibration is also apparent [108]. However, the low transformation ratios between the area of the reflected lamina of the angular and the fenestra ovalis would have provided little compensation for the energy lost due to impedance mismatch between the surrounding air and fluids inside the otic capsule [89,110].

### Epicynodonts

Near the Permo-Triassic boundary the epicynodonts, along with other therapsid groups, show the complete subdivision of the oronasal cavity into dedicated oral and nasal cavities by the completion of an osseous secondary palate [112,113]. The resultant caudal aperture of the nasal cavity, the internal choanae, created a new communication to the rostral pharynx (nasopharynx). The nasopharynx itself became a hub for gaseous communication to the larynx caudally, and its lateral apertures are also reconstructed as having communicated with the recessus mandibularis [114].

The epicynodonts also show the first subdivision of the sacculocochlear recess of the perilymphatic labyrinth into a dedicated saccular recess (recessus sphericus) and a separate bony cul-de-sac termed the cochlear recess [100]. This morphology of the perilymphatic labyrinth suggests that the corresponding pars inferior of the endolymphatic labyrinth was also at least incipiently subdivided into a discrete saccule (confluent at one end to the endolymphatic duct and utricle, and the cochlear duct at the opposite end) and cochlear duct. In this transitional cochlear organization (as demonstrated by *Thrinaxodon;* [115]) the rostral border of the ventromedially pointing cochlear recess within the prootic bone does not extend anterior to the rostral border of the fenestra ovalis. As such, these forms lack a true pars cochlearis of the prootic bone [100].

### Eucynodonts

Eucynodonts show a progressive trend of increasing relative size and differentiation of the jaw adductor musculature, while simultaneously reducing the size, strength, and cranial integration of the bony jaw apparatus itself. This counterintuitive development was permitted by the reorientation of adductor leverage to minimize the reaction force experienced by the quadrate-articular joint and postdentary elements [116]. Other morphological features seen in this taxon include the change from a plate-like reflected lamina to a rod-like reflected lamina of the angular bone, creating a laterally facing gap between the reflected lamina and main body of the angular, that was almost certainly spanned in life by an incipient tympanic membrane [112]. The remaining postdentary bones were synostosed into a gracile but rigid postdentary rod that remained attached to the angular. As the postdentary rod, the postdentary bones lost their sutural connections with the dentary bone but remained appressed to the dentary within a smoothly concave postdentary trough. The rostrocaudal orientation of the postdentary rod, with its flexible posterior connection to the squamosal, and lack of strong rostral attachments to the dentary, allowed the postdentary rod to function as a first-class lever with a longitudinally oriented fulcral axis. This provided a mechanical linkage for transmitting airborne vibrations impinging on the angular tympanic membrane to the quadrate and stapes medially [108]. The air-filled space medial to the tympanic membrane (presumably a vestige of the recessus mandibularis), combined with the approximately 1/30 area ratio between the fenestra ovalis and angular tympanic membrane [90], would create a precursory condition to the middle ear seen in the later and most advanced cynodonts. However, the existence of a true cavum tympani medial to the angular tympanic membrane and the usefulness of this apparatus for frequencies above 2 kHz is doubtful [90], as the estimated sum mass of the postdentary rod would have been over ten times greater than the homologous elements in a similarly sized therian mammal. The eucynodonts also show the first appearance of a true pars cochlearis. This neomorphic region is defined in [19] (also see [10,100]) as the portion of the otic capsule accommodating the saccule and cochlear duct, that were likely present as discrete structures in epicynodonts. The weight of morphological evidence suggests that while not within crown mammalian performance levels, the auditory capacities of the early eucynodonts were heightened beyond anything preceding them in synapsid history [108].

### Mammaliamorphs

The most advanced group of cynodonts are the mammaliamorphs, that include the true mammaliaforms and several clades showing incredible morphological convergences with them [117]. This group also includes the only synapsid taxa to survive beyond the Early Cretaceous. These forms show many significant apomorphies related to the function of their inner ear and newly acquired middle ear - including the consolidation of the prootic and opisthotic into the true petrosal bone [112,100], the segregation of the perilymphatic foramen from the jugular foramen, and separation of the foramina for the cochlear and vestibular branches of the vestibulocochlear nerve [118]. However, far and away the most lauded morphological characteristics seen in this group are the articulations between the dentary and squamosal bone, and development of a true cavum tympani (the MME character state; [56,77]. The development of a tympanic middle ear in synapsid ancestry is an important event with functional implications regarding the sensitivity and maximal frequency attained by these and later forms; however, the accurate interpretation of these features, especially the contents of the otic capsule, rely on comparative inferences derived from the study of the tympanic middle ears in fossils and extant non-mammalian amniotes. It should therefore be reiterated here that, (for anatomical and developmental criteria skillfully outlined in [20,119,96] inter alios) the best interpretation of the available evidence suggests that the acoustically significant intracranial air spaces developed in advanced synapsids (including the MEC and its later derivations) are not homologous with tympanic cavities developed in many other tetrapod lineages [92]. The initial form of the synapsid cavum tympani (associated with the MME) likely displayed a broadly confluent relationship with the nasopharynx. The medial (proximal) subdivision of the cavum tympani was therefore not separated from the nasopharynx by an extended auditory tube and likely allowed free motion of air between the lateral portion of the tympanic cavity to the nasopharynx and onward across to the contralateral cavum tympani. Modern *Ornithorhynchus* also shows this condition [51].

An unconstricted (if tortuous) air-filled passageway connecting both ears is hypothesized as a mechanism for sound localization in advanced cynodonts and other, more plesiomorphic synapsids [120]. In the majority of extant sauropsids the sole method of sound localization in the horizontal (azimuthal) plane is by the simultaneous use of the ipsilateral and contralateral ears, and the air-mass medially connecting both, as a pressure-gradient receiver [121,122,92,31,123]. With the successive medial-posterior-ventral relocation of the articular and quadrate relative to the newly formed dentary squamosal contact, open contact between both ears likely became progressively more constricted, requiring the development of uniquely mammalian mechanisms of sound localization discussed below [122,120].

The degree of ossification of the otic capsule in Mammaliamorpha is also significantly greater than in other known cynodonts, with two discrete foramina for the vestibular and cochlear branches of the vestibulocochlear nerve, respectively. These are recognizable in the medial wall of the petrosal even in early forms [78,77], and in the most plesiomorphic condition they are of subequal diameter. These forms are also the first to show an appreciable amount of “waisting” between the components of the vestibular endocast contributed by the pars cochlearis and pars canalicularis. The development of a more or less prominent crista vestibuli also separates the areas of attachment for the utricular and saccular maculae, forming discrete vestibular recesses to accommodate these structures (the recessus elipticus and recessus sphericus, respectively; [22,77]).

### Mammaliaformes

Having inherited the MME from advanced mammaliamorph cynodonts, the Mammaliaformes reinforce the new dermatocranial jaw articulation by developing a true mandibular condyle on the dentary, matching an equally developed glenoid fossa on the squamosal [117, 33]. However, with formation of the fossa incudis on the petrosal and redistribution of masticatory musculature onto the dentary bone, the placement and size of the ancestral jaw articulation became more directly under the influence of the auditory mechanism, while the dentary and squamosal became increasingly specialized for masticatory purposes [109, 116, 33].

Along with the increasing specialization of the articular and quadrate for acoustic purposes, a wider reorganization of external and internal aspects of the petrosal can also be seen in early mammaliaform fossils. Externally, the enlargement of the pars cochlearis at the expense of the basisphenoid and other midline structures, loss of a thickened rim around the fenestra ovalis, and the separation of the hypoglossal and jugular foramina [74, 33] can be seen in early members of the clade. These features are possibly related to the insulation and stabilization of structures surrounding otic capsule, or in the case of the reduced rim of the fenestra vestibuli to facilitate a greater range of motion between the stapes and petrosal (i.e. a “rocking”-type motion between the stapes and petrosal, in addition to a pistion like motion; [20]). More significantly, these mammaliaformes also show the allometric enlargement and ventral inflation of the pars cochlearis of the petrosal relative to other advanced cynodonts, forming a variably flattened or convex promontorium on its newly exposed ventral surface. As mentioned above, a promontorial structure has also been observed in juvenile non-mammaliaform probainognathians [124], and as [74] observed, at the time of its initial appearance the majority of the increased volume in the pars cochlearis is not immediately utilized by the elongated but relatively small cochlear canal. Instead, the persistence of the promontorium into adult stages in mammaliaformes and other mammalimorph taxa may solely be a product of paedomorphosis, as in these early forms most of the volume of the promontorium is taken up by a complex network of venous canals and sinuses termed the circumpromontorial venous plexus (CVP; [30,37,125]).

In many mammaliaformes the CVP includes two sets of venous canals dorsal and ventral to the cochlear canal accommodating the epipromontorial and hypopromontorial sinuses, that communicate with the inferior petrosal sinus (Fig 14a). The inferior petrosal sinus is located in an intramural location between the petrosal and basisphenoid in these early taxa. The ventral bulging of the pars cochlearis relative to the cranial base also variably impinged the stapedial artery, forming an impression along the margins of the fenestra ovalis in *Morganucodon* and more advanced forms [118]. Endocranially, earliest mammalaiformes still lacked an excavated internal acoustic meatus, with the small prefacial commissure showing little association with the course of the vestibulocochlear nerve (Fig 14a; [30]). As described above it is also very likely that a posterior epipromontorial sinus, running between the inferior petrosal sinus and prootic sinus, was present within the bone separating the primary facial foramen and the foramen for the vestibular branch of the vestibulocochlear nerve (Fig 14a,b). Other significant features of the pars cochlearis are internal, such as the initiation of the abneural curvature (concave toward the side of insertion of the cochlear nerve) of the cochlear canal and, as seen in *Morganucodon* and more derived stem mammaliaformes, the terminal inflation of the cochlear canal for the accommodation of the lagenar macula [39].

The reorganization of the ancestral amniote depressor mandibulae muscle to the stem mammalian levator hyoideus, and ultimately the modern stapedius muscle in seen in therians, is plausibly also a mechanism for preferentially filtering high-frequency vibrations within the ossicular apparatus by raising the reactance (inertial resistance to sound conduction) disproportionately for lower frequency sounds [126, 20,48,95].

### Mammalia

The characterization of the crown mammalian ear solely from the morphology of extant taxa would provide a markedly distorted reconstruction of the mammalian common ancestor compared to what modern paleontological and developmental research now suggests. This is because of the over 150 million years of independent transformation seen in both living mammalian lineages (monotremes and therians), making extant mammals an unrepresentative sample of mammalian diversity as a whole; and because of the widespread and detailed homoplasy often seen in the evolution of petrosal structures [33]. Based on the distribution of characteristics within extant mammals alone, one would reasonably conclude that the condition of the crown mammalian common ancestor consisted of 1) a Detached Middle Ear (DME), with auditory ossicles fully independent from the lower jaw apparatus, and 2) an osseus armature for the distribution of the cochlear nerve axons termed the tractus foraminosus (creating the perforated cribiform plate within the internal acoustic meatus). Fossil evidence clearly shows these traits to be homoplastic ([127, 56], also see [128]).

The base of the secondary bony lamina is variably present in monotremes and now seen in a more elongate state in *Priacodon*, suggesting that some development of this structure may have been a derived feature of mammals generally (Figs 12g,h and 13b,c; although according to [17] in monotremes the base of the secondary lamina does not contact the basilar membrane). Of greater significance for the reconstruction of auditory capacities in early mammals are soft-tissue characteristics, many of which can only be inferred based on their distribution in extant taxa only. These considerations suggest that the mammalian ancestor contained a secondary tympanic membrane (a membrane interface between the airspace of the cavum tympani and liquid-filled subarachnoid space, [51]), that was not at this stage suspended by a bony armature (such as the round window that formed in later trechnotherian mammals). The status of the secondary tympanic membrane as an apomorphy or symplesiomorphy in crown mammals is unclear. Several less ambiguous soft-tissue symplesiomorphies in the mammalian ancestor, however, are the retention of the lagenar macula and the vascular plexus within Reissner’s membrane (as is still seen in monotremes today; [66]). These soft-issue components of the cochlear duct were complemented by the presence of the stria vascularis, a specialized endolymph secreting organ (Fig 14b-e), that developed within the synapsid lineage sometime before the divergence of the modern mammalian clades.

Most importantly, the mammalian common ancestor can be reliably inferred to have attained a true organ of Corti, the characteristically mammalian homolog of the amniote basilar papilla. The organ of Corti is diagnosed by the functional differentiation of two subgroups of auditory hair cells along an axis perpendicular to the axis of tonotopy [129,16]. This division of labor between less derived and more afferently innervated Inner Hair Cells (IHCs) on the neural side of the basilar membrane, and more specialized and efferently innervated Outer Hair Cells (OHCs) on the abneural side of the basilar membrane, is a fundamental component macromechanical tuning. In therian mammals the functional specialization of OHCs to amplify weak (i.e. quiet) or dampened pressure waves propagating along the cochlear canal is implemented through the action of an apomorphic reverse transduction mechanism [105, 12]. Possibly correlated with this adaptation is the lack of evidence for significant contributions from either electrical or micromechanical tuning in mammalian cochlear function ([13]; although for evidence of a possible minor role of micromechanical active processes in mammals see [130,131].

The segregation of two subpopulations of auditory hair cells is delimited morphologically by a patent tunnel of Corti running longitudinally within the endorgan, and specialized populations of non-sensory supporting cells within the cochlear epithelium. Additionally, the mammalian common ancestor seems to have also defined the allometric size of the combined auditory hair cell population, with extant monotremes and similarly sized therian mammals (dogs, cats, etc.) having a similar total number of these cells [132]. However, the relatively much more elongate and organized organ of Corti seen in modern therians creates a thinner and longer distribution of the auditory hair cells, compared to monotremes.

It is unclear how innovative these soft tissue characteristics are to the mammalian clade itself, and the organ of Corti and stria vascularis in particular are likely to have appeared at an earlier point in synapsid phylogeny, within Mammaliaformes if not earlier. It is therefore also unclear if the petrosal of the earliest crown mammals would be diagnostically recognizable from the petrosals seen in other early mammaliaforms such as *Hadrocodium* and *Morganucodon* (see [133] for two examples of this frustrating ambiguity). Finally, it is also unknown if the uniquely mammalian external ear, with its characteristic pinna structure, would have appeared within the crown mammalian ancestor, within other advanced synapsid taxa, or at some later point solely within the stem therian lineage. The soft-tissue evidence for an involuted cartilaginous pinna in *Tachyglossus* reported by [134] is difficult to homologize with extant therian structures. Otherwise the phylogenetically deepest evidence for the presence of external pinnae is provided by *Spinolestes*, an excellently preserved eutriconodont [135], and member of the diverse stem therian clade Theriimorpha.

### Theriimorphans

The monophyletic group containing the carnivorous eutriconodonts, the more derived “acute symmetrodonts,” and therian mammals – Theriimorpha is the most inclusive well-supported group within the therian lineage [117, 56]. Despite the relative abundance of excellently preserved fossil remains referable to early members of this group (such as *Spinolestes* mentioned above), several authors have mentioned the relative lack of knowledge on the internal petrosal morphology in this clade (e.g. [68, 58]). However, the prior descriptions of external petrosal morphology in eutriconodonts [36, 3], combined with the observations of the internal petrosal anatomy of *Priacodon* outlined above, suggest that basal theriimorphans can be characterized by a similar but slightly straighter morphology of the cochlear canal than the earliest mammaliaformes. This is reflected by the steeper lateral aspect of the promontorium in ventral view, and lack of an apparent apical inflation for the lagenar macula within the labyrinthine endocast (although in *Priacodon* the cochlear canal remains relatively untapered and thick to its apex).

The cochlear endocast of *Priacodon* (Fig 9e,f) also shows the first appearance of a elongate and projecting secondary lamina in early theriimorphs. In extant therians the secondary lamina is associated with tonotopic locations of the cochlea dedicated to hearing above 10 kHz [136, 14]. However, at the time of its first definite appearance in the theriimorph taxa described here, its functional significance is much less certain. Given that the secondary lamina appears without an opposing primary bony lamina or even tractus foraminosus to provide an opposing attachment for the basilar membrane, it seems unlikely that a significant magnitude of tensile force would be able to be transmitted across the scala tympani side of the cochlear duct. Additionally, as the variably present “base of the secondary lamina” in extant monotremes has been observed to lack a direct connection (histological adherence) with the cochlear duct when present [17], it is conceivable that the (much longer and wider) secondary lamina seen in *Priacodon* and the Höövör similarly lacked an association with the cochlear duct. For reasons outlined in the next section, this seems like an improbable scenario, especially for the Höövör specimens. However, even in *Priacodon*, arguably the most plesiomorphic stem therian for which internal petrosal structure is known, it seems likely that the large secondary lamina is a true homolog of the secondary lamina seen in Mesozoic cladotherians and modern therians. For example, the secondary bony lamina in *Priacodon* runs longitudinally onto the crista interfenestralis, that separates the fenstra ovalis and perilymphatic foramen (Figs 6c and 9e,f). This is essentially the same positional relationship seen in later cladotherian petrosals [136], suggesting that some association of the abneural side wall of the cochlear canal and the cochlear duct had begun to form even at this early stage in the ancestral therian lineage. Thus, while its role in the generation and transmission of tension across the basilar membrane may not have been as functional as in modern therians, the appearance of the large secondary lamina may still have served as a mechanism to better match the stiffness between middle and inner ears [14], or as a means of stabilizing the position of the cochlear duct with respect to the larger cochlear canal.

The status of the middle ear on the other hand can be relatively well characterized in the theriimorph common ancestor, and has been confidently reconstructed as having attained the PME character state [56]. Interestingly, many members of this clade show the further calcification of Meckel’s element which forms the structural attachment between the malleus and dentary; a peramorphosis possibly related to the continued reliance on the detection of seismic sound sources through direct conduction even after the mediolateral separation of the postdentary bones from the medial surface of the lower jaw [56]. The fortunate conversion of Meckel’s element into a fossilizable structure therefore provides some possible evidence for the use of non-tympanic low-frequency sound conduction even after the attainment of a true middle ear and probable stenosis and elongation of the interaural canal. However, [96] have advanced arguments for why this element may not be homologous with the embryonic Meckel’s cartilage: if so some of our consideration here would need to be nuanced, but the evidence for non-tympanic conduction is still defensible.

### Trechnotheres

The clade Trechnotheria includes “northern tribosphenic” taxa (crown therians and close relatives), the “acute symmetrodonts” (spalacotheroids and amphidontids), and various “pre-tribosphenic” taxa (such as *Vincelestes* seen in Fig 12d, and dryolestoids from the Northern and Southern hemispheres) [33]. While this group is known from a considerable diversity of dental and postcranial remains, the description of Höövör petrosal 1 provided by [2] represents the first tentative information on the external petrosal anatomy in either a basal trechnothere or a closely related stem theriimorphan (i.e. a tinodontid symmetrodont or a gobiconodontid). The descriptions of the second Höövör petrosal provided here demonstrate several more derived features of H2, making it more likely (but still not certain) that this specimen is referable to Trechnotheria. However, because of the limited amount of information on mammals with this kind of petrosal organization, and possible close relatives of trechnotheria (especially gobiconodontids) showing broadly similar features, the following characterization of the trechnotherian common ancestor can only be tentative at best.

Probably the most salient feature (if not synapomorphy) seen in the Höövör petrosals and other more confidently referred trechnotherians, is the formation of a true fenestra cochleae (round window) by the bony subdivision of the ancestral perilymphatic foramen (Figs 2a and 6a). This character has uncertain mechanical implications for the isolation and function of the cochlear apparatus. This process is a neomorphic bony strut that bifurcates the ancestral perilymphatic foramen, forming the elongate cochlear aqueduct (containing the membranous perilymphatic duct) medially and fenestra cochleae laterally (i.e. Fig 7e; [46, 47]). The enclosure of the perilymphatic duct in a bony canal is also seen in some multituberculates and late adult tachyglossids [44]; in addition to several groups of squamates and archosaurs [137]. In addition to the widely homoplastic distribution of an enclosed perilymphatic canal (aqueductus cochleae), reference to the fenestra cochleae as a “window” may also be misleading in that the metaphorical “glass” of the round window is actually the secondary tympanic membrane (separating the airspace of the cavum tympani from an endocranial cerebrospinal fluid filled space). This structure has been reconstructed as being present in the mammalian common ancestor, if not the ancestor of all amniotes [51, 137]. Therefore, only about one quarter of the bony “frame” of the round window can be considered an apomorphy, giving it a secondary and more ambiguous functional significance. The lack of exposure of a relatively long perilymphatic duct into the middle ear cavity may be considered perhaps a more significant transformation, with the formation of a fenestra rotunda being a byproduct of the progressive inclusion of the perilymphatic duct into the osseous perilymphatic canal.

The most plesiomorphic trechnotherians [2, 38, 50] also show features related to the increased ventral extension of the pars cochlearis relative to the surrounding pars canalicularis and other cranial elements. These include the more vertical orientation of the crista interfenestralis which, as in the Höövör petrosals described above, terminates caudal to the promontorium without contacting the mastoid area or paroccipital process. The area immediately caudal to the promontorium within the tympanic aspect of the pars canalicularis also forms a post-promontorial tympanic sinus, that is broadly confluent laterally with the lateral trough [44]. In trechnotheres the post-promontorial tympanic sinus is possibly a j mechanism for increasing the volume of the cavum tympani and thereby reducing the effective rigidity of the (primary) tympanic membrane [20]; and/or represents a bony accommodation for the lateral head vein [26]. As also seen in monotremes and multituberculates, the earliest trechnotherians also reduce or lose a robust quadrate ramus of the alisphenoid (epipterygoid), while retaining a thin lateral flange of the petrosal (the ancestral attachment of the quadrate ramus); and similar to several eutriconodontans, one early spalacothere shows a calcified Meckel’s element [138].

### Cladotheres

A significant characteristic of this clade is the dorsoventral coiling of the cochlear canal, above and beyond the initial lateral (abneural) curvature of the cochlear canal developed in early mammaliaforms [39]. This reconfiguration of the cochlear apparatus is expressed on the ventral surface of the pars cochlearis as the projecting bulbous morphology of the tympanic surface of the promontorium. The cladotherian common ancestor likely also attained a transpromontorial course of the internal carotid artery, as is evidenced by the sulcus left by the impingement of the expanding pars cochlearis with this vessel.

Ventral bulging is likely not the result of volume restrictions within the pars cochlearis, due to the variably incomplete filling of the bony space available within the pars cochlearis by the cochlear canal [6, 52]. However, limitations in the relative dorsoventral linear distance available to the cochlear canal within the pars cochlearis may be responsible for the evolutionary timing of the dorsoventral coiling, as the images described above demonstrate that the progressive reduction and loss of the epipromontorial and hypopromontorial sinuses is a trend taken to completion in the stem therian circumpromontorial venous plexus before the initiation of dorsovental coiling in cladotheres.

While all previously described stem cladotherian petrosals from northern continents (i.e., dryolestoids [5,39,6,52] and possibly [133]) and the pre-tribosphenic mammal *Vincelestes* (Fig 12d; [37]) show at most 270° of dorsoventral coiling, cladotherian petrosals from Argentina [139,140,4] demonstrate that complete (360° and beyond) cochlear coiling was attained by later Mesozoic and Cenozoic stem cladotherian lineages in parallel to crown therians.

In Northern Hemisphere dryolestoids, complete cochlear coiling is not known to occur; however, the cochlear endocasts of *Henkelotherium* [5] and *Dryloestes* [6] demonstrate that the suspension of the basilar membrane between a true bony primary lamina and secondary lamina had developed in these forms, also possibly in parallel to crown therians. In dryolestoids and therians, this increased contact between the endolymphatic cochlear duct and perilymphatic cochlear canal is thought to provide better impedance matching between the middle and inner ears [11], that may have allowed for adaptive increases in maximal detectable frequency. However, theoretical concerns outlined in [71,141] suggest that the gain in selectivity and sensitivity by the cochlear apparatus from these osteological features would be mainly limited to frequencies less than 20 kHz in Mesozoic taxa, where a complete primary lamina is only present basally. The variable extent of cochlear coiling, and presence/absence of bony laminae, make the restructuring of the cochlear canal difficult to interpret. However, all cladotherian taxa invariably show the presence of a neomorphic bony armature, termed the tractus foraminosus, for the individual distribution of cochlear nerve axons and supporting tissues through the foramen acusticum inferius ([52]; although the presence of a tractus foraminosus is ambiguous in the specimen described by these authors). The presence of the foramen acousticum inferius itself, composed of the foramina for the cochlear and sacculoampullar branches of the vestibulocochlear nerve and their derivative and supporting structures, may also be a neomorphic feature of Cladotheria, or cladotheres and their closest trechnotherian relatives (Fig 14f-i).

### Theria

With the advent of the crown clade Theria, mammals attained the most sophisticated form of airborne sound detection known among terrestrial vertebrates. Counterintuitively, members of this clade to which our human species belongs are typically characterized by the capacity for the detection of ultrasonic frequencies (defined as frequencies above human reception). These capacities are facilitated by the coordinated development of many significant osteological characters that have been widely commented on in the paleontological literature [142,23, 58, 56]. Prominent among these are the attainment of a DME, and the completion of at least one full whorl by the dorsoventrally coiled cochlear canal. The level of cochlear coiling in therians is also more derived than that seen in most other cladotheres because of the presence of a modiolar structure (bony armature for the spiraled cochlear nerve) within the concavity defined by the coiled cochlear canal. This morphology causes the peripheral processes of auditory afferent neurons to take on a radial, as opposed to parallel, distribution [142]. However, despite the diverse literature on these and other osteological apomorphies, displayed by all or particular subsets of crown Theria, precise functional implications of the bony structures of the therian middle and inner ear remain ambiguous.

Because of their representation by the majority of extant mammals, soft-tissue characteristics within the earliest crown therians can also be reasonably estimated. For instance, unique vascular features that can be inferred to have been present in the petrosal of the therian common ancestor, if not earlier, include the almost-completely intracranial course of the superior ramus of the stapedial artery and the loss of the post-trigeminal vein [10,26]. The progressive loss of the bony canal for the prootic sinus, and reduction of the lateral head vein are also an apparent trend in plesiomorphic members of both major therian clades, Eutheria and Metatheria (Figs 10a-d and 11a-d; [10,26,58]). More pertinent for the reconstruction of auditory sensitivity in early therians is their exclusive reliance on the stria vascularis for the production of endolymph (Fig 13e), and their uniquely high-positive endocochlear potential [63,70]. A recent publication [18] hypothesizes that these and other distinguishing features of the therian cochlear apparatus are related to the evolutionary compensation required by the loss of the lagenar macula, yet another unique feature seen in the therian inner ear. As detailed in the next section, the observations of stem therian petrosals described here can be seen as broadly concordant with this hypothesis, and can ostensibly provide boundaries for the reconstruction of auditory performance in the first crown therians and their earlier ancestors.

Concurrent with the evolutionary development of the most sophisticated and high-frequency sensitive ear known among tetrapods, crown therians also sever the last vibrational interconnections mediolaterally linking both ears (if they in fact existed) by the elongation and stenosis of the lateral parts of the ancestral interaural canal [51,110,128], forming a muscular and facultatively patent eustachian tube. The implications of this increased separation for the reconstruction of sound localization and central processing capacities in early therians are discussed below; however, fossilizable evidence of this trend in interaural isolation is shown by the widespread and homoplastic development of ossified bullae within many crown therian lineages ([143]; for evidence of convergent and functionally analogous “pseudobullae” in specialized multituberculates see [144]).

## Significance of stem therian petrosals

### Phylogenetic Placement of the Höövör Petrosals

The close similarity between H1 and H2 in terms of size, morphology, and provenance supports the hypothesized sister-relationship [2] between the two taxa represented by these specimens (Fig 1). However, the above descriptions of the external anatomy of H2, and the internal features of both Höövör petrosals, also do not provide a decisive conclusion to the problem of the phylogenetic location of these specimens with respect to the mammalian taxa known from dental remains recovered from Höövör.

However, for the following reasons, the hypothesis that the Höövör petrosals represent both of the major stem therian lineages known from dental elements at this locality is still defensible as one of the two most probable hypotheses for the phylogenetic assignment of these specimens (the other, and equally probable, hypothesis is that they are both gobiconodontid taxa). First, irrespective of the relative relationship between the H1 and H2, the robust development of caudal and lateral structures on H1 (Fig 6a) strongly suggest its affinity as a gobiconodontid (as mentioned by [2]). The anatomical evidence supporting this assignment is derived from its elongate and robust paroccipital process (as also seen in *Repenomamus;* [145]), and the lateral deflection of the anterior lamina and the thickened area of bone caudal to it (termed the anterolateral recess by [2]). While these characteristic have not previously been scored and used in phylogenetic analyses, both of these features are likely associated with mediolaterally widened mandibular condyle seen in gobiconodontids and their nearest relatives (such as *Repenomamus).* Additionally, relative to H2, H1 shows a shorter rostrocaudal extent of the floor of the cavum supracochleare and the tympanic aperture of the prootic canal positioned more posterolaterally to the secondary facial foramen (Fig 2a), both features suggested as being characteristic of “triconodonts” [36,2,3]. While the clear presence of a cochlear aqueduct, processus recessus, and true fenestra cochleae are characteristics which have not been reported in prior descriptions of gobiconodontid cranial remains, they have been observed in gobiconodntids (GWR Pers Obs) and are possibly present in other members of the likely non-monophyletic group Eutriconodonta (e.g. the evidence for a cochlear aqueduct is inconclusive in the amphilestid taxon *Juchilestes* [146]). If the evidence supporting the gobiconodontid affinities of H1, and the possible trechnothere affinities of H2 is reliable, then based on the known roster of dental remains recovered from Höövör, the most likely taxonomic attribution for H1 would be to the fossil species *Gobiconodon borissiaki*.

While the similarities and suggested sister relationship between H1 and H2 have been remarked on here and by [2] (making it inconclusive as to which if either of these taxa are referable to *G.borissaiki)*, many of the advanced features of H1 suggest its more proximate relationship with trechnotherian mammals. These features are all detailed in the above descriptions, but include: the general reduction of venous proliferation within the pars cochlearis and loss of the hypopromontorial sinus (Fig 2b), 2) greater development of the vein of the cochlear aqueduct and its confluent sulcus within the scala tympani (Fig 12f), and 3) closer approximation of the foramen for the sacculoampullar branch of the vestibular nerve to the foramen for the cochlear nerve ventral to the extension of the crista transversa (Fig 14f and g). In addition to not displaying the above mentioned “gobiconodontid” characteristics seen in H1, H2 also does not display the apparently apomorphic lack of the fossa incudis on the petrosal, which is seen in H1 [2]. Even though the fossa incudis is not a preserved structure in H1, the separation of the fragmentary base of the mastoid region from its sutural surface for the squamosal demonstrates that these two regions would not have been in overlapping contact (as in H1 they are, causing the fossa incudis to be lost or relocated to the squamosal bone in this taxon) (Fig 6a). The advanced features seen in H2, combined with the gobiconodontid features seen in H1, support the closer phylogenetic affiliation of H2 with “symmetrodonan” mammals within the clade Trechnotheria. Under this interpretation the most likely dentally sampled taxon to which the H2 petrosal could be referred to is tinodontid genus *Gobiotheriodon* [147,148] however the rarity of these specimens makes specific assignment unreliable. The plesiomorphic trechnotherian status of H2 is also supported by the less laterally deflected margin of the tympanic surface (Fig 2a), suggesting that the glenoid fossa would have been located far posterolaterally, possibly distal to a pedicle-like concavity (“postglenoid depression”) on the posterior root of the zygomatic arch. This condition is seen in *Zhangheotherium* and *Maotherium* [38,50]. Unlike *Zhangheotherium*, however, the mediolateral extent of the medial margin of the pars cochlearis (the sulcus medialis, medial to the surface expression of the promontorium) is considerably wider in both H1 and H2.

While the new information on the Höövör petrosals provided here does not definitively resolve the phylogenetic location of these specimens, their status as stem therians, somewhat more closely related to crown therians than *Priacodon*, is secure. The following discussion on the functional implications of the morphological transition series provided by stem mammaliaformes, *Priacodon*, and more advanced stem therians (as represented by the Höövör specimens), only relies on this synecdoche.

### Osteological correlates of the Stria Vascularis

As can be sensed from the above synopsis, the current narrative of mammalian auditory evolution has been assembled predominantly from insights gained from mechanical and developmental sources (e.g. [95,91]). This is understandable given the expertise of most contributing authors, and the amenability of fossil material and preserved museum specimens to mechanical and embryological studies of the middle ear in particular (see [96]). However, this traditional emphasis has systematically neglected many of the most critical evolutionary developments known to occur within the inner ear of the stem therian lineage. Because most unique features of the therian auditory percept rely on apomorphic histological and physiological characteristics, such as the large and tortuous stria vascularis and the highly positive endocochlear potential, the relative silence on these aspects represent a deficit in our understanding [149].

One under-emphasized question that is apparent in the inner ear of all extant mammals is the conflicting set of demands placed on the auditory hair cells, which are required to alternately transmit and resist transduction currents at unparalleled rates while simultaneously being forced to function at a far remove from the vasculature providing for their nutrition, waste removal and oxygenation [28,62]. While the use of potassium as opposed to sodium is an adaptation for metabolic efficiency seen in the hair cells of all vertebrates [67, 70], the demands for high performance in the therian auditory endorgan seems to place this group’s auditory hair cells in a unique metabolic crisis. In response to this, the relatively large stria vascularis and uniquely formulated endolymphatic composition developed as a partial solution [150,151,18,12]. Because monotremes retain the ancestral endolymph-producing capillary plexus in Reissner’s membrane (see Fig 13b and f; [16,17]) in addition to the stria vascularis, and a greater number of radially oriented vessels crossing the cochlear duct [66], they seem to be less liable to metabolic distress than therians. However, the lamentable lack of physiological research on the cochlear apparatus of extant monotremes makes it impossible to judge whether the less specialized anatomy of the cochlear apparatus and supporting vasculature is attributable to weaker cochlear performance compared to therians, or because of some currently unrecognized requirement for increased cochlear vascularization developed in monotremes [71,62]. The exquisite sensitivity of the stria vascularis and endocochlear potential to induced hypoxia is an immediate and often exploitable feature in physiological studies of modern therians, however [70].

Because of the ease of misinterpreting bony anatomy in extinct animals at both anatomical and functional levels, caution must be used when attributing significance to any morphological novelty. However, when seen against the wider trend of reduction of the CVP in mammaliaformes generally, the localized venous hypertrophy at the sole intersection of the CVP and cochlear endocast strongly suggests an adaptive significance for the VCAQ at its earliest known instantiation in the Höövör petrosals described here. Because the stria vascularis is the most prominent organ within the abneural cochlear duct in both mammalian lineages bracketing the phylogenetic location of the Höövör specimens, the most reasonable functional attribution for this neomorphic vein is that it alleviated congestion within the hypertrophied stria vascularis in stem therians as it became the predominant or sole endolymph producing organ (Figs 9a-d and 13d). It is reasonable that this morphological innovation is itself a response is to the increasing demands on the stria vascularis for the production of the high endocochlear potential and potassium recycling biochemistry seen in extant therians [151, 62]. In extant therians, these functions of the stria vascularis make it one of the most metabolically demanding organs in the body, with specific requirements for arteriovenous and capillary ramifications within its parenchyma. The physiologically expensive nature of the stria vascularis is also reflected in extant mammals by its intricate and developmentally complex structure, which requires contribution from both cranial mesoderm (ectomesenchyme) andother specialized neural crest cell populations (e.g. the melanocyte-like strial intermediate cells), and in adults contains the only vascularized epithelial tissues known in mammalian anatomy [65]. While other vertebrate groups do show contributions from several embryonic tissue sources in the membranous lining of the endolymphatic labyrinth, nowhere outside of theria do these cell populations reach the level of organization seen in the therian stria vascularis [65,59]. It is therefore fortunate that the increasing demands of this unique organ would have required an enlarged and conspicuous vasculature detectable in the cochlear endocast of ancestral therians (Fig 12c-f).

The neomorphic appearance of the VCAQ in the Höövör petrosals may also reflect a more general reorganization of the vascular supply to the cochlear duct, allowing for the withdrawal of those blood vessels that adially span the unadhered membrane exposed toward the scala tympani and scala vestibuli sides of the cochlear duct (Fig 13d,e). Because the formation of the sulcus for the VCAQ seen in early trechnotheres represents the first bony integration of the vasculature of the cochlear duct within mammals, it is probable that the tissues of the membranous abneural wall of the cochlear duct relied to some extent on vasculature reaching it through its area of adhesion to the spiral ligament and bony cochlear canal (Fig 13 a-f). This is the situation seen in extant therians (e.g. [152]) where the VCAQ and other vessels do not cross the free basilar and vestibular surfaces of the cochlear duct (Fig 13e). This removal of blood vessels radially spanning the basilar and especially vestibular membranes is an apomorphic feature of the cochlear duct seen only in extant therians, and allows for the acoustic isolation of low-frequency interference generated by cardiovascular pulsations, away from the highly sensitive cochlear endorgan and supporting structures. As mentioned above, in extant monotremes and sauropsids the membranous tissues of the cochlear duct does not adhere to the bony side walls of the cochlear canal, leaving this structure mechanically unsupported and requiring that all blood vessels supplying the auditory epithelia travel in radial and longitudinal directions across the full length and circumference of the cochlear duct [66,24,17].

While the veins servicing the stria vascularis are the only soft-tissue component of the mammalian cochlear duct to leave a recognized osteological correlate, the high performance of the stria vascularis in modern therians is predicated on the presence and precise functioning of many unique molecular, cellular, and histological structures. The acquisition and localized hypertrophy of the vein of the cochlear aqueduct is therefore likely associated with the pre-existent or incipient presence of unfossilizable characteristics of the therian-style cochlear apparatus. Chief among these features is the reinforced compartmentalization of the endolymphatic and perilymphatic spaces, necessitated by the requirement to limit paracellular diffusion of potassium and calcium salts between these fluids (e.g. [153]). In extant therians, the stable attachment of the cochlear duct to the spiral ligament within the cochlear canal allows for the segregation and recycling of potassium and other ions through interconnected epithelial and connective tissue syncytia [150,153]. For therians, this fluid homeostasis is reliant on the efficient transfer of material through the slender and precariously located processes of root cells within the abneural cochlear duct [153], which would be liable to mechanical damage if left unsupported within the free membrane of the cochlear duct in monotremes. The single foramen and linear sulcus for the vein of the cochlear aqueduct within the cochlear canal endocasts of the Höövör petrosals, as opposed to a highly perforated and reticulating venous network, may also support the potassium recycling function of the stria vascularis and cochlear syncytia by allowing salts resorbed into the VCAQ an opportunity to diffuse back into the intrastrial space before being conducted into systemic venous circulation outside the otic capsule(e.g see [150]).

Even though nothing has been reported on the composition or electrical potential of endymph in extant monotremes, it has been observed that the effects of voltage changes in OHC membranes are not as dramatic as those seen in therian mammals, and that monotreme prestin has a peak non-linear capacitance voltage optimum which is far from the intracellular resting potential of OHCs [154,155]. The calcium concentration of monotreme endolymph is likely also much higher than the ~ 20 micromolar concentration seen in the cochlear duct of most therians, because of the continued existence of the lagenar otoconial mass in this group and the need to prevent uncontrolled dissolution of this mass [18].

### Osteological Correlates of Macromechanical tuning

The problem of transducing airborne sound into an electrical signal and its further decomposition it into spectral components, are a sophisticated functionality endowed to the cranium of many terrestrial vertebrates. As summarized in [15] the convergent innovations for airborne sound perception present in extant anurans, archosaurs, squamates, and mammals all operate with comparable levels of sensitivity and selectivity for frequencies under approximately 8 kHz. At the level of the auditory endorgan, the strategies for frequency selectivity (tuning) seen in these groups are effected through a homoplastic mixture of molecular, bony, and histological adaptations. The field of comparative hearing [156] therefore can valuably inform hypotheses of auditory capacities in extinct amniotes through the ancestral reconstruction of symplesiomorphic character states, or comparisons with taxa showing analogous adaptations [71,12]. As such, inferences about which of several non-mutually exclusive tuning mechanisms are present in early mammals are necessarily based on the distribution of the several forms of tuning in extant mammals and sauropsids [157, 105]. Because of this dependence on extant representatives, the almost complete lack of physiological studies on the living monotremes in particular imposes a major obstacle to our understanding of tuning mechanisms in the synapsid lineage [98,64]. However, the capabilities of extant therians to process frequencies well above 10 kHz emphasizes the necessity to synthesize available bodies of physiological research on auditory tuning and paleontological provenance in order to reconstruct a consistent scenario of the timing and ecological context of the evolution of this valuable sensory modality.

What is currently known about the distribution of tuning mechanisms across tetrapods suggests a major dichotomy in strategy between ancestral electrical tuning (described above) and several forms of mechanical tuning [105]. The varieties of mechanical tuning can be conceptually decomposed into intrinsic (action at the level of the hair cell) versus extrinsic (action at the organ level or larger), and active (requiring cellular energy) versus passive (based on inert geometrical and material properties) mechanisms (creating a total of four discrete catagories; [157]). All of these tuning mechanisms are characterized by some degree of tonotopy (the correlation of best frequency response with anatomical distance) within the auditory endorgan, and therefore a corresponding Space Constant (SC) expressed in units of millimeters per octave (an octave is a doubling of frequency). As mentioned above, the extant amniotes that rely solely on the plesiomorphic mechanism of electrical tuning *(Sphenodon* and the chelonians), show some of the smallest SC values (0.3 mm or smaller) because of the extremely short length of the auditory papilla. As such, these forms, and most likely all early amniotes, show only short and undifferentiated sacculocochlear recesses within their labyrinthine endocasts. Conversely, in all non-mammalian amniotes showing a differentiated middle ear a form of mechanical tuning is emphasized that relies on tonotopic variation in the mass, stiffness and number of stereovilli of the hair cells within the auditory papilla (homolog of the mammalian organ of Corti). This mechanism is instantiated in active (molecularly driven motion of stereovilli) and passive forms [105]. This category of tuning is dependent on organelle-level features present within the auditory hair cells themselves and so is termed micromechanical tuning; and it is associated with an intermediate range of SC values (less than 1 mm/octave up to several mm/octave; [15]). Because of the elongate but absolutely short cochlear canals seen in the earliest mammaliaformes (such as *Morganucodon;* [11,73,158], it has been hypothesized that micromechanical tuning provided the initial impetus for the development of the bony cochlear canal within the synapsid lineage as well [18]. However, in all known extant mammals (monotremes and therians) no evidence for significant electrical or micromechanical tuning has been physiologically recorded ([149]; although see [130] for possible evidence of active micromechanical tuning). Where characterized best in advanced therians, the sole form of tuning is based on an apomorphic extrinsic mechanism termed macromechanical tuning. Because this form of tuning is based on organ-level vertical dislocations of the cochlear duct and its contents, it is associated with SC values averaging 2.5 mm/octave in modern small therians [13]. This form of tuning is unlikely to have had an innate evolutionary advantage within the synapsid lineage until forms appeared with cochlear canals long enough to accommodate at least several octaves (somewhere near the emergence of Crown Mammalia; Fig 15).

Therefore, even though little is known about tuning modalities in monotremes, there are strong reasons to suspect that somewhere along the backbone of synapsid evolution, between earliest mammaliaformes and crown therians, a shift toward extrinsic macromechanical tuning and away from more plesiomorphic forms of tuning should be recognizable. The presumed reliance on macromechanical tuning in all extant mammals is likely also reflected by the universal presence of a true organ of Corti, with its functional differentiation of Inner and Outer Hair Cells (described above) and characteristic arrangement of membranes within the cochlear duct [136].

While the presence of macromechanical tuning appears to be consistently present within Mammalia, there is an obvious spectrum in terms of its performance and its morphological/molecular accommodation across mammalian species; with the monotremes defining the lower end and eutherians the higher end of the spectrum [12]. This spectrum is also recognizable in both the active and passive mechanisms of macromechanical tuning [105]. For instance, wide scale comparative studies on the structure of the prestin protein [154] estimate that monotreme prestins are much less capable of useful electromotility at physiological voltages. Additionally, monotremes show a lower proportion of Outer Hair Cells (expressing prestin on their basolateral surface) to Inner Hair Cells [136]. These observations support the generally comparable level of sensory traffic, but weaker instantiation of the macromechanical active process in the monotremes.

The macromechanical passive process is effected by the gradient in compliance of the basilar membrane, and its monotonic increase in width and decrease in depth as it runs apically beneath the organ of Corti [105,106]. Therefore, the passive process is present even in an inert and lifeless basilar membrane; and its capacity to tonotopically propagate traveling waves is modulated by its lateral attachments to the cochlear canal, or lack thereof [159,160]. The concerted action of this mechanical arrangement filters the range of frequencies presented to each individual auditory hair cell, allowing each cell to specialize for the transduction of a filtered frequency bandwidth [105]. Morphological features suggesting a weaker commitment to the macromechanical passive process in monotremes include their large and relatively untapered basilar membrane (which is wider than most of the largest and lowest frequency basilar membranes in found in extant therians; [136]), and lack of any adherence of the basilar membrane (or any other part of the endolymphatic cochlear duct) to rigid supports within the bony cochlear canal [17].

The stem therian labyrinthine endocasts described here contain osteological features which allow them to be placed along the spectrum of passive macromechanical adaptation defined by extant monotremes and therians. In particular, *Priacodon* and both Höövör petrosals differ from early mammaliaformes and monotremes, and resemble the more derived cladotherians, in the presence of a well-developed secondary bony lamina (Fig 14c,d). When present in modern therians, the secondary bony lamina provides a rigid attachment for the abneural cochlear duct and contributes to the concentration and transmission of tensile forces across the basilar membrane (Fig 14e; [22]). The first appearance of a well-formed secondary bony lamina in these stem therian endocasts (Figs 9 and 13c, d) is therefore strong evidence for the abneural adhesion of the cochlear duct with the bony cochlear canal, and may also signify the presence of the spiral ligament and its specialized populations of fibroblasts (e.g. tension fibroblasts; [161]). These soft-tissue specializations are known to be associated with the secondary lamina in extant therians and are critical for normal hearing function. However, as mentioned above, lack of an opposing primary bony lamina in *Priacodon* and the Höövör petrosals, or even an ossified tractus foraminosus, makes it unlikely that tensile forces could be transmitted and concentrated across the basilar membrane through its attachment along the secondary bony lamina in these forms. The stem therians described here therefore show an intermediate level of passive macromechanical adaptation by showing some rigid mechanical support for the cochlear duct, but lack the diametrically opposing bony laminae required for the suspension of the basilar membrane, which is associated with the tonotopic region of frequencies greater than 10 kHz in modern therians [14]. Macromechanical active and passive processes were likely also associated with each other [154]. As such, it is reasonable to hypothesize that taxa showing a bony secondary (or primary) lamina represent useful phylogenetic calibrations for the initiation of modern selective regimes responsible for the development of rapid electromotility in the prestin molecule [154], which is known to be much more weakly instantiated in modern monotremes [155].

Finally, the formation of a true round window and perilymphatic canal (aqueductus cochleae), such as that seen in both Höövör specimens (Figs 2a and 6a), does not have a clear functional interpretation but has been hypothesized as improving the vibrational insulation of the inner ear [136]. Conversely, the first appearance of these features and the development of a process recessus may have been initiated as a structural byproduct of increasing braincase width and associated lateral displacement of the perilymphatic foramen relative to the jugular foramen.

This comparative and fossil evidence for micro-and macromechanical tuning in the earliest crown mammals, and especially along the therian stem, strongly suggests that a cochlear space constant within the range of values associated with macromechanical tuning in extant mammals would be applicable to these fossil members of crown Mammalia as well [13]. As mentioned above, these values range from ~1 mm/octave minimally for monotremes and small therians, to an average of 2.5 mm/octave, and extending to increasingly higher values in large-bodied species [64]. Therefore, given the ~4 mm length of the cochlear duct in the stem therian endocasts described here, a range of four octaves maximally seems reasonable as an estimate for the frequency bandwidth available to these forms. The relationship between the interval of detectable frequencies and upper detectable frequency is, however, also dependent on the lowest detectable frequency. Given a plausable lower frequency limit for small amniotes of 500 Hz (many modern lizards and birds have even lower limits, ~ 50-100 Hz), this would place the upper frequency limit for these stem therians at ~ 8 kHz; and if given lower frequency limit of 100 Hz typical of plesiomorphic amniotes the corresponding upper limit would be much lower at ~ 3.2 kHz [24]. Conversely, an exorbitantly high low-frequency limit of 1 kHz, as is seen in modern *Mus* and other high-frequency specialists, would correspond to an upper frequency limit of 16 kHz, similar to the upper limit seen in modern *Ornithorhynchus* [162,16]. This generous estimate of a low-frequency limit is likely unrealistic, given that it requires the abandonment of the amniotic ancestral range of 1 kHz and lower frequencies in morphologically plesiomorphic amniotes. The probable retention of a lagenar macula in *Priacodon* also diminishes the frequency band available to this taxon as well, by taking up ~ 1 mm of length of the cochlear canal (Fig 12h; [68]). The most realistic range of upper frequency limit values in *Priacodon* and the Höövör specimens is therefore likely between ~ 5-10 kHz. However, it is important to emphasize that even under the extreme assumption that macromechanical tuning was entirely absent in early mammals, and therefore space constants within the macromechanical range would not be applicable to the stem therians described here, no form of micromechanical or electrical tuning would feasibly have allowed the upper frequency limit to extend above approximately 10 kHz either [15,12]. Despite the existence of several autapomorphic high-frequency non-mammalian tetrapods [163,164], no combination of the phylogenetically typical forms of auditory tuning would have allowed the stem therians described here to detect ultrasonic or even near ultrasonic frequencies (~10 kHz or higher; [15]).

### Osteological correlates of the Lagenar Endorgan (or lack thereof)

The hair cells comprising the lagenar macula are perhaps the most variable and least understood of the many epithelial cell types found around the heterogenous endolymphatic lining of the pars inferior [98]. This is especially so in the case of extant mammals, where in the monotreme lineage this sensory epithelium is located between the scala medial and scala vestibuli within a specialized lagenar sac [17]; while extant therians are unanimous in their lack of any morphological expression of the lagena altogether. Still, several inducible genetic atavisms seen in rodent models suggest that the distal extent of the therian organ of Corti persists as the syngenetic homolog of the lagenar macula [98,80].

Nonetheless, as a discrete organ, supporting an otolithic mass and recruiting the innervation of a dedicated lagenar nerve and ganglion, the absence of the lagena is one of the unique and important features of the therian inner ear. It has also been suggested that the loss of the lagenar macula acted as a proximal cause, or immediate correlate, of the development of several other synapomorphic features of therian cochlear physiology; such as the extremely low calcium concentration and high electrical potential of therian endolymph [62,18]. In sauropsids, and presumably also the earliest synapsid taxa, a high (several hundred micromolar up to 1 millimolar; [18]) calcium concentration of the cochlear fluids is required to support electrical tuning and protect the lagenar otolithic mass. A minimal ambient calcium concentration is also known to be required to prevent dissolution of vestibular gravistatic structures [165] which (where examined in the otoconia of modern rodents; [166]) show compositional turnover on a monthly timescale. The released obligation to generate high calcium endolymph within the cochlear duct likely allowed stem therians the adaptational leeway to reformulate several aspects of their cochlear biochemistry, resulting in the low (~20 micromolar) calcium concentration in cochlear endolymph, and the correspondingly sharp calcium gradient along the endolymphatic labyrinth generally [167,168]. There is also some evidence that the capillary plexus in Reissner’s membrane is specifically associated with the support of the lagenar otolithic mass because of the retention of localized Calcium ATPases to this membrane in extant rodent models [151]. The increasingly exclusive reliance on the mammalian stria vascularis for endolymph production occurring during the evolution of crown therian mammals may therefore be directly correlated with the lost capacity to support a functional lagenar endorgan [18].

The timing and functional significance of lagenar loss is complicated by the lack of pertinent comparative and paleontological evidence, however. In particular, the lack of experimental recordings of the normal functioning of the lagenar endorgan in sauropsids and monotremes, and conflicting hypotheses regarding the relative importance of its gravistatic versus auditory modalities [24,69,98], make the physiological preconditions of lagenar loss in stem therians difficult to interpret. The available fossil material is also ambiguous because of the loose osteological association seen between the lagena and the surrounding skeleton of the otic capsule. Bony structures such as an apical inflation of the cochlear canal, and sulci or canals for the lagenar nerve, are variably present among mammaliaformes; however, based on the wider distribution of the lagena among amniotes, a functional lagenar macula almost definitely existed in synapsid taxa lacking such osteological correlates, both preceding the early mammaliaformes, and succeeding them within the mammalian crown group. Therefore, absence of evidence for a functional lagenar endorgan is not evidence of its absence, and it is likely that early stem therians such as *Priacodon* that lack obvious osteological correlates of a lagenar macula retained it nonetheless; both because of the presence of lagenar correlates in multituberculates and other theriiform taxa diverging later in stem therian phylogeny (Fig 15; [87,56], and the lack of bony features indicating a switch to the modern therian-style of cochlear physiology predicated on the lack of the lagenar macula [36,69,18]. The reference to the loss of the lagena as “the Cretaceous Cochlear Revolution” [18] may therefore be misleading in that it is currently unclear as to whether the morphological expression of the lagena was lost in an evolutionarily punctuated event or a gradual interval of decreasing usefulness. Based on the stratigraphic distribution of fossil therians, and most phylogenetic hypotheses of their relationships (i.e [33,169] inter alios), the loss of the lagena certainly occurred within the Jurassic if not earlier, and possibly several times.

The internal bony anatomy visible in the high-resolution scans of the Höövör petrosals provide the earliest indications of advanced features seen today only in therian mammals. As outlined above, several of the osteological features seen in these specimens suggest that the cochlear duct achieved at least some adhesion to the abneural margin of the cochlear canal (Fig 14c-e) and show increased (if not exclusive) reliance on the functioning of the stria vascularis as an endolymph producing organ [70]. These characteristics are unique to therian mammals among extant vertebrates, and (combined with the presence of a straight and distally tapering cochlear canal in the Höövör cochlear endocasts; Fig 12e,f) strongly suggest that these stem therians have greatly reduced or lost the lagenar macula altogether. These forms would therefore have attained other soft tissue characteristics seen in modern therians such as a terminal helicotrema [17].

The consistent presence of dorsoventral coiling to accommodate the lengthened cochlear canal seen in later cladotherian mammals may have then been enabled by the lagenar macula reorienting to a position where its sensory input was mostly or exclusively responsive to vertical linear acceleration [16], and therefore completely redundant to stimulus from the saccule and utricle [18]. The complete loss of the remnant hypopromontorial sinus in therians may also be an effect of dorsoventral expansion of the cochlea within the pars cochlearis.

The high likelihood that the Höövör petrosals described here were beneficiaries of the “cochlear revolution” provides a useful perspective on the selective regime responsible for the loss of the lagena and the development of the unique therian cochlear physiology. Particularly, because of criteria outlined in the discussion of macromechanical tuning, there was almost no capacity for high-frequency (above ~ 10 kHz) hearing in these early stem therians. Therefore, whatever factors led to lagenar loss must not be related to the development of ultrasonic hearing capacities. This is complementary to hypotheses that stem therians relied to a substantial extent on substrate based (i.e. non-tympanic) sound conduction [170,56]; more importantly, this evidence is contradictory to the hypothesis that these unique features of the therian inner ear are related to the unique capacity of extant therians to detect and localize ultrasonic sound sources [11,64]. As with the innovation of the three ossicle middle ear found in mammaliamorphs, the unique inner ear mechanism developed in ancestral stem therians (advanced theriimorphans or early trechnotheres) developed in service of acoustic performance within an ancestral frequency range, and later proved capable of being exapted into ever higher frequencies in descendant therian taxa. The late Cretaceous meridiolestidan *Coloniatherium* [4], with a fully coiled cochlea and a centrally located modiolus, optimizes in most phylogenies as an independent acquisition of these features. A highly derived inner ear morphology (with complete coiling, and without a lagenar inflation) is also present in more plesiomorphic taxa like *Cronopio* [171] and terminal taxa such as *Peligrotherium* [140] and *Necrolestes* [139,7]. The detailed similarities between meridiolestidan and therian inner ears, with their potential auditory convergences, have yet to be fully explored.

### Auditory Localization in Stem Therians

Aside from the difficulties related to the sustenance of greater mass-specific caloric requirements, within topographically more heterogenous habitats, small mammals are faced with novel challenges for the segregation and localization of sound sources. This “small mammal problem” has been a central focus in the traditional explanation for the advent of ultrasonic hearing in therian mammals, because of the requisite use of high frequencies for sound localization at small body sizes (e.g. [136]). Indeed, the abilities of even very “primitive” small mammalian insectivores to utilize broadband and high-frequency cues for auditory localization, even to the point of echolocation in many cases, is well recorded (e.g. [172]). Therefore, it is probable that ultrasonic capacities evolved in the earliest crown therians in response to selection for greater localization capabilities; however, the evidence reported here of the likely very low upper detectable frequency limits in stem therians, and the manifest capacity of small-bodied sauropsids to locate sound sources with frequencies below 5 kHz, suggest that traditional narratives of the evolution of ultrasonic hearing require qualification [1, 31,123,64].

The one-dimension waveform presented to each ear carries very limited, and difficultly extracted, information on the spatial location of its source. These difficulties in localization are both physical and computational in nature. For instance, the physical coupling of sound frequencies in air ranging from 20 Hz – 20 kHz to corresponding wavelengths ranging from 17 m – 17 mm (respectively), require sensitivity to frequencies with corresponding wavelengths larger than the head size (interaural distance) of many low-frequency limited terrestrial vertebrates. The many sauropsid groups to have developed a tympanic middle ear have convergently solved this physical challenge by the coupling of both right and left tympanic membranes through the medial air mass comprising their interaural canal (I.e the cavum tympani and other contiguous cavities). This has the effect of increasing interaural delay times and allows the detection of interaural phase differences at the level of the auditory transducers themselves, alleviating the neurological requirement to develop a complex internal representation of the external environment within the central nervous system [17,121, 122]. This “pressure-gradient receiver” form of auditory localization is the only mechanism for the perception of low-frequency limited sound sources known in terrestrial vertebrates, and because of its wide and homoplastic distribution in sauropsids, it possibly also evolved in early eucynodonts and later synapsid taxa with angular tympanic membranes [108].

Conversely, modern therian mammals have developed a predominantly computational strategy for the localization of sound sources, based on the isolated functioning of both ears as simultaneous and independent “pressure receivers”. The adaptations providing this capacity are categorized into monaural and binaural mechanisms, each making specific minimal demands for broadband and high-frequency (near-ultrasonic) hearing. They are also) commonly specialized for vertical and azimuthal localization, respectively. All of these mechanisms also make substantial demands on the central nervous system, such as the required detection of binaural coincidence on submillisecond timescales, and novel processing in the lower auditory pathways [174, 123]. Because of the conflicting demands that the pressure-gradient receiver versus pressure receiver forms of auditory localization j place on the specific structure of the external, middle, and inner ear, the synapsid lineage must have reduced and ultimately lost its pressure gradient receiver capacities before the advent of the modern form of therian sound localization (if pressure gradient receivers existed in early synapsid taxa at all). As described in the synopsis of synapsid evolution above, the progressive constriction of the cynodont interaural passage [110] into a facultatively patent) eustachian tube may represent the progressive reduction of the hypothetical pressure-gradient receiver and the concomitant loss of the ancestral capacity for sound localization; as attenuations of greater than 15 dB by the interaural channel would effectively acoustically decouple both ears [1]. The sequential development of the MME, PME, DME, and development of discrete bullae in the therian crown group may therefore represent the progressive reduction and eventual loss of the ancestral mechanisms for sound localization. Likewise, the presence of a broadly open interaural canal in some extant therians [175] which could plausibly be interpreted as atavisms to a pre-mammalian condition. The platypus *Ornithorhynchus* [51] also shows a patent interaural communication which in fact narrows substantially before merging with the proximal segment of the pharynx. Depending on how the primitive condition for the last common ancestor of amniotes is reconstructed the condition of *Ornithorhynchus* could be primitive, or that of the Echidnas with a recognizable Eustachian tube would be symplesiomorphic for Mammalia. At present we consider this question unresolved and the primitive mammalian condition equivocal.

#### Binaural sound localization

Both the crown mammalian common ancestor and the stem therian taxa described here most likely attained a PME state of the middle ear [56], and therefore would have attenuated or lost the tentatively ancestral pressure-gradient capacity for sound localization if indeed it had previously existed. However, regardless of whether or not the pressure-gradient receiver mechanism operated in earlier members of the synapsid lineage, the functional limitations imposed by the short length and macromechanical adaptations seen in the stem therian cochlear canals described above would have also precluded the useful functioning of the modern sound localization strategies seen in extant therians as well. The two mechanisms seen in extant therians most useful for azimuthal localization are based on the comparison of stereo binaural input, and are termed the Interaural Time Difference (ITD) and Interaural Level Difference (ILD) mechanisms [1]. These two mechanisms are based on the capacity to contrast the time of arrival of distinctive spectral features (ITD) or the instantaneous amplitude of the stimulus (ILD), respectively. The ITD mechanism is further subdivided into two subtypes, with both being most accurate when the sound source is composed of frequencies low enough (lower than ~4 kHz) to preserve the temporal periodicity of the stimulus within its neural encoding (i.e. the “phase-locking” mechanism; [176]). However, the ability to precisely contrast arrival times of spectral features is contingent on a sufficiently large binaural time delay, which itself is a function of the Functional Head Size (FHS a metric of interaural distance measured in microseconds; [31]) of an animal. Therefore, ITD based localization is emphasized to the exclusion of ILD in modern therians with large FHS values (e.g. domesticated ungulates). Conversely, modern small mammals (with smaller than 200 microseconds FHS) emphasize ILD, and small mammals using only one binaural cue use ILD. The known modern small therians that use both ITD and ILD are low-frequency specialist rodents [31,177,95]. As such, even with a possibly convoluted interaural canal extending the binaural time delays somewhat [120], the estimated FHS of ~ 50 microseconds or less in the crown mammalian common ancestor [178] would place the earliest mammals within the predominantly ILD size range. This inference is also supported by the purportedly more plesiomorphic construction of the ILD neural circuitry within the lateral surperior olive and related mammalian auditory brain-stem features [174].

However, while it is very unlikely that the first stem therians were able to use ITD as a localizing mechanism, the low estimated upper frequency limit of the stem therian cochlear endocasts presented here also suggests that the ILD mechanism would also have been inoperative in these animals as well. While the exact frequency requirements for ILD functioning are somewhat variable across extant therian species, because the attenuation of sound amplitude is produced by cranial “shadowing” of the stimulus as it propagates across the tissues of the head, ILD requires frequencies with corresponding wavelengths shorter than the interaural distance of that particular taxon [179]. In the case of the small stem therians presented here, and with a very generous estimate of interaural distance estimate of ~1 inch, the corresponding minimal ILD frequencies of ~13 kHz would be likely be just marginally within or above the upper frequency limits of these small taxa. However, while it is likely that the lowest frequencies required for ILD were perceptible by the stem therians described here, the proper functioning of the ILD mechanism (and the other localization mechanisms) in extant small therians relies on the availability of a wide band of frequencies beginning with the lowest useful frequency. In the case of stem therians, the ILD mechanism would therefore have been poorly functional relative to its performance in modern therians if it had been present at all [123]. This is also complementary with what has been suggested as the most likely evolutionary trajectory for the assembly of the modern neural circuitry supporting ILD, where the hypothesized incipient stages of binaural processing was likely only sufficient for the segregation of discrete simultaneous sound sources, and possibly their relative localization [174].

#### Monaural Sound Localization

Therefore, while the capacity for sound localization based on the physical interconnection of both ears, or the simultaneous comparison of the electronically encoded input from both ears, would be inoperative or poorly functional at best in stem therians, the final method of auditory localization known in extant tetrapods does actually have some empirical support from the fossil record of Mesozoic mammals. This final form of auditory localization is based on the spectral alteration of monaural input (termed Head-Related Transfer Functions; [174]) by the presence of a specialized external pinna. In extant therians this pinna-based form of auditory localization is most important in front-back discrimination of sound sources, and in specialized species allows the vertical localization of sound sources, such as along a mid-sagittal plane. Where tested in llamas frequencies, ~ 3 kHz and higher allow for consistent resolution of the front versus back location of a sound source [31]. In cats tested with frequencies greater than ~ 10 kHz, the vertical location of sounds is resolvable, as is the precise location of a sound source within each ear’s “cone of confusion” [180]. As summarized above, the anatomical evidence of an involuted pinna in *Tachyglossus* [134] and the excellent soft tissue preservation in the eutriconodont *Spinolestes*, suggest that an elaboration of the external ear may have been present in crown mammals (and was very likely present in early theriimorphs; [135]). Because the earliest crown mammals may have faced a situation where the use of the pressure-gradient receiver was becoming untenable and the use of binaural localization was still unfeasible at both a physical and neurological level, the mammalian pinna may have initially appeared as an inefficient but sufficient method to monaurally localize sound sources, possibly only for front-back discrimination (there is also a significant amplification effect provided by the pinna as well; [20]). The more sophisticated capacities for monaural vertical localization, predicated on minimal frequencies higher than approximately 10 kHz, likely only developed in crown therians or their immediate ancestors as near-ultrasonic frequencies became detectable within the therian lineage. While the observation, provided in [12], that the dimensions of the soft tissue impression of the external pinna in *Spinolestes* would most efficiently provide localizing information (such as spectral cues) at frequencies above 20 kHz, for the reasons outlined above is appears unlikely that these types of advanced localization mechanisms were present in gobiconodontids such as *Spinolestes* and (possibly) the Höövör specimens.

This hypothesis regarding the acoustic capacities in Jurassic and Early Cretaceous stem therians, synthesized from both paleontological and physiological evidence, presents an unimpressive picture of the ancestors of modern therian mammals as poorly equipped nocturnal insectivores compared to modern standards. While many extant small mammals show sophisticated sound localization capacities, even with frequencies below 20 kHz, the reasoning outlined above suggests that stem therians would not have attained a sufficient octaval range of frequencies, a high enough upper frequency limit (or both), to have been able to usefully localize short duration sounds faster than visual localization alone [31]. This may reflect the less competitive nature of the small insectivore niche within Mesozoic terrestrial ecosystems, but could equally support the presence of non-tympanic forms of conduction such as the hypothesized direct conduction of sound through Meckel’s element as suggested by [56]. The combination of poor auditory localization abilities with adaptations suggesting increased sensitivity and selectivity at low frequencies in stem therians may therefore be a compromise between the low-frequency requirements for seismic sound conduction and the increasingly specialized capabilities for airborne sound localization.

It is currently ambiguous what behavioral and autecological implications the transitional morphology of these stem therian petrosal has for the reconstruction of Mesozoic mammals. However, the inference of poor sound localization capabilities within the advanced stem therians is not based on any single ancestral reconstruction of the cochlear tuning mechanism or auditory physiology within the crown mammalian ancestor or early stem therians. All known cochlear tuning mechanisms typical of modern amniotes, working individually or in combination, would be incapable of extending the upper frequency limit in the fossil taxa described here to frequencies above ~ 10 kHz. If, as seems most likely, macromechanical tuning was present within the theriimorphans and trechnotheres, the large concomitant SC values associated with this type of tuning would prevent the short cochleae in these forms from extending into frequency ranges much higher than 10 kHz. Even if a more ancient form of intrinsic tuning (electrical or micromechanical) was present in the earliest stem therian mammals, the low upper frequency limits associated with these forms of tuning in modern tetrapods would also limit the maximal detectable frequencies in stem therians to under ~ 10 kHz (exceptions to this frequency limit are seen only in very specialized and phylogenetically restricted taxa among sauropsids and amphibians [163,164].

## Conclusions

The descriptions and discussion provided here highlight the phylogenetically heterogenous nature of stem therian petrosal evolution throughout the Mesozoic. The approximately 50 million year duration separating the stratigraphic proveniance of the Upper Jurassic triconodontid and Lower Cretaceous Höövör specimens focused on here represent a wider temporal interval of morphological aggregation during which several derived internal labyrinthine features make their first appearance within the backbone of therian evolution. Even before the advent of the dorsoventral cochlear coiling characteristic of modern crown therians and their cladotherian relatives, both *Priacodon* and the Höövör petrosals demonstrate morphologies suggestive of greater acoustic performance unique to this lineage within Mammalia.

Osteologically, these specimens demonstrate that several of the internal labyrinthine features appearing in the earliest mammaliaformes (e.g. lateral curvature of the cochlear endocast and lagenar inflation) were lost before most of the advanced cladotherian morphological features related to cochlear coiling and the bony support of the cochlear nerve appeared (Fig 9). Interestingly, this evolutionary loss of lateral curvature of the cochlear canal before the advent of its dorsoventral coiling is not matched by the developmental trajectory of the membranous cochlear duct; as seen in rodent models [181,182,91], where lateral curvature and subsequent dorsoventral coiling form two discrete stages in the development of the endolymphatic labyrinth [183].

For reasons outlined above, it also seems likely that the perceptual capacity for sound source localization was undeveloped or reduced in early mammaliaformes. The hypothetical pressure gradient receiver form of auditory localization was either reduced or absent for an extended interval before the capacity to use advanced ILD and ITD forms of localization developed in the immediate ancestors of crown therians. Because of their attainment of a PME, short cochlear canal, and small head size, it appears that the stem therians described here represent taxa that existed in this phylogenetic interval (paraphyletic group) of forms with weak sound-source localization capabilities. The one possibly operative localization mechanism in these forms would have been based on monaural pinna-based signals, for which the preservation of an external pinna in one exceptionally preserved theriimorph specimen can be considered evidence [135]. However, even using these pinna-based cues, this form of localization in the stem therians described here would mostly be competent for front-back localization of sound sources.

This is not to suggest that early stem therians displayed poor hearing capacities generally, and the presence of a secondary bony lamina within the cochlear endocasts described here suggests some amount of adhesion between the cochlear duct and spiral ligament with the abneural cochlear canal (Fig 14c,d). This is an advanced level of structural organization beyond the state seen in even modern monotremes, and is likely associated with a greater commitment to macromechanical tuning than that seen in extant monotremes. The attachment of the basilar membrane to the newly evolved secondary lamina before the advent of the primary bony lamina also suggests that the basilar membrane in these forms was less tense and stabilized than is typical of modern therians. It is also likely that the low-frequency limitations of the stem therians described here would not have precluded these forms from relying on an insect-based diet; which has been predicted as the mainstay of most generalized Mesozoic lineages including those represented by the fossils described here [184-188]. One study [189] estimates that at least one Middle Jurassic katydid species produced frequencies (~6.4 kHz) which would very plausibly be detectable by these stem therians.

This report details the bony features pertinent for the phylogenetic and soft tissue reconstruction of *Priacodon*, Höövör petrosal 1 and especially the newly described Höövör petrosal 2 (Figs 1–9). However, perhaps the most significant aspect of the morphology presented by these specimens is what they entail regarding the rate of auditory evolution near the therian crown group. If the uniquely derived and phylogenetically unstable clades Gondwannatheria and Multituberculata are excluded, the triconodontid and symmetrodont specimens used here provide a phylogenetic bracket around the advanced clade Cladotheria. To date the majority of previously described Mesozoic petrosal specimens belong to Cladotheria, and several convergent derivations of the fully coiled cochlear canal, tractus foraminosus (convergent with monotremes; Fig 14b,e), and primary bony lamina can be observed within this group. The petrosals described here corroborate the slow rate of upper frequency limit increase in the synapsid lineage up to the advent of Cladotheria; and that cladotheres may therefore be thought of as an evolutionary radiation into a high-frequency world [190]. Within the Cretaceous, both crown therians and South American dryolestoids both achieve a structurally “modern” fully-coiled form of the cochlear canal [72,135,4]. This may have been a response to selection for sound-source localization, particularly the capacity to locate brief environmental cues faster than visual inspection alone [1,31]. However, our hypothesis that this capacity was lacking, or poorly developed, in the immediate stem therian ancestors of the cladotheres suggests that either an appropriate selective pressure was absent in earlier Mesozoic environments, or that these Mesozoic groups were forced into evolutionary compromises between the high-frequency requirements for sound localization and possible behavioural requirements for low-frequency perception (such as non-tympanic sound conduction). Whatever the original impetus for the development of ultrasonic frequency sensitivity, the segregation of terrestrial vertebrate faunas into a high-frequency therian component and low-frequency sauropsid component has persisted from the Cretaceous to the present day.

## Final remarks on the value of fossil research in neurobiology

The fossil evidence summarized in this report provides a valuable constraint on estimates of frequency sensitivity in ancestral mammals produced in the neurobiological literature. These reconstructions, based on the geometrical (e.g. ideal transformer model; [89,95] and/or material properties of the middle ear and empirical correlations among extant model organisms [191], have typically produced unrealistically high upper frequency estimates. One particularly influential instance of this is the statement in [192]: “the results show that high frequency hearing [above 32 kHz] is a characteristic unique to mammals and, among members of this class one which is commonplace and primitive”. Based on the morphology of the early crown mammalian fossils described here, and the auditory performance of living monotremes [16], this statement is obviously false (see also [71] for other criticism of this paper). A later and more paleontologically informed report by [11] provide a similarly high estimated upper frequency limit, extrapolated from the observed negative correlation between length of the basilar membrane and upper detectable frequency in a sample of extant therians.

While the low high-frequency limitations of early mammals suggested by the fossil record and comparative hearing research are suggestive, more basic research (including investigations of the possible presence and strength of the endolymphatic potential in modern monotremes; [18]) is required before inferences on the performance and phylogenetic context of complex features of therian cochlear physiology (such as the endolymphatic potential) in ancestral mammals can be usefully understood. The evolution and development of complex abneural structures within the mammalian cochlea is however an excellent field of inquiry which stands to benefit from paleontological, neurobiological and molecular input. This research trajectory has a unique potential to return societal dividends through the illumination of several of the most common causes of age-related hearing loss. Diseases such as metabolic presbycussis [193], are a likely byproduct of the relatively recent evolutionary construction of the stria vascularis, and the evolutionary propensity of more recently acquired traits to be particularly liable to pathology [194].

## Acknowledgments

The authors would like to thank Michael Novacek, Curator of Paleontology at the American Museum of Natural History, for access to the Höövör specimens. Additionally, TH is grateful to Timothy Phelps at the Johns Hopkins Art as Applied to Medicine Program for his guidance in the production of the line art figures used in this report; Simone Hoffmann (NYIT College of Osteopathic Medicine); Julia Schultz (Universität Bonn); and Brian Davis and Nobuyuki Kuwabara (University of Louisville) for their insight and assistance in understanding the complex literature on the anatomy and physiology of the inner ear.

This work was performed in part at the Duke University Shared Materials Instrumentation Facility (SMIF), a member of the North Carolina Research Triangle Nanotechnology Network (RTNN), which is supported by the National Science Foundation (Grant ECCS-1542015) as part of the National Nanotechnology Coordinated Infrastructure (NNCI). The CT lab, Austin Texas produced images of some specimens and provided technical and instructional support to GWR. Specimen collecting and data analysis was funded in part by PICT-2016-3682 from CONICET-Agencia de Promoción Cientifíca y Técnica, Argentina; and NSF grants DEB 0946430, and DEB 1068089 to GWR and the Department of Anatomical Sciences, University of Louisville.

